# Deep Phenotyping and Lifetime Trajectories Reveal Limited Effects of Longevity Regulators on the Aging Process in C57BL/6J Mice

**DOI:** 10.1101/2022.03.25.485824

**Authors:** Kan Xie, Helmut Fuchs, Enzo Scifo, Dan Liu, Ahmad Aziz, Juan Antonio Aguilar-Pimentel, Oana Veronica Amarie, Lore Becker, Patricia da Silva-Buttkus, Julia Calzada-Wack, Yi-Li Cho, Yushuang Deng, A. Cole Edwards, Lillian Garrett, Christina Georgopoulou, Raffaele Gerlini, Sabine M. Hölter, Tanja Klein-Rodewald, Michael Kramer, Stefanie Leuchtenberger, Dimitra Lountzi, Phillip Mayer-Kuckuk, Lena L. Nover, Manuela A. Oestereicher, Clemens Overkott, Brandon L. Pearson, Birgit Rathkolb, Jan Rozman, Jenny Russ, Kristina Schaaf, Nadine Spielmann, Adrián Sanz-Moreno, Claudia Stoeger, Irina Treise, Daniele Bano, Dirk H. Busch, Jochen Graw, Martin Klingenspor, Thomas Klopstock, Beverly A. Mock, Paolo Salomoni, Carsten Schmidt-Weber, Marco Weiergräber, Eckhard Wolf, Wolfgang Wurst, Valérie Gailus-Durner, Monique M.B. Breteler, Martin Hrabě de Angelis, Dan Ehninger

## Abstract

Current concepts regarding the biology of aging are based on studies aimed at identifying factors regulating natural lifespan. However, lifespan as a sole proxy measure for aging can be of limited value because it may be restricted by specific sets of pathologies, rather than by general physiological decline. Here, we employed large-scale phenotyping to analyze hundreds of phenotypes and thousands of molecular markers across tissues and organ systems in a single study of aging male C57BL/6J mice. For each phenotype, we established lifetime profiles to determine when age-dependent phenotypic change is first detectable relative to the young adult baseline. We examined central genetic and environmental lifespan regulators (putative anti-aging interventions, PAAIs; the following PAAIs were examined: mTOR loss-of-function, loss-of-function in growth hormone signaling, dietary restriction) for a possible countering of the signs and symptoms of aging. Importantly, in our study design, we included young treated groups of animals, subjected to PAAIs prior to the onset of detectable age-dependent phenotypic change. In parallel to our studies in mice, we assessed genetic variants for their effects on age-sensitive phenotypes in humans. We observed that, surprisingly, many PAAI effects influenced phenotypes long before the onset of detectable age-dependent changes, rather than altering the rate at which these phenotypes developed with age. Accordingly, this subset of PAAI effects does not reflect a targeting of age-dependent phenotypic change. Overall, our findings suggest that comprehensive phenotyping, including the controls built in our study, is critical for the investigation of PAAIs as it facilitates the proper interpretation of the mechanistic mode by which PAAIs influence biological aging.

**Highlights:**

- Phenotyping at scale defines lifetime trajectories of age-dependent changes in C57BL/6J mice
- Central genetic and environmental lifespan regulators (putative anti-aging interventions; PAAIs) influence age-sensitive phenotypes (ASPs) often long before the appearance of age-dependent changes in these ASPs
- Corresponding genetic variants in humans also have age-independent effects
- Many PAAI effects shift the baseline of ASPs rather than slowing their rate of change

## Introduction

A large body of work, carried out over the past decades in a range of model organisms including yeast, worms, flies and mice, has identified hundreds of genetic variants as well as numerous dietary factors, pharmacological treatments and other environmental variables that can increase the length of life in animals ^1–3^. Current concepts regarding the biology of aging ^4^ are in large part based on results from these lifespan studies. Much fewer data, however, are available to address the question of whether these factors, besides extending lifespan, in fact also slow aging, particularly in the context of mammalian models.

It is important to distinguish lifespan vs. aging because it is well known that lifespan can be restricted by specific sets of pathologies associated with old age, rather than being directly limited by a general decline in physiological systems. In various rodent species, for instance, the natural end of life is frequently due to the development of lethal neoplastic disorders: Cancers have been shown to account for ca. 70 – 90% of natural age-related deaths in a range of mouse strains ^5–10^. Accordingly, there is a strong need to study aging more directly, rather than to rely on lifespan as the sole proxy measure for aging.

‘Aging’ is used as a term to lump together the processes that transform young adult individuals (i.e., individuals that have attained full growth and maturity) into aged ones with functional changes across multiple physiological systems, elevated risk for multiple age-related diseases, and high mortality rates ^3, 11, 12^. It is associated with the accumulation of a large number of phenotypic changes, spanning across various levels of biological complexity (molecular, cellular, tissue and organismal level) and affecting virtually all tissues and organ systems ^13, 14^. Aging can hence be approached analytically by assessing age-dependent phenotypic change, from young adulthood into old age, across a large number of age-sensitive traits covering multiple tissues, organ systems and levels of biological complexity ^15, 16^.

Deep phenotyping represents a powerful approach to capture a wide range of aging-associated phenotypic changes, since it takes into account alterations at molecular, cellular, physiological and pathological levels of analysis, thereby providing a very fine-grained view of the consequences of aging as they develop across tissues and organs ^10, 15–17^. The approach is therefore ideally suited to assess genetic variants, pathways, dietary or pharmacological factors previously linked to lifespan extension and, potentially, delayed aging. Deep phenotyping examines hundreds of parameters, many of which are expected to differ between young and old animals (hereafter called age-sensitive phenotypes; ASPs); these can be collectively used to address if and how a given intervention interacts with the biological processes underlying the signs and symptoms of aging (Fig. 1a).

**Figure 1:**
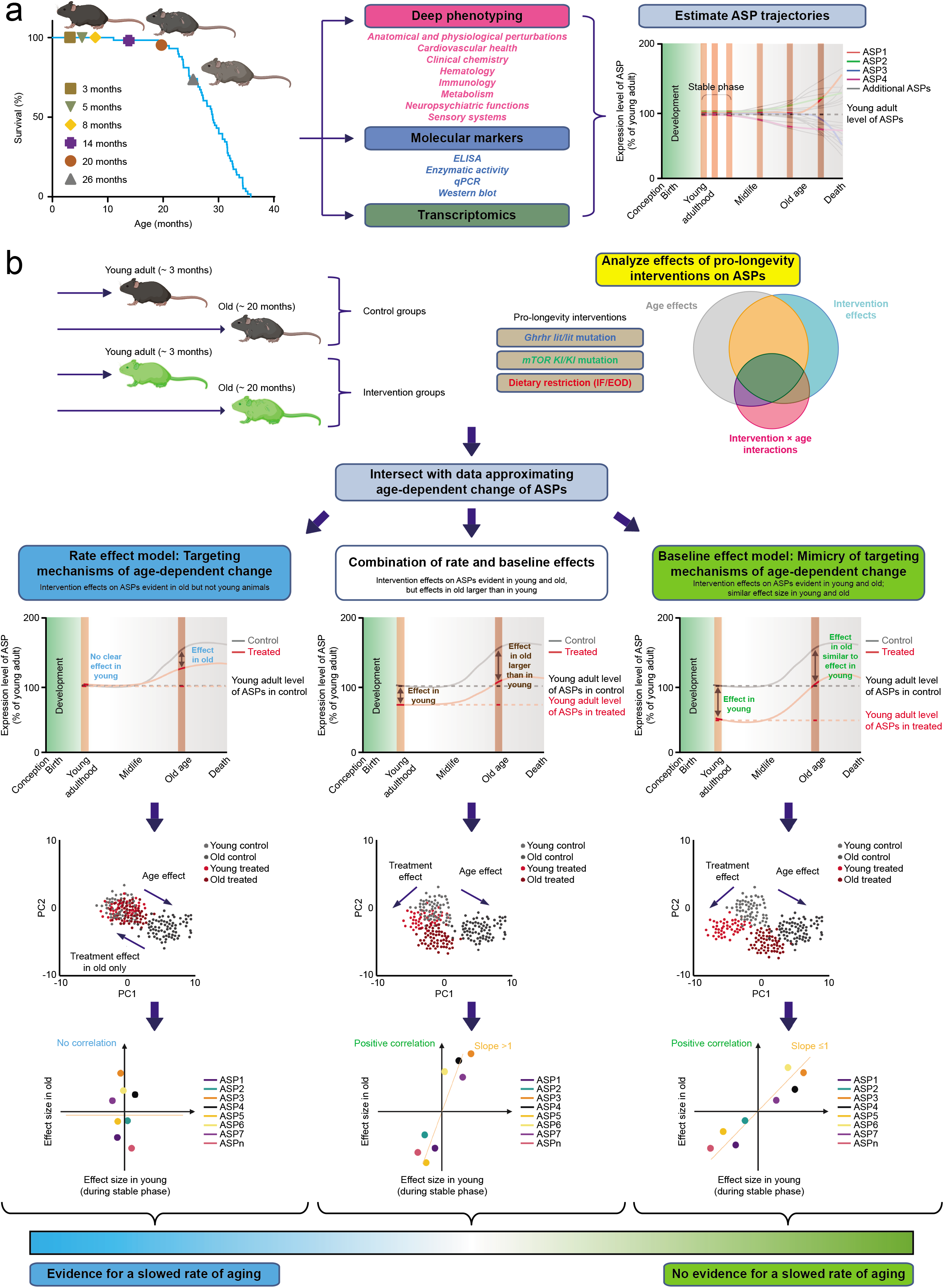
**a**, **Large-scale assessment of age-sensitive phenotypes (ASPs) to measure age-dependent phenotypic change in mice.** We assessed a large number of phenotypes across the lifespan of C57BL/6J mice (3, 5, 8, 14, 20 and 26 months old), including hundreds of phenotypes derived from multi-dimensional deep phenotyping, a range of molecular markers as well as transcriptomic profiles. The panel to the left shows how these age groups relate to the natural survival curve of C57BL/6J mice in our setting (few losses up to 20 months; approx. 30% of the population is lost between 20 and 26 months). This baseline study served to estimate aging trajectories for individual age-sensitive phenotypes (ASPs). All data were collected from a single synchronized cohort of animals using a cross-sectional study design. We hypothesized that ASPs follow different lifetime trajectories but that there is an initial stage of relative stability in young adulthood with more limited measurable changes in most of the parameters examined (see schematic to the right). **b**, **To what extent can aging be slowed?** We assessed three important pro-longevity interventions for their effects on aging: loss-of-function genetic manipulations of growth hormone signaling (*Ghrhr^lit/lit^* mice) as well as mTOR signaling (hypomorphic *mTOR^KI/KI^* mice) and a dietary restriction model (intermittent fasting/every other day feeding). For each PAAI, we generated a young as well as an old cohort of experimental animals and controls, all of which were analyzed concurrently in one single study (i.e., using a cross-sectional study design). If not stated otherwise, we performed multi-dimensional phenotypic analyses in cohorts of young (∼3 months old) and aged (∼20 months old) control as well as experimental animals for each PAAI. For each phenotype in each of these studies, we determined age effects, intervention effects and intervention × age interactions based on the data derived from young and old control as well as experimental animals. These analyses revealed that some ASPs were influenced (countered or accentuated) by the PAAIs, others not. For ASPs countered by PAAIs, we considered the following scenarios: PAAIs could influence ASPs in a way consistent with slowing the rate of age-dependent change in ASPs (rate effect), via age-independent effects on ASPs (baseline effect) or via a combination of rate and baseline effects. To address what the age at first detectable change is for each ASP influenced by an intervention, we intersected data on ASPs from these intervention studies (see panel **b**) with data from our baseline study (see panel **a**). We compared effect sizes to examine for each ASP individually whether PAAI effects differed measurably between young and old mice. In addition, we used dimensionality reduction approaches as well as intraclass correlation analyses of intervention effect sizes in young and old animals to determine whether PAAIs overall act on ASPs primarily in a way consistent with slowing their rate of age-dependent change (left panels; in this case, one would expect PAAI effects in the old but not in the young animals), via age-independent effects (right panels; under this scenario, one would expect similar PAAI effects in young and old animals) or via a combination of rate and baseline effects (middle panels; under this scenario, one would expect PAAI effects in young and old mice, however, with larger effects in old than in young animals). For further details on our analytical approach, see **Extended Data Figure 1**.

We here refer to the mechanisms of aging as the sets of processes that underlie age-dependent phenotypic change ^3, 11, 12^. Accordingly, an intervention that targets the mechanisms underlying aging should slow the transformation of a phenotypically young to a phenotypically aged organism. In other words, the intervention should attenuate the age-dependent change in ASPs (the delta in phenotype between young and old). For instance, a specific intervention or genotype could ameliorate the age-dependent loss of neurons by promoting processes concerned with maintaining the integrity of neurons over time.

An intervention could mimic a targeting of age-dependent change by affecting ASPs directly (i.e., independently of age-dependent change in these phenotypes). For instance, a specific genetic variant may increase the number of neurons by promoting neurogenesis during brain development, without affecting the rate of subsequent age-dependent neuron loss. This variant would regulate neurodevelopmental processes but would not affect the mechanisms underlying age-dependent change. Although this would also result in increased neuronal numbers in old age, it cannot be taken as evidence of a slowed progression of aging because the rate of age-dependent change remains unaltered ^18, 19^. Such a mimicry of effects on age-dependent change can be uncovered by dissociating the intervention’s effects on ASPs from age-dependent changes in ASPs. Experimentally, this can be achieved by testing the intervention in young animals, prior to the onset of age-dependent change in ASPs.

These considerations are similar to the distinction between disease-modifying vs. symptomatic treatments made in clinical medicine ^20–23^. While both can be useful for patients, the former approach implies targeting the root causes of disease, whereas the latter does not. For instance, while a drug that enhances cognitive function in healthy people could serve well as a symptomatic treatment for subjects affected by Alzheimer’s disease (AD), it does not provide clues regarding the mechanisms underlying cognitive decline in AD. Likewise, a drug that enhances cognitive function in pre-symptomatic AD patients, well before the onset of cognitive decline, is not lending insights into the mechanisms underlying AD-related cognitive decline because that’s not what it is targeting. Clues regarding underlying pathogenetic processes can, however, be derived from a disease-modifying treatment that changes the rate of cognitive decline in AD.

Building on the considerations above, it is relatively straightforward to design experiments that distinguish between an intervention targeting age-dependent change and a mimicry of such an effect (Fig. 1; detailed analysis workflow is illustrated in **Extended Data Fig. 1**). One needs to 1) generate knowledge of lifetime profiles of ASPs in order to determine when age-dependent changes in ASPs are first detectable (Fig. 1a) to then 2) design experiments that include young treated reference groups, which are subjected to a putative anti-aging intervention (PAAI) prior to age-dependent changes in ASPs (Fig. 1b).

Based on these fundamental considerations, we sought to estimate aging trajectories for a compendium of ASPs. Towards this end, we profiled hundreds of phenotypes, and thousands of molecular markers, across the lifespan of mice; these analyses included multi-dimensional deep phenotyping, assessments of a range of molecular markers as well as transcriptomic profiling and were carried out in 3, 5, 8, 14, 20 and 26 month old male C57BL/6J mice (Fig. 1a). We hypothesized that individual ASPs follow different lifetime trajectories and that for many ASPs there is an initial stage of relative stability in young adulthood, with limited changes in many of the parameters examined (Fig. 1a, see schematic to the right). If this were correct, young groups (younger than the age at first detected age-dependent change in many ASPs) could be used to determine whether a PAAI interacts with age-dependent changes by either modifying their root causes or by acting on ASPs in an age-independent manner. Consistent with our hypothesis, we demonstrate that most of the phenotypes examined in this study feature a period of relative stability in young adulthood (i.e., between 3 and 5 months of age).

We then applied the strategy outlined above to assess key longevity interventions in animal models for their effects on aging (slowing aging rate vs. age-independent effects). Major insights into longevity-associated pathways have predominantly derived from lifespan studies of genetically modified organisms ^1, 3^. mTOR signaling and growth hormone signaling are amongst the most central regulators of lifespan according to prior longevity studies in *C. elegans*, *D. melanogaster* and mice. The mTOR pathway also represents a major focus of efforts to develop pro-longevity drugs ^24^. Accordingly, to cover key genetic longevity interventions and study their effects on aging in mice, we here chose genetic models targeting the mTOR pathway (hypomorphic *mTOR^KI/KI^* mice ^25–27^) as well as growth hormone signaling (*Ghrhr^lit/lit^* mice ^28, 29^). In parallel to our studies in mice, we applied multi-dimensional phenotyping combined with stratification based on genetic expression variants in *GHRHR* and *MTOR* in a human population across a wide age range, spanning from 30 to 95 years ^30^. The analyses in humans complement our work in animal models and allowed us to address, in parallel to the work in mice, whether or not a potential genetic modification of human ASPs occurs in an age-independent fashion or not.

In addition to these genetic factors, we applied our deep phenotyping strategy to assess an important environmental longevity intervention for its effects on aging (slowing aging rate vs. age-independent effects). Among the most intensely studied environmental factors are dietary restriction effects on longevity and age-related changes, with many thousands of publications since the 1930s when the effects of food restriction on lifespan in rodents were first discovered ^31^. Accordingly, in our study, we chose to examine a dietary restriction model (specifically, a form of intermittent fasting/every other day feeding) that has been previously linked to lifespan extension in mice ^10, 32^.

Finally, we integrated our aging trajectory dataset with the analyses of PAAIs to address whether PAAIs primarily counteracted signs of aging in ways consistent with a slowing of age-dependent changes in ASPs (Fig. 1b, panels to the lower left; rate effects) or via a mimicry of such effects (Fig. 1b, panels to the lower right; baseline effects).

PAAIs can also affect ASPs via a mixture of baseline and rate effects (Fig. 1b, panels in the lower middle). This pattern corresponds to having effects in both age groups with effects being larger in old mice than in young mice. One possible interpretation of such a pattern is that PAAIs could have age-independent effects in addition to slowing aging-associated change in phenotype. Alternatively, this constellation of findings could be caused simply by differences in treatment exposure time between young (shorter-term exposure leading to weaker effects) and old animals (longer exposure leading to stronger effects). Our current study design does not allow us to distinguish between these two possible interpretations. Hence, this intermediate category (Fig. 1b, panels in the lower middle) identifies ASPs with candidate status for a slowed rate of age-dependent change which, however, requires further study and corroboration.

Altogether, our analyses revealed that, among all PAAIs examined, many anti-aging effects were age-independent in nature (i.e., interventions had similar effect sizes in young and old mice), suggesting these phenotypes were not affected by a deceleration of age-dependent change. We also identified phenotypes influenced by PAAIs in a way consistent with a slowed aging rate, although these reflected a minority of ASPs analyzed. Our findings have important implications regarding the extent to which different aspects of the aging process can be modulated, at least by the set of PAAIs investigated in the present study.

## Results

### Age-dependent phenotypic change in our older groups of mice

To map age-dependent trajectories of a broad range of phenotypes over the lifespan of mice, we carried out deep phenotyping analyses in 3, 5, 8, 14, 20 and 26 month old male C57BL/6J mice. This covered phenotypes within the areas of cardiovascular health, neuropsychiatric functions, sensory systems, clinical chemistry, hematology, immunology, metabolism, as well as anatomy and physiology (for details regarding the assays used and phenotypes analyzed, see **Extended Data Table 1** and **2**, **Supplementary Data 1** and the Methods section; for details regarding pathological findings in this cohort of animals, see **Extended Data Fig. 2**) (Fig. 2a).

**Figure 2:**
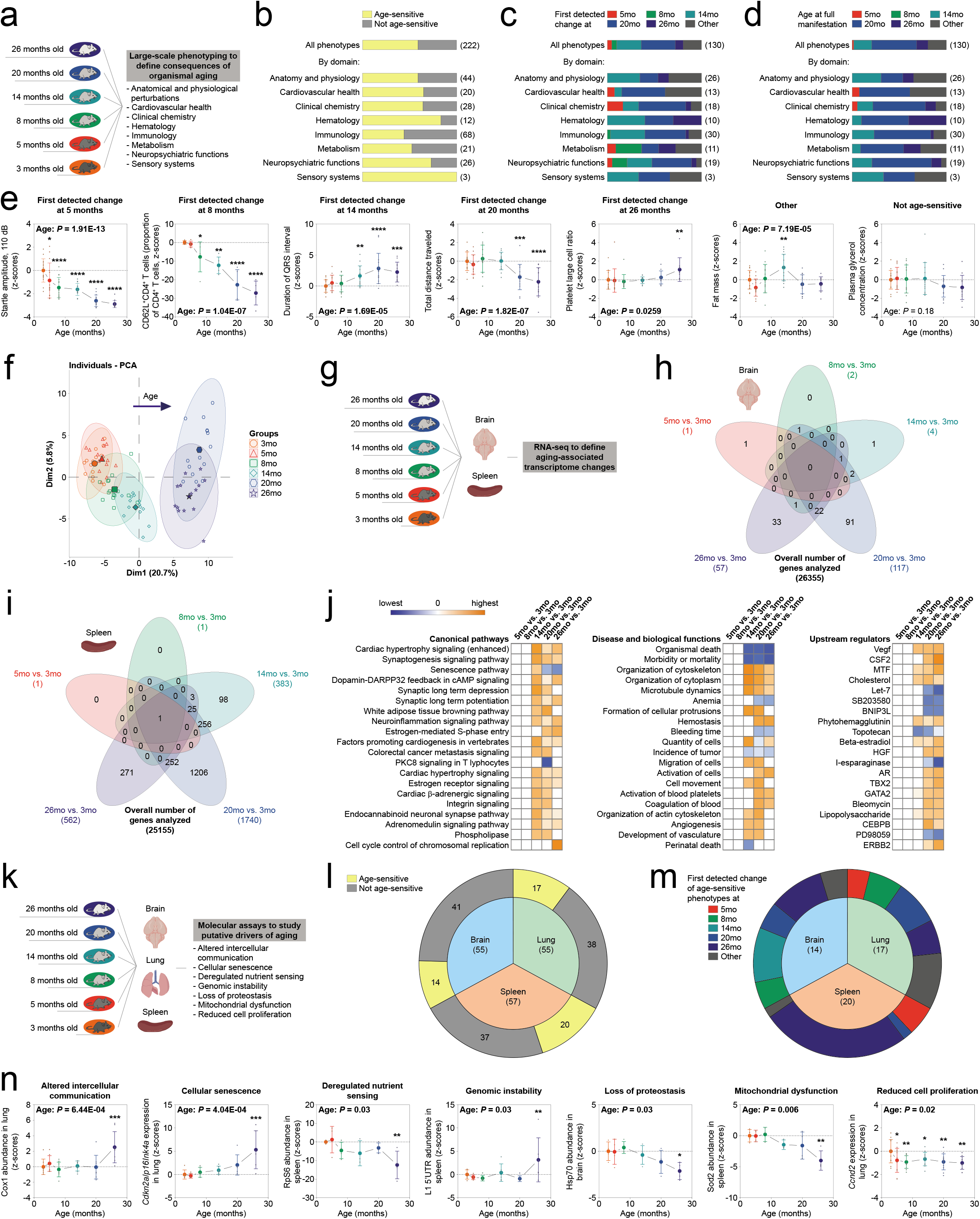
Multidimensional analyses of age-dependent phenotypic change in C57BL/6J mice. **a**–**f**, Deep phenotyping results in wildtype C57BL/6J mice. **a**, Schematic illustration of deep phenotyping study design (number of mice per group: 3-month, n=15; 5-month, n=14; 8-month, n=15; 14-month, n=14; 20-month, n=15; 26-month, n=14). **b**, Relative proportion of age-sensitive phenotypes among all phenotypes examined. **c**,**d**, Age at first detectable change (**c**) and age at full manifestation (**d**) of age-sensitive phenotypes (ASPs) shown as proportion of all ASPs. **e**, Representative examples of ASPs with various ages at first detectable phenotypic change (number of mice per group: acoustic startle amplitude at 110 dB, n≧13; CD4^+^CD62L^+^ T cells % of CD4^+^ T cells, n=5; duration of the QRS interval measured by electrocardiography, n≧11; total distance traveled in the open field test, n≧14; platelet large cell ratio, n≧10; fat mass as measured by nuclear magnetic resonance, n≧13; plasma glycerol concentration, n≧13; for full information, see **Supplementary Data 1**). **f**, Principal component analysis of deep phenotyping data. **g**–**j**, Summary of RNA-seq data. **g**, Schematic illustration of RNA-seq study design (number of mice per group: 3-month, n=7; 5-month, n=9; 8-month, n=8; 14-month, n=9; 20-month, n=7; 26-month, n=5). **h**,**i**, Venn diagram shows, for brain (**h**; number of mice per group: n≧5) and spleen (**i**; number of mice per group: n≧5), the number of differentially expressed genes (FDR<0.05) relative to the 3-month old reference group together with the intersection of the corresponding gene sets. **j**, Ingenuity Pathway Analysis shows top canonical pathways, diseases and biological functions as well as predicted upstream regulators of genes differentially expressed in spleen relative to the 3-month old group. Positive z-scores (in orange) indicate activating effects, while negative z-scores (in blue) indicate inhibitory effects on corresponding processes. Pathway analyses of brain data are shown in **Extended Data** Fig. 3**. k**–**n**, Summary of molecular analyses designed to study putative driver mechanisms of aging in spleen, lung and brain. **k**, Schematic illustration of study design (for sample size information, see **Supplementary Data 5**). **l**, Proportion of age-sensitive molecular parameters obtained in individual tissue types. **m**, Proportion of the different age-at-first-detectable-change categories among all age-sensitive molecular markers in individual tissue types. **n**, Representative examples of molecular markers covering the different hallmarks of aging (number of mice per group: Cox1 abundance in lung, n=8; *Cdkn2a*/*p16ink4a* expression in lung, n=6; RpS6 abundance in spleen, n≧3; L1 5’UTR abundance in spleen, n≧5; Hsp70 abundance in brain, n=4; Sod2 abundance in spleen, n≧3; *Ccnd2* expression in lung, n=12; for full information, see **Supplementary Data 5**). Line plots (**e**,**n**) show means +/- S.D. (individual data points are superimposed; we did not use jittering to separate data points with identical values). *p<0.05, **p<0.01, ***p<0.001, ****p<0.0001 relative to 3-month young adult reference group.

Overall, this analysis included 222 phenotypes, ∼59% of which we found to be age-sensitive (p < 0.05 by one-way ANOVA with between-subjects factor age, Kruskal-Wallis-test or Fisher’s exact test, as appropriate). ASPs were observed across all functional domains examined (Fig. 2b). Based on the outcomes of posthoc analyses relative to the young reference group (3 months old group; for details, see Material and Methods), we next assigned ASPs into any one of the following categories: ASPs first detectable at either 5, 8, 14, 20 or 26 months, or others (Fig. 2c,d).

Only ∼5% of ASPs featured very early alterations, with an age at first detected change of 5 months; a progressively diminishing acoustic startle response, indicative of early-onset age-related hearing loss associated with a degeneration of cochlear hair cells and spiral ganglion neurons in C57BL/6J mice ^33^, represents an example in this category (Fig. 2e). We also noted very few ASPs with an age at first detected change of 8 months (∼5% of all ASPs). For instance, the abundance of naïve CD4^+^ T cells (CD62L^+^CD4^+^ T cells), which is well known to decline with advancing age ^16, 34–36^ and is linked to age-related impairments in adaptive immune responses ^34–36^, started to show measurable decrements at 8 months and continued to decrease further in the older age groups (Fig. 2e). We noted that ∼26% of ASPs showed a difference first measurable at 14 months. An increased duration of the QRS interval, a well-known electrocardiographic aging phenotype in mice ^10, 37^ and men ^38^ that might reflect slowed ventricular depolarization due to altered intercellular communication between cardiomyocytes ^37^, serves as an example to illustrate this pattern of age-related change (Fig. 2e).

Many ASPs (∼36%) were characterized by changes that became detectable only later in life, with reliable alterations first identified at 20 months. The age-related reduction of exploratory activity in an unfamiliar environment (open field), for instance, constitutes a well-known ASP ^10, 16, 39^ and was first observed in 20-month old mice (Fig. 2e). Finally, few changes (∼8%) were first noted in 26-month old mice. Alterations in platelet morphology showed a detectable departure from baseline at 26 months (Fig. 2e). In addition to age-related alterations showing a consistent direction of change once they had emerged, we also noted a subset of phenotypes with other lifetime profiles (∼19%, denoted as ‘other’): Fat mass for instance first increased to a peak in midlife and then decremented in older age groups (Fig. 2e).

We carried out principal component analysis (PCA) to determine how the animals from all our age groups cluster in 2D-PCA space based on all phenotypes (measured on a continuous scale) included in the deep phenotyping analysis (Fig. 2f). Age effects were mostly seen in PC1. There was no apparent difference between 3-month and 5-month old animals on PC1. A PC1 shift to the right appeared to be first evident at 8 months and progressively increased up to 20 months.

In conclusion, our deep phenotyping analysis identified a large number of ASPs and showed only very limited age-related changes in these phenotypes between 3 and 5 months of age. The analyses also indicated that, overall, most changes in ASPs from baseline were detected in the second year of life of the animals (between 14 and 20 months).

Next, we employed RNA-seq analyses to determine the age in life when transcriptomic changes relative to the young adult baseline are first discernable in our cohort of animals (Fig. 2g-j; **Extended Data Fig. 3**; **Supplementary Data 2**-**4**). As starting material, we used brain and spleen tissue, respectively, of 3, 5, 8, 14, 20 and 26 month old male C57BL/6J mice. Consistent with the data described above, we observed very limited changes in gene expression when comparing 3-month old mice to 5-or 8-month old animals for both brain and spleen tissue (Fig. 2h,i; **Extended Data Fig. 3**). While in spleen many changes relative to the young adult baseline began to be detectable at 14 months (Fig. 2i,j; **Supplementary Data 2**), significant differences relative to the 3-month baseline in brain were largely restricted to the two oldest groups (20 and 26 months; Fig. 2h; **Extended Data Fig. 3**; **Supplementary Data 3**).

To establish how molecular and cellular mechanisms that have been suggested to drive aging are altered across the murine lifespan ^4^, we analyzed a panel of molecular markers that we designed to cover many hallmark processes of aging (summarized in **Extended Data Tables 3-5**; **Extended Data Fig. 4**). These included markers to assess alterations in intercellular communication, cellular senescence, deregulated nutrient sensing, genomic stability, loss of proteostasis, mitochondrial dysfunction and reduced cell proliferation (Fig. 2k). These analyses were focused on spleen, lung and brain of 3, 5, 8, 14, 20 and 26 month old male C57BL/6J mice. Based on the set of molecular markers tested, age-associated alterations were noted in 14 out of 55 in the brain, 17 out of 55 in the lung and 20 out of 57 in the spleen (Fig. 2l; **Supplementary Data 5**). Among all age-sensitive markers, most showed relative stability between 3 and 5 months and clear changes, compared to the young adult baseline (3 months), were detectable primarily in the oldest groups (20 and 26 months) (Fig. 2m; **Supplementary Data 5**). An exception to this notion were age-related changes in cell proliferation markers that decremented early (between 3 and 5 months) and remained stable afterwards (Fig. 2m,n; **Supplementary Data 5**). In summary, our analyses of molecular markers were consistent with the deep phenotyping data described above in showing relative stability in early life; most age-related changes relative to the young adult baseline were identified past the first year of life.

Altogether, the data discussed thus far, including deep phenotyping, molecular and transcriptomic data, revealed that few age-sensitive markers show alterations detectable early in life (between 3 and 5 months). Rather, most changes relative to the young adult baseline (3 months) detected in our study became apparent in the second year of life.

### Do key longevity factors slow aging in mice?

We wanted to establish whether key longevity and putative anti-aging interventions (PAAIs), on a large scale, counter age-sensitive phenotypes (ASPs). We also wanted to address whether effects on ASPs are best explained via (1) slowing the development of age-related changes in ASPs, (2) age-independent effects on ASPs or (3) a combination of (1) and (2). To explore which of these scenarios is supported best empirically, we analyzed deep phenotyping and transcriptomic effects of key longevity interventions. For each PAAI, we generated a young as well as an old cohort of experimental animals and controls, all of which were analyzed concurrently. We chose the young group to be 3 months of age when the analyses commenced and ∼5 months at their completion, implying that the measurements were carried out during a period of relative stability of most ASPs (Fig. 2; **Extended Data Table 1**; **Supplementary Data 1**). Accordingly, effects on ASPs seen in the young groups should be largely interpreted as aging-independent effects. We chose the old groups to be ∼20 months of age when the analyses started (completion at ∼22 months) because the animals were old enough to have accumulated clear changes in ASPs but were still at an age before the steep increase in age-related mortality (Fig. 1a). This design choice was made to minimize confounding effects of differential survival in our analysis of mutant mice vs. controls.

### A loss-of-function *Ghrhr* mutation attenuated ASPs often via age-independent mechanisms

We focused our analyses on two single-gene mutants, both associated with extension of lifespan in mice ^25, 26, 29^ and each affecting a pathway generally considered to be pivotal in regulating lifespan and aging ^16, 39–44^. First, we analyzed a mouse line with a loss-of-function mutation in *Ghrhr* (coding for the growth hormone releasing hormone receptor) ^28^. These *Ghrhr^lit/lit^* mutant mice display deficiencies in growth hormone signaling, a dwarf phenotype, extension of lifespan and an amelioration of several age-sensitive phenotypes when analyzed in old age ^29^.

PCA of the deep phenotyping dataset we generated for the *Ghrhr^lit/lit^* mutant line (Fig. 3; **Extended Data Table 6**; **Extended Data Fig. 5; Supplementary Data 6**) indicated similar genotype effects in the young and aged group of animals (see PC1 and PC2 in Fig. 3b). Based on axis contributions, the effect of genotype was about twice as large as the effect of age. Age effects on the first two principal components were similar in WT and *Ghrhr^lit/lit^* mutants (Fig. 3b) with no clear evidence for an interaction of age and genotype.

**Figure 3:**
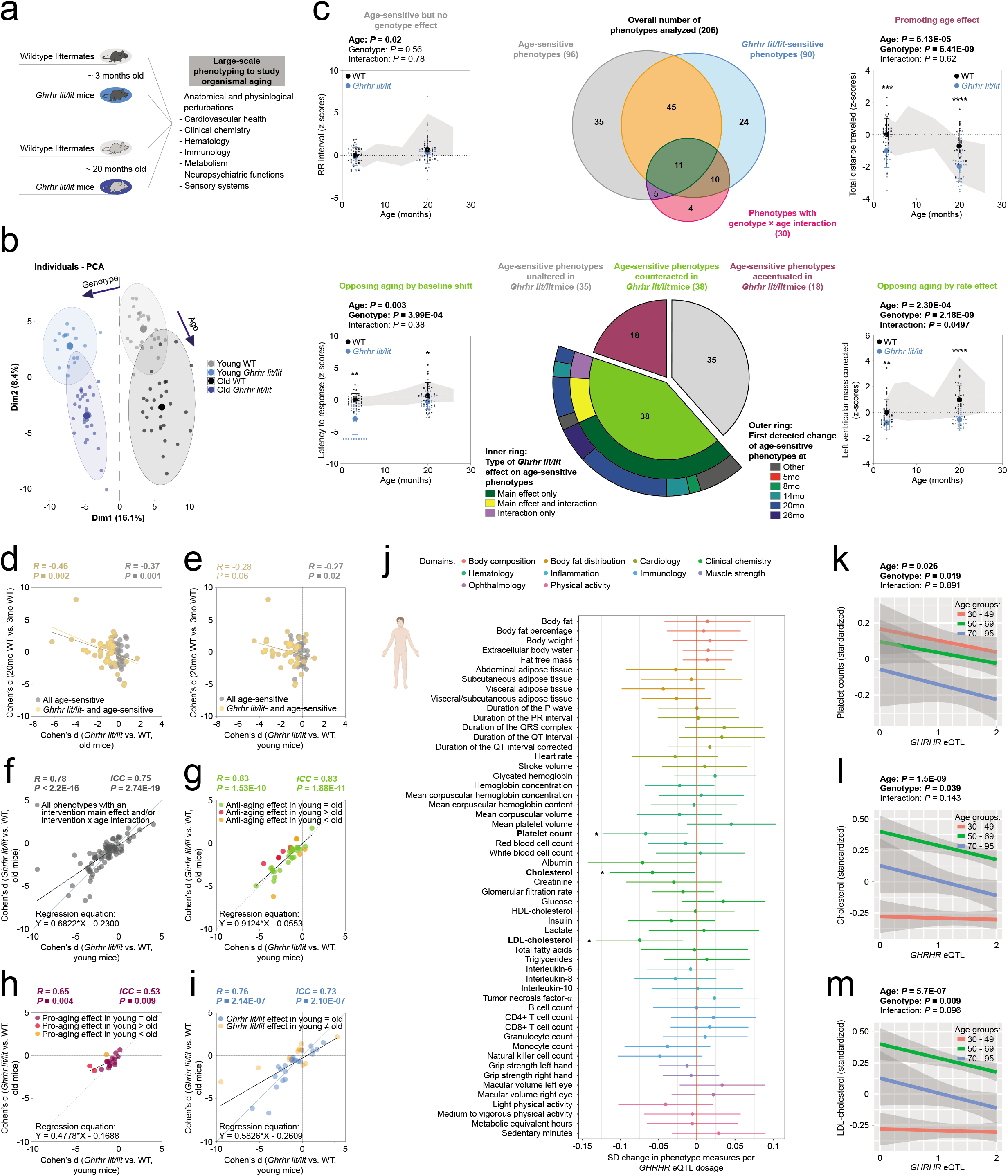
‘Anti-aging’ effects induced by the *Ghrhr^lit/lit^* mutation manifested mostly in young mice (prior to detectable age-dependent phenotypic changes). **a**–**i**, Deep phenotyping results in *Ghrhr^lit/lit^* mice. **a**, Schematic illustration of deep phenotyping study design (number of mice per group: young WT, n=30; young *Ghrhr^lit/lit^*, n=20; old WT, n=29; old *Ghrhr^lit/lit^*, n=30). **b**, Principal component analysis of deep phenotyping data. **c,** Top middle panel: Venn diagram shows the number of age-sensitive phenotypes, genotype-sensitive phenotypes, phenotypes with a genotype × age interaction and their intersection. **c**, Bottom middle panel: Sunburst chart shows the number of age-sensitive phenotypes either unaltered (in grey), counteracted (in green) or accentuated (in magenta) by the *Ghrhr^lit/lit^* mutation. For age-sensitive phenotypes counteracted by the *Ghrhr^lit/lit^* mutation, the inner ring shows the proportion of phenotypes with a main effect of genotype (in dark green), a genotype × age interaction (in violet) or both a main effect and an interaction (in yellow). The outer ring shows when changes in the corresponding ASPs were first detected based on data available from our baseline study. Line charts (top left/right and bottom left/right panels) show representative examples of phenotypes influenced by age and/or intervention in the different possible ways (number of mice per group: duration of the RR interval measured during electrocardiography, n≧20; total distance traveled in the open field test, n≧20; latency to first response in the hot plate test, n≧20; corrected mass of the left ventricle as measured by echocardiography, n≧20). Data were transformed to z-scores (normalized to the young WT group) and are plotted as mean +/- S.D. for each group (individual data points are superimposed). *p<0.05, **p<0.01, ***p<0.001 relative to age-matched wildtype littermate controls. Life-time trajectories of the corresponding phenotypes are shown by the grey-shaded area in the background (upper bound: mean + S.D.; lower bound: mean – S.D.) which represent the measurements obtained in 3-, 5-, 8-, 14-, 20- as well as 26-month old C57BL/6J wildtype mice (values standardized to the 3-month old reference group). **d**,**e**, Scatter plot shows the effect size of *Ghrhr^lit/lit^* genotype in old mice (**d**) or young mice (**e**) plotted vs. the effect size of age (20 months vs. 3 months; data from baseline study shown in Fig. 2 to ensure independence of measures used in correlation analysis) for all ASPs (all data points) and those intersecting with genotype (via a genotype main effect and/or interaction; in yellow). **f**–**i**, Scatter plots show the effect size of *Ghrhr^lit/lit^* genotype in young mice plotted vs. the effect size of *Ghrhr^lit/lit^* genotype in old mice for different sets of phenotypes: **g**, ASPs ameliorated by genotype via a main effect and/or an interaction (i.e., corresponding to the central green section of the sunburst chart in **c**); green dots denote phenotypes in which genotype effects in young and old mice did not differ significantly; orange denotes phenotypes in which anti-aging effects of genotype were significantly larger in old mice than in young mice; red denotes phenotypes in which anti-aging effects of genotype were significantly larger in young mice than in old mice. **h**, ASPs accentuated by genotype. **i**, Phenotypes featuring a main effect of *Ghrhr^lit/lit^* genotype and/or a genotype × age interaction but not a main effect of age; blue dots denote phenotypes in which the genotype effect did not differ significantly between young and old mice. Yellow denotes phenotypes in which the genotype effect size differed significantly between young and old mice. **f**, all phenotypes shown in **g**-**i** collapsed into one panel. ICC = intraclass correlation. For further details, see **Supplementary Data 6**. **j**–**m**, Phenotyping results in a large deep-(endo)phenotyped human cohort. **j,** change in phenotype (in standard deviations (SD) from the mean) associated with *GHRHR* eQTL dosage with the horizontal whiskers indicating the 95% confidence intervals of the mean effect estimate; * denotes p<0.05 for the linear association between *GHRHR* eQTL dosage and (endo)phenotype. **k**–**m**, Change of platelet count (**k**), total cholesterol (**l**) and LDL-cholesterol (**m**) associated with *GHRHR* eQTL dosage in 30 – 49 years old (red line), 50 – 69 years old (green line) and 70 – 95 years old humans (blue line); the lines represent the best-fit least squares regression lines with surrounding 95% confidence intervals of the mean indicated in grey. The eQTL dosage was coded as GG=0, AG=1, and AA=2 (GG is associated with lowest expression levels, AA with highest; see **Extended Data Fig. 8a**).

Analyses of individual phenotypes (based on two-way ANOVA or aligned rank transform; for details, see also Material and Methods) revealed that, out of 206 phenotypes examined, 96 showed a significant main effect of age, 90 showed a significant main effect of genotype and 30 showed a significant interaction between genotype and age (Fig. 3c). Out of the 96 age-sensitive phenotypes, 35 were not significantly affected by genotype, 45 showed a significant main effect of genotype (but no interaction) and 16 featured a significant genotype × age interaction (Fig. 3c). Further analyses of ASPs based on the results of posthoc tests are described in **Supplementary Results**, **Extended Data Fig. 7** and **Supplementary Data 6**.

To assess whether genotype effects countered age effects or whether genotype and age effects influenced a phenotype in the same direction, we compared for each phenotype the directionality of Cohen’s d effect sizes of age with those of Cohen’s d effect sizes of genotype in the old group of mice. These analyses revealed that out of the set of 96 age-sensitive phenotypes 18 were further accentuated by the *Ghrhr^lit/lit^* genotype, while 38 were opposed by the *Ghrhr^lit/lit^* genotype (Fig. 3c; **Extended Data Table 6**; **Supplementary Data 6**; 5 ASPs could not be evaluated because Cohen’s d effect sizes could not be computed due to 0 values in the denominator). Most of the 38 ASPs counteracted by *Ghrhr^lit/lit^* genotype showed a significant main effect of genotype, but no significant interaction between genotype and age (Fig. 3c). Closer inspection of the ASPs featuring a significant interaction term (considering the directionality of change) indicated that ca. 10.4% of all ASPs identified correspond to ASPs counteracted by *Ghrhr^lit/lit^* in ways consistent with the “rate effect model” or “combined rate/baseline effect model” introduced in Fig. 1b. Based on a significant genotype main effect, but a lack of an interaction, ca. 29.2% corresponded to ASPs consistent with the “baseline effect model” shown in Fig. 1b. The remaining ASPs were not affected (ca. 36.5%), accentuated (ca. 18.8%) by *Ghrhr^lit/lit^* or could not be evaluated (ca. 5.2%). All the *Ghrhr^lit/lit^*-opposed ASPs we were able to evaluate had an age at first detectable departure from young adult baseline of 8 months (Fig. 3c); hence, all the corresponding genotype effects on ASPs in young animals appear independent of age-related change in those ASPs (since age-dependent changes in ASPs have not yet manifested in young animals).

To examine further the relationships between age and genotype effects, we performed correlation analyses comparing the Cohen’s d effect sizes of age vs. the *Ghrhr^lit/lit^* genotype. We found a modest inverse relationship between age effects and genotype effects in old mice (Fig. 3d). We noted a similar inverse relationship when regressing the effect size of age with the effect size of the *Ghrhr^lit/lit^* mutation in the young cohort of animals (Fig. 3e).

We next asked whether genotype effects are similar across age groups within the category of ASPs countered in *Ghrhr^lit/lit^* mutants (n=38 phenotypes). Towards this end, we performed linear regression analyses of genotype effects in the young vs. the old group of mice for these n=38 phenotypes. For all ASPs countered by the *Ghrhr*^lit/lit^ genotype, these analyses revealed an overall high similarity of genotype effects across age groups (Fig. 3g; R=0.83, p=1.53E-10), indicating that homozygosity for the *Ghrhr*^lit^ mutant allele resulted in similar phenotypic consequences on ASPs irrespective of the age of the animal. The slope of the regression line was 0.91 ± 0.1 (95% CI: 0.70, 1.12; p=0.3975), thereby supporting the notion of overall similar effect sizes in young and old mice in this category of genotype-sensitive ASPs (Fig. 3g; 1 corresponding to the same effect sizes in young and old; values significantly < 1 to effect sizes overall larger in young mice; values significantly > 1 to effect sizes overall larger in old mice). Similar results were obtained using intraclass correlation analyses (Fig. 3g; ICC=0.83, p=1.88E-11) which reflect not only the degree of correlation but also the agreement between measures in the young and old group. For instance, consistent with prior research ^16, 45^, advancing age led to an increased latency to respond on the hot plate test, indicative of aging-associated alterations in nociceptive function, and the *Ghrhr*^lit^ allele antagonized this aging-associated phenotype (Fig. 3c). However, we found similar effects of the *Ghrhr*^lit^ allele in old mice as well as in young animals that were younger than the age at which age-dependent changes in this phenotype are first detectable (Fig. 3c). Statistical comparison of genotype effect sizes in young mice vs. effect sizes in old mice revealed only five cases in which there was a significantly larger *Ghrhr*^lit^ effect in the aged group of animals (Fig. 3g), for instance blood hemoglobin concentration or plasma alkaline phosphatase activity. In most cases, however, effect sizes in young and old mice were not significantly different (Fig. 3g; **Extended Data Table 6**; **Supplementary Data 6**). Hence, based on the analysis of genotype effect sizes in young vs. old mice, only ca. 5.2% of ASPs were countered by the *Ghrhr*^lit^ allele in ways consistent with either the “rate effect model” or “combined rate/baseline effect model” introduced in Fig. 1b (larger effect in old than in young). Ca. 34.4% of all ASPs were countered in ways consistent with the “baseline effect model” shown in Fig. 1b (effect in old not larger than in young). As mentioned above, the remaining ASPs were either not affected (ca. 36.5%), accentuated (ca. 18.8%) or could not be evaluated (5.2%). Clear correlations between genotype effects within young vs. aged animals were also observed when we analyzed either ASPs accentuated by genotype (Fig. 3h; ICC=0.53, p=0.009), age-insensitive phenotypes influenced by genotype (Fig. 3i; ICC=0.73, p=2.10E-07) or all of these categories combined (Fig. 3f; ICC=0.75, p=2.74E-19). Together, these observations indicate that *Ghrhr* genotype effects were, overall, largely independent of age and this was the case for the set of ASPs countered by genotype and other phenotypic categories (Fig. 3f-i).

Our studies in *Ghrhr^lit/lit^* mice showed a range of physiological consequences of *Ghrhr* loss of function and highlighted how a subset interacts with age-dependent alterations in mice. We also wanted to explore whether *GHRHR* expression differences are associated with phenotypic consequences in humans and, if so, whether these effects are age-dependent or not (Fig. 3j-m, **Extended Data Table 7**). To address this question, we analyzed multi-dimensional phenotypic data, covering a range of areas of human physiology (such as body composition, body fat distribution, cardiology, clinical chemistry, hematology, inflammation, immunology, muscle strength, ophthalmology and physical activity), collected from n=3034 30- to 95-year old human individuals (**Extended Data Table 8**). We identified *GHRHR*-sensitive phenotypes by stratifying the human phenotypic data by *GHRHR* genotype, taking advantage of a SNP (rs11772180), which is located upstream of the *GHRHR* gene and has been identified as an independent cis-eQTL, i.e. a polymorphism significantly associated with *GHRHR* expression levels (**Extended Data Fig. 8**) ^46^. This eQTL had significant effects on several age-sensitive phenotypes examined, including platelet count and cholesterol-associated measures (Fig. 3j-m). In one of these ASPs, the *GHRHR* variant associated with low expression appeared to influence the ASP in ways that counter the direction of age-dependent change (Fig. 3k); there was no genotype x age interaction (Fig. 3k; **Extended Data Table 7**), which is in line with our observation of predominantly age-independent *Ghrhr* effects on ASPs (including ASPs countered by genotype) in mice.

Together, these data indicate that changes in growth hormone signaling are associated with a range of phenotypic effects, including a subset of effects that counteract age-dependent changes. Our data support that most effects on ASPs are evident already in young animals, long before age-dependent changes in ASPs are detectable, indicating that these are age-independent effects. We also identified ASPs that were influenced by the *Ghrhr*^lit^ allele and showed larger effect sizes in old mice than in young mice. These ASPs potentially correspond to phenotypes in which aging trajectories were slowed by the *Ghrhr* mutation.

### A hypomorphic *mTOR* mutant allele attenuated ASPs via a mixture of age-independent effects and effects that were more pronounced in old mice

We then asked whether the pattern of observations in the *Ghrhr* mouse model would hold true for other candidate longevity interventions as well. We applied the same analytical approach to a hypomorphic mTOR mutant mouse line featuring mTOR expression levels reduced to 25% of those seen in WT littermate controls ^25^ (Fig. 4; **Extended Data Table 9**; **Extended Data Fig. 5; Supplementary Data 7**). PCA-based dimensionality reduction of deep phenotyping data from young and old hypomorphic mTOR mutant mice as well as WT littermate controls suggested that age and genotype effects in 2D-PCA space are largely independent of each other (Fig. 4b). However, interestingly, the distance between the young and old groups of mice appeared reduced in the case of mTOR mutant mice relative to WT controls (Fig. 4b).

**Figure 4:**
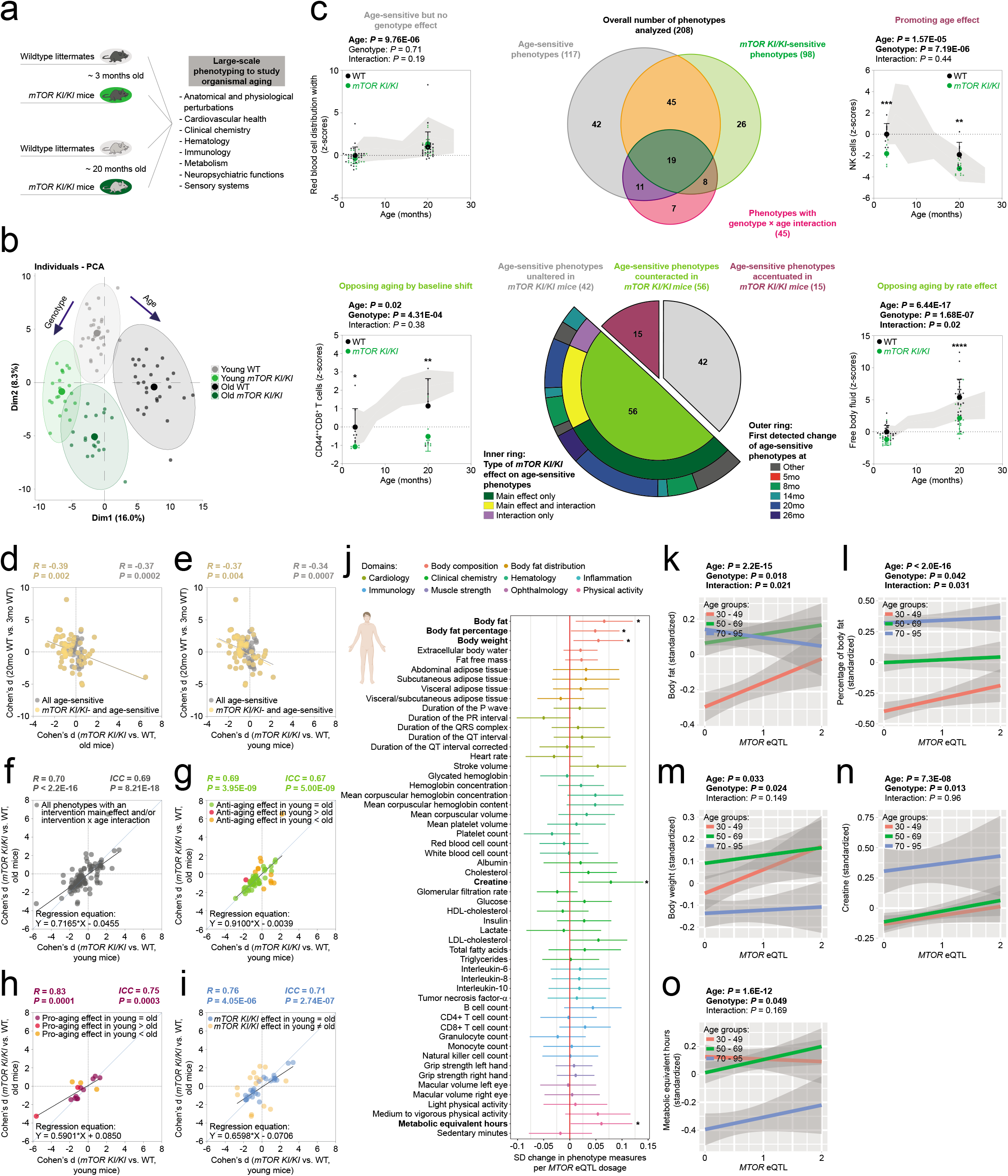
A hypomorphic mTOR mutant allele attenuated ASPs via a mixture of age-independent effects and effects that were more pronounced in old mice. **a**–**i**, Deep phenotyping results in *mTOR^KI/KI^* mice. **a**, Schematic illustration of deep phenotyping study design (number of mice: young WT, n=27; young *mTOR^KI/KI^*, n=21; old WT, n=26; old *mTOR^KI/KI^*, n=19). **b**, Principal component analysis of deep phenotyping data. **c,** Top middle panel: Venn diagram shows the number of age-sensitive phenotypes, genotype-sensitive phenotypes, phenotypes with a genotype × age interaction and their intersection. **c**, Bottom middle panel: Sunburst chart shows the number of age-sensitive phenotypes either unaltered (in grey), counteracted (in green) or accentuated (in magenta) by the *mTOR^KI/KI^* mutation. For age-sensitive phenotypes counteracted by the *mTOR^KI/KI^* mutation, the inner ring shows the proportion of phenotypes with a main effect of genotype (in dark green), a genotype × age interaction (in violet) or both a main effect and an interaction (in yellow). The outer ring shows when changes in the corresponding ASPs were first detected based on data available from our baseline study. Line charts (top left/right and bottom left/right panels) show representative examples of phenotypes influenced by age and/or intervention in the different possible ways (number of mice per group: red blood cell distribution width, n≧14; NK cells % of all leukocytes, n≧8; CD44^++^CD8^+^ T cells % of CD8^+^ T cells, n≧8; free body fluid as measured by nuclear magnetic resonance, n≧15). Data were transformed to z-scores (normalized to the young WT group) and are plotted as mean +/- S.D. for each group (individual data points are superimposed). *p<0.05, **p<0.01, ***p<0.001 relative to age-matched wildtype littermate controls. Life-time trajectories of the corresponding phenotypes are shown by the grey-shaded area in the background (upper bound: mean + S.D.; lower bound: mean – S.D.) which represent the measurements obtained in 3-, 5-, 8-, 14-, 20- as well as 26-month old C57BL/6J wildtype mice (values standardized to the 3-month old reference group). **d**,**e**, Scatter plot shows the effect size of *mTOR^KI/KI^* genotype in old mice (**d**) or young mice (**e**) plotted vs. the effect size of age (20 months vs. 3 months; data from baseline study shown in Fig. 2 to ensure independence of measures used in correlation analysis) for all ASPs (all data points) and those intersecting with genotype (via a genotype main effect and/or interaction; in yellow). **f**– **i**, Scatter plots show the effect size of *mTOR^KI/KI^* genotype in young mice plotted vs. the effect size of *mTOR^KI/KI^* genotype in old mice for different sets of phenotypes: **g**, ASPs countered by genotype via a main effect and/or an interaction (i.e., corresponding to the central green section of the sunburst chart in **c**); green dots denote phenotypes in which genotype effects in young and old mice did not differ significantly; orange denotes phenotypes in which anti-aging effects of genotype were significantly larger in old mice than in young mice; red denotes phenotypes in which anti-aging effects of genotype were significantly larger in young mice than in old mice. **h**, ASPs accentuated by genotype. **i**, Phenotypes featuring a main effect of *mTOR^KI/KI^* genotype and/or a genotype × age interaction but not a main effect of age; blue dots denote phenotypes in which the genotype effect did not differ significantly between young and old mice. Yellow denotes phenotypes in which the genotype effect size differed significantly between young and old mice. **f**, all phenotypes shown in **g**-**i** collapsed into one panel. ICC = intraclass correlation. For further details, see **Supplementary Data 7**. **j**–**o**, Phenotyping results in humans. **j**, Change in phenotype associated with *MTOR* eQTL dosage in a large deep-(endo)phenotyped human cohort, with the horizontal whiskers indicating the 95% confidence intervals of the mean effect estimate; * denotes p<0.05 for the linear association between *MTOR* eQTL dosage and (endo)phenotype. **k**–**o**, Change of body fat (**k**), percentage of body fat (**l**), body weight (**m**), plasma creatine concentration (**n**) and metabolic equivalent hours (**o**) associated with *MTOR* eQTL dosage in 30 – 49 years old (red line), 50 – 69 years old (green line) and 70 – 95 years old humans (blue line); the lines represent the best-fit least squares regression lines with surrounding 95% confidence intervals of the mean indicated in grey. The eQTL dosage was coded as GG=0, CG=1, and CC=2 (GG is associated with lowest expression levels, CC with highest; see **Extended Data Fig. 8b**).

Analyses of 208 individual phenotypes covered in these studies identified 117 phenotypes with a significant main effect of age, 98 with a significant main effect of genotype and 45 with a significant interaction between genotype and age (Fig. 4c). Out of the 117 age-sensitive phenotypes, 42 were not significantly affected by genotype, 45 showed a significant main effect of genotype (but no interaction) and 30 featured a significant genotype × age interaction (Fig. 4c). Further analyses of ASPs based on the results of posthoc tests are described in **Supplementary Results**, **Extended Data Fig. 7** and **Supplementary Data 7**.

To assess whether genotype effects counteracted or accentuated age effects, we compared for each phenotype the directionality of Cohen’s d effect sizes of age with those of Cohen’s d effect sizes of genotype (in the old group of mice). Out of 75 ASPs influenced by genotype, the clear majority (56) was countered by the *mTOR^KI/KI^* genotype (Fig. 4c; **Extended Data Table 9**; **Supplementary Data 7**; 15 ASPs were accentuated by the *mTOR^KI/KI^* genotype; 4 ASPs could not be evaluated because Cohen’s d effect sizes could not be computed due to 0 values in the denominator). Interestingly, a sizeable fraction (22 of the 56 ASPs) ameliorated by the *mTOR^KI/KI^* genotype showed a significant interaction between genotype and age (Fig. 4c). It should be noted, however, that the majority of these 56 ASPs showed a main effect of genotype, without evidence for a significant interaction (Fig. 4c). Further inspection of the ASPs with a significant interaction term (considering the directionality of change) indicated that ca. 16.2% of all ASPs identified correspond to ASPs counteracted by *mTOR^KI/KI^* in ways consistent with the “rate effect model” or “combined rate/baseline effect model” introduced in Fig. 1b. Based on a significant genotype main effect, but a lack of an interaction, ca. 31.6% corresponded to ASPs consistent with the “baseline effect model” shown in Fig. 1b. The remaining ASPs were not affected (ca. 35.9%), accentuated (ca. 12.8%) by *mTOR^KI/KI^* or could not be evaluated (ca. 3.4%). All the *mTOR^KI/KI^*-ameliorated ASPs we were able to evaluate had an age at first detectable departure from young adult baseline of 8 months (Fig. 4c), indicating that all corresponding genotype effects on ASPs in our young cohort are independent of age-related changes in these ASPs.

Effect sizes of *mTOR^KI/KI^* genotype showed a moderate inverse correlation with effect sizes of age (Fig. 4d,e); this was the case, when effect sizes of genotype in either the old (Fig. 4d) or young (Fig. 4e) group were used for correlation analyses.

Next, we addressed whether genotype effects are similar across age groups within the category of ASPs counteracted in *mTOR^KI/KI^* mutants (n=56 phenotypes). Linear regression analyses of *mTOR^KI/KI^* genotype effect sizes in young vs. old animals showed clear correlations (Fig. 4g; R=0.69, p=3.95E-09), suggesting that the *mTOR^KI/KI^* genotype resulted in overall similar phenotypic consequences on ASPs in young and old mice. This was also supported by the slope of the regression line, which did not significantly differ from 1 (slope estimate: 0.91 ± 0.13; 95% CI: 0.65, 1.17; p=0.491). Similar results were obtained using intraclass correlation analyses (Fig. 4g; ICC=0.67, p=5.00E-09). Statistical comparison of genotype effect sizes in young vs. old mice revealed 11 cases in which there was a significantly larger *mTOR^KI/KI^* effect in the aged group of animals (Fig. 4g), for instance hematocrit or plasma concentration of triglycerides. In most cases, however, effect sizes in young and old mice were not significantly different (Fig. 4g; **Extended Data Table 9**; **Supplementary Data 7**). Together, based on the analysis of genotype effect sizes in young vs. old mice, only ca. 9.4% of ASPs were countered by the *mTOR^KI^* allele in ways consistent with either the “rate effect model” or “combined rate/baseline effect model” introduced in Fig. 1b (larger effect in old than in young). Ca. 38.5% of all ASPs were countered in ways consistent with the “baseline effect model” shown in Fig. 1b (effect in old not larger than in young). As mentioned above, the remaining ASPs were either not affected (ca. 35.9%), accentuated (ca. 12.8%) or could not be evaluated (3.4%).

We also observed correlations between genotype effects within young vs. aged mice when either analyzing ASPs exacerbated in *mTOR^KI/KI^* mice (Fig. 4h; ICC=0.75, p=0.0003) or genotype-but not age-sensitive phenotypes (Fig. 4i; ICC=0.71, p=2.74E-07), indicating that the *mTOR^KI/KI^* genotype had overall very similar effects in young and old mice, irrespective of whether phenotypes were age-sensitive or not.

We next wanted to establish whether transcriptomic effects on age-sensitive genes are similar in young and old *mTOR^KI/KI^* mice. We performed RNA-seq analyses in spleen tissue of young (∼3 months old; i.e., well before the onset of age-dependent transcriptomic changes; see Fig. 2i) as well as old (∼20 months old) mTOR mutants and WT littermate controls. These analyses identified 54 genes that were age-sensitive (FDR<0.05) and 855 genes that were genotype-sensitive with an intersection between these populations of 9 genes (**Extended Data Fig. 9a**; **Supplementary Data 8**). The overlap of age- and mTOR-sensitive genes is greater than expected by chance (representation factor = 4.9, p=7.68E-05). No genes with a significant (FDR<0.05) genotype × age interaction were detected (**Extended Data Fig. 9a**). To assess whether the *mTOR* mutant allele counteracted or accentuated age-dependent gene expression alterations in spleen, expression levels were compared between young and old *mTOR* mutants and WT littermate controls. Aging-associated changes of five genes were counteracted and age-related alterations of four genes were accentuated in the *mTOR^KI/KI^* mice (**Extended Data Fig. 9b**).

We also wanted to explore how altered *MTOR* expression may affect age-sensitive phenotypes in human subjects. We assessed the associations between multi-dimensional phenotypic human data with polymorphisms at a SNP in the promoter region of the *MTOR* gene (rs2295079) that has previously been associated with variations in *MTOR* expression levels (**Extended Data Fig. 8**) ^46^. Out of 54 phenotypes (Fig. 4j), we identified 5 (Fig. 4j-o) to be associated with variations in *MTOR* eQTL, including body fat content (Fig. 4k,l) and body weight (Fig. 4m), which are known to be sensitive to changes in mTOR function ^16, 25, 47^. Next, for these 5 *MTOR*-sensitive phenotypes, we examined whether the effects of *MTOR* eQTL was age-specific or not. For 3 out of 5 parameters, we found significant effects of *MTOR* eQTL as well as of age, but no genotype x age interaction (Fig. 4l-o; **Extended Data Table 10**), suggesting that *MTOR* eQTL was largely associated with similar effects in younger and older individuals. Two phenotypes (body fat and % body fat) showed a significant genotype x age interaction which appeared to be driven by an *MTOR* variant effect in the youngest group of individuals (Fig. 4k,l; **Extended Data Table 10**). The *MTOR* variant associated with low expression appeared to influence some age-sensitive phenotypes in ways that counter the direction of age-dependent change (Fig. 4k,l,n). In other cases, age and genotype effects were in the same direction (Fig. 4m,o). Altogether our mTOR-based analyses in mouse and humans indicate that mTOR effects on age-sensitive phenotypes (including ASPs countered by mTOR effects) are often similar in young and old groups; accordingly, age-independent mechanisms need to be taken into account when interpreting mTOR effects on age-sensitive parameters.

### An intermittent fasting-based variant of dietary restriction ameliorated ASPs frequently through age-independent mechanisms

While genotype × age interactions were also observed, the data summarized above endorses an important role of age-independent effects on ASPs in the context of two central genetic interventions, targeting *mTOR* or *Ghrhr*. We also applied our analytical approach to a major environmental factor studied in aging and longevity-dietary restriction ^48, 49^. Specifically, we assessed whether and to what extent age-dependent phenotypic changes in mice are ameliorated by intermittent fasting (IF)/every other day feeding (EOD) (for details regarding study design, see Fig. 5a and Methods section). We had previously reported a significant lifespan extension induced by IF in this cohort of mice ^10^. Food intake, body weight and body composition data have also been previously reported ^10^. PCA of all deep phenotyping data revealed an additive nature of age and IF effects (Fig. 5b): PC1 was shifted to the right by age in both groups. Fasting acted mainly by decrementing PC2.

**Figure 5:**
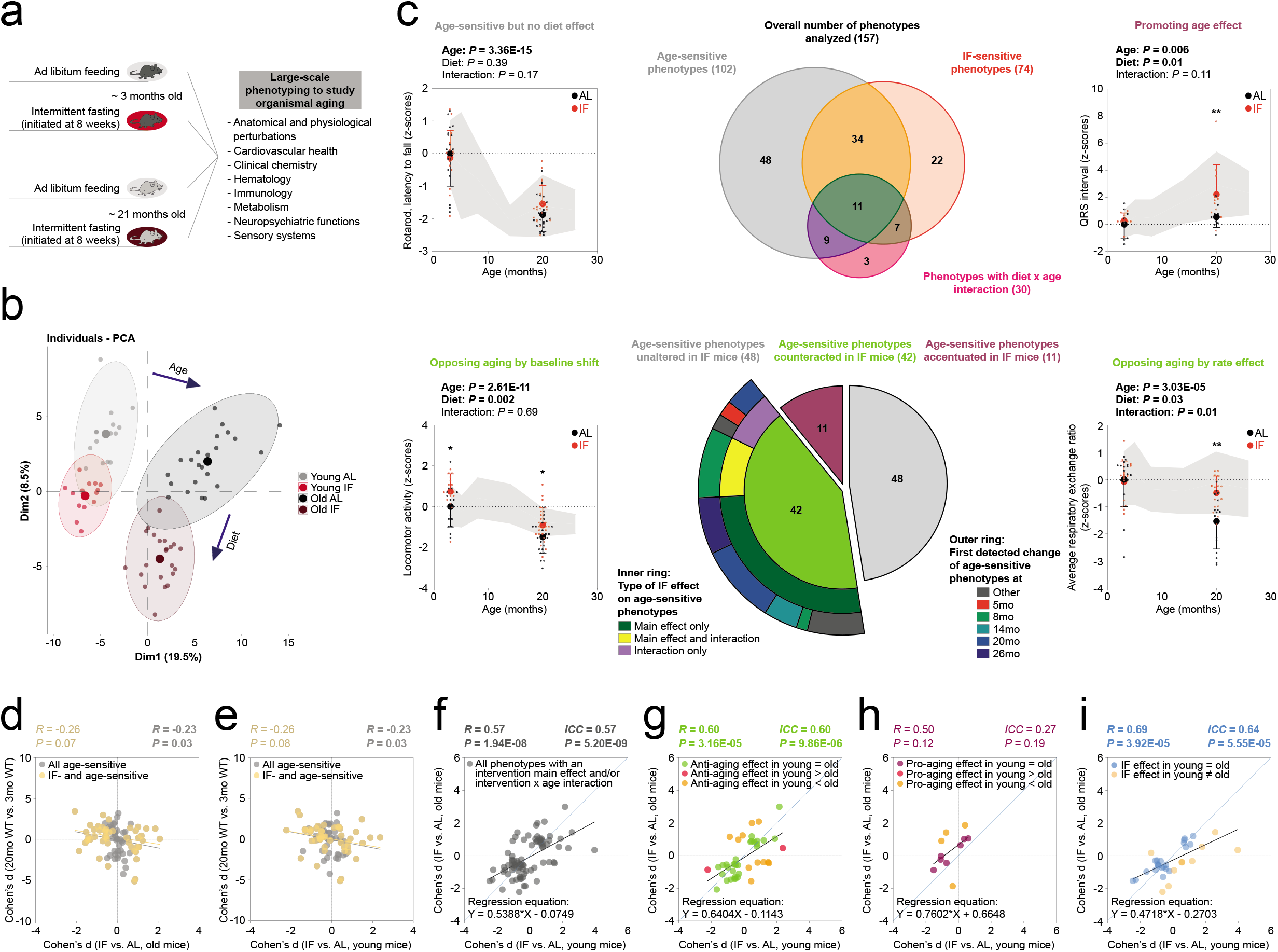
‘Anti-aging’ effects induced by intermittent fasting (IF) often manifest in young mice (prior to detectable age-dependent phenotypic changes). **a**, Schematic illustration of deep phenotyping study design (number of mice: young AL, n=16; young IF, n=16; old AL, n=23; old IF, n=23). Intermittent fasting was initiated at 8 weeks of age and was continued throughout the study. **b**, Principal component analysis of deep phenotyping data. **c,** Top middle panel: Venn diagram shows the number of age-sensitive phenotypes, diet-sensitive phenotypes, phenotypes with a diet × age interaction and their intersection. **c**, Bottom middle panel: Sunburst chart shows the number of age-sensitive phenotypes either unaltered (in grey), counteracted (in green) or accentuated (in magenta) by IF. For age-sensitive phenotypes counteracted by IF, the inner ring shows the proportion of phenotypes with a main effect of diet (in dark green), a diet × age interaction (in violet) or both a main effect and an interaction (in yellow). The outer ring shows when changes in the corresponding ASPs were first detected based on data available from our baseline study. Line charts (top left/right and bottom left/right panels) show representative examples of phenotypes influenced by age and/or intervention in the different possible ways (number of mice per group: latency to fall in the rotarod task, n≧16; duration of the QRS interval measured by electrocardiography, n≧16; locomotor activity during the SHIRPA test, n≧16; average respiratory exchange ratio determined by indirect calorimetry, n=16). Data were transformed to z-scores (normalized to the young WT group) and are plotted as mean +/- S.D. for each group (individual data points are superimposed). *p<0.05, **p<0.01, ***p<0.001 relative to age-matched wildtype littermate controls. Life-time trajectories of the corresponding phenotypes are shown by the grey-shaded area in the background (upper bound: mean + S.D.; lower bound: mean – S.D.) which represent the measurements obtained in 3-, 5-, 8-, 14-, 20-as well as 26-month old C57BL/6J wildtype mice (values standardized to the 3-month old reference group). **d**,**e**, Scatter plot shows the effect size of IF in old mice (**d**) or young mice (**e**) plotted vs. the effect size of age (20 months vs. 3 months; data from baseline study shown in Fig. 2 to ensure independence of measures used in correlation analysis) for all ASPs (all data points) and those intersecting with diet (via a diet main effect and/or interaction; in yellow). **f**–**i**, Scatter plots show the effect size of IF in young mice plotted vs. the effect size of IF in old mice for different sets of phenotypes: **g**, ASPs counteracted by IF via a main effect and/or an interaction (i.e., corresponding to the central green section of the sunburst chart in **c**); green dots denote phenotypes in which IF effects in young and old mice did not differ significantly; orange denotes phenotypes in which anti-aging effects of IF were significantly larger in old mice than in young mice; red denotes phenotypes in which anti-aging effects of IF were significantly larger in young mice than in old mice. **h**, ASPs accentuated by IF. **i**, Phenotypes featuring a main effect of diet and/or a diet × age interaction but not a main effect of age; blue dots denote phenotypes in which the diet effect did not differ significantly between young and old mice. Yellow denotes phenotypes in which the diet effect size differed significantly between young and old mice. **f**, all phenotypes shown in **g**-**i** collapsed into one panel. ICC = intraclass correlation. For further details, see **Supplementary Data 9**.

Analyses of individual phenotypes (157) revealed 102 with a significant main effect of age, 74 with a significant main effect of diet and 30 with a significant diet × age interaction (Fig. 5c; **Extended Data Table 11**; **Extended Data Fig. 5; Supplementary Data 9**). Out of the 102 phenotypes with a main effect of age (ASPs), 48 were not significantly influenced by diet, 34 showed a significant main effect of diet (but no interaction) and 20 featured a significant diet × age interaction. Further analyses of ASPs based on the results of posthoc tests are described in **Supplementary Results**, **Extended Data Fig. 7** and **Supplementary Data 9**.

Next, we wanted to more closely examine ASPs influenced by diet (either via a main effect or a diet × age interaction). To do so, we computed Cohen’s d effect sizes of age and of diet (in the old cohort of mice) to examine whether they acted in opposing directions or not (Fig. 5c). These analyses showed that age and diet effects operated in opposing directions in 42 cases; 11 ASPs were exacerbated by IF; 1 ASP could not be evaluated because Cohen’s d effect sizes could not be computed due to a 0 value in the denominator. Most of the 42 ASPs countered by IF showed a significant main effect of diet, but no significant interaction between diet and age (Fig. 5c). Further analysis of the ASPs featuring a significant interaction term (considering the directionality of change) indicated that ca. 13.7% of all ASPs identified correspond to ASPs counteracted by IF in ways consistent with the “rate effect model” or “combined rate/baseline effect model” introduced in Fig. 1b. Based on a significant genotype main effect, but a lack of an interaction, ca. 27.5% corresponded to ASPs consistent with the “baseline effect model” shown in Fig. 1b. The remaining ASPs were not affected (ca. 47.1%), accentuated (ca. 10.8%) by IF or could not be evaluated (ca. 1%). Most of the IF-attenuated ASPs examined had an age at first detectable departure from the young adult baseline of 8 months (Fig. 5c), implying that corresponding IF effects on ASPs in our young group are independent of age-related change in those ASPs.

Correlation analyses of diet effect sizes in old (Fig. 5d) and young (Fig. 5e) mice vs. effect sizes of age showed a modest inverse correlation, which is consistent with an antagonistic relationship between IF and aging.

We examined whether diet effects are similar across age groups within the category of ASPs countered by IF (n=42 phenotypes). Correlation analyses of IF effect sizes in young vs. old mice showed a significant positive relationship for ASPs antagonized by diet (Fig. 5g; R=0.60, p=3.16E-05), indicating that ASPs in young and old animals were affected by IF in similar ways. The slope of the regression line was 0.64 ± 0.14 (95% CI: 0.36, 0.92; p=0.0119), indicating that effects overall did not tend to be larger in old mice than in young mice. Similar results were obtained using intraclass correlation analyses (Fig. 5g; ICC=0.60, p=9.86E-06) which reflect not only the degree of correlation but also the agreement between measures in the young and old group. Statistical comparison of genotype effect sizes in young mice vs. effect sizes in old mice showed that in 31 out of 42 cases effect sizes in young and old mice were not significantly different (Fig. 5g; **Extended Data Table 11**; **Supplementary Data 9**). For instance, in agreement with previously published data ^16, 39, 50, 51^, advanced age was associated with decreased exploratory locomotor activity in a novel environment and this aging-associated phenotype was antagonized by IF (Fig. 5c). We identified 11 phenotypes with significantly larger effect sizes in old mice than in young mice (Fig. 5g; **Extended Data Table 11**; **Supplementary Data 9**), for example average respiratory exchange rate or NKT cell count. Based on the analysis of genotype effect sizes in young vs. old mice, ca. 10.8% of ASPs were countered by IF in ways consistent with either the “rate effect model” or “combined rate/baseline effect model” introduced in Fig. 1b (i.e., larger effect in old than in young). Ca. 30.4% of all ASPs were countered in ways consistent with the “baseline effect model” shown in Fig. 1b (i.e., effect in old not larger than in young). As mentioned above, the remaining ASPs were either not affected (ca. 47.1%), accentuated (ca. 10.8%) or could not be evaluated (1%).

Finally, we wanted to address to what extent the phenotypes used in our analyses are potentially interrelated. To address this, we performed hierarchical clustering on the phenotypic data from the young control groups for each of our three intervention studies (**Extended Data Fig. 9-11; Supplementary Data 10-12**). These analyses revealed the expected consistently low distances between phenotypes known to be related, such as e.g. peripheral blood hemoglobin concentration and hematocrit. However, they also show relatively large distances between many of the phenotypes, suggesting that much of the variation in the data would be lost if our analyses were restricted to a small subset of phenotypes only. Analyses of intervention influences on clusters (based on different cluster definitions) are summarized in **Supplementary Data 10-12**.

For all PAAIs assessed in the present study, pro-longevity effects have been demonstrated previously ^10, 26, 29^. We have shown IF-induced lifespan extension in a prior study ^10^ that was carried out side-by-side with the collection of the aging data on which the current analyses are based. Although our present experiments were not designed to ascertain pro-longevity effects (which would have required aging substantially larger groups of animals over longer time periods), provisional survival estimates based on animals aged in our facility are not inconsistent with previously reported pro-longevity effects of the *Ghrhr^lit/lit^* and *mTOR^KI/KI^* genotype, respectively (**Extended Data Fig. 6**). Moreover, comprehensive macropathological and histopathological analyses revealed significantly reduced tumor burden in *mTOR^KI/KI^* and *Ghrhr^lit/lit^* mutants relative to their wildtype littermate controls (**Extended Data Fig. 6**), which is consistent with earlier findings in *mTOR^KI/KI^* ^26^ and dwarf mice ^52, 53^, respectively. Neoplastic disease is a major factor in limiting natural lifespan in C57BL/6J animals as well as other stocks of mice ^5–10^.

As outlined above, aging and neoplastic disease often are overlapping conditions. We deliberately made no attempt to pre-select animals (for inclusion in our study) based on their apparent fitness or health status as this could have complicated the interpretation of our findings. For instance, if a longevity intervention does not improve age-sensitive phenotypes in a pre-selected set of healthy mice, this observation is difficult to interpret: Possibly, the intervention has truly no effects on age-sensitive phenotypes but, alternatively, differential inclusion could confound this analysis (e.g. controls with the poorest aging outcomes may have been excluded from the analysis and, hence, cannot be analyzed, while mice with poor - but not sufficiently poor to lead to exclusion - aging outcomes may still be existing in the intervention group, thereby resulting in a biased estimate of aging outcomes across these two populations). Nonetheless, in order to address whether neoplastic disease could have influenced some of our parameter estimates in aging mice, we subjected all animals to a macropathological assessment after completion of phenotypic analyses. Repeating the analyses outlined above on the tumor-free (i.e., free of macropathologically detectable tumors) set of mice revealed qualitatively similar results compared to the entire set of animals (**Extended Data Fig. 13-16**; **Supplementary Data 1**,**6**,**7**,**9**), suggesting that key observations of our study also hold up when considering only aged animals that are free of detectable neoplastic disease.

Altogether, our deep phenotyping analyses based on three different central longevity interventions revealed that intervention-sensitive ASPs are in many cases influenced in age-independent ways, with similar effects in aged mice and in animals younger than the age of onset of change in the corresponding phenotypes. These observations provide support for the view that age-dependent phenotypic change in these cases is not broadly slowed by these interventions. Rather, ASPs tend to alter the point of departure under these interventions; the progression of aging remains unaltered in these cases. We also identified some phenotypes that were predominantly influenced by PAAIs in old mice (with more limited or no clear effects in young mice). These cases represent phenotypes in which the progression of age-related change appears to be modified by PAAIs, consistent with a slowed rate of aging.

## Discussion

We herein defined aging trajectories of hundreds of phenotypes and thousands of molecular markers across the lifespan of male C57BL/6J mice. Our newly established atlas revealed that most age-sensitive phenotypes (ASPs) showed relative stability in young adulthood (between 3 and 5 months). Moreover, age-dependent changes in ASPs mostly began to be detectable in the second year of life in our dataset. These data serve as a critical resource for the proper interpretation of the nature of anti-aging effects induced by genetic, pharmacological or dietary interventions in mice.

We also analyzed central genetic and environmental longevity regulators (putative anti-aging interventions; PAAIs) for their mechanistic influences on hundreds of phenotypes in young and old cohorts of animals. Integration of these data with our aging trajectory dataset revealed that many intervention effects were clearly measurable not only in the old but also, with similar effect sizes, in the young cohorts of mice, at an age long before age-dependent changes in ASPs began to be detectable. Accordingly, these PAAI effects cannot be taken as evidence that the PAAIs slowed aging (age-dependent change). These observations are consistent with data we obtained in humans that also showed age-independent effects on age-sensitive phenotypes of *GHRHR* and *MTOR* genetic variants. In addition to ASPs that were influenced in age-independent ways, we also identified subsets of ASPs that were predominantly affected in the old cohort of animals, suggesting that PAAIs may potentially target age-dependent changes in these traits. Hence, our dataset allowed us to isolate different modes of actions (age-independent vs. age-dependent influences) of PAAIs acting on different ASPs.

Prevailing molecular damage theories of aging posit that aging is fundamentally caused by the age-dependent accumulation of molecular damage linked to progressive telomere shortening, accumulation of misfolded proteins, genomic instability, epigenetic changes, increased numbers of senescent cells, metabolic dysfunction, progressive and irreversible changes of the extracellular matrix, etc. ^4^. However, one current limitation of these concepts of the biology of aging is that they are largely based on lifespan data or on analyses of aging traits more limited in scope than the present study. There is also in particular a shortage of studies in mammalian models and of research that considers the controls we built in the present work. Our study shows that the PAAIs we examined -that are concerned with some of the very core mechanisms proposed to be involved in aging ^4^ - did often not seem to work through targeting age-dependent change (Fig. 6). This is not to say that we did not observe individual anti-aging effects that were consistent with a slowed aging rate; parameters that followed this pattern did, however, represent the minority of cases of anti-aging influences observed in the present study. We were able to come to this conclusion because we had included young treated (mutant, fasted) groups in our study design and determined the age at which phenotypic change began to be detectable for the parameters examined. Had we not done this, we would have substantially overestimated PAAI effects on the progression of aging. We recommend that comprehensive phenotyping, including the controls built in our study, should be adopted in future work investigating PAAIs, since this facilitates the proper interpretation of the mechanistic mode by which PAAIs influence biological aging.

**Figure 6:**
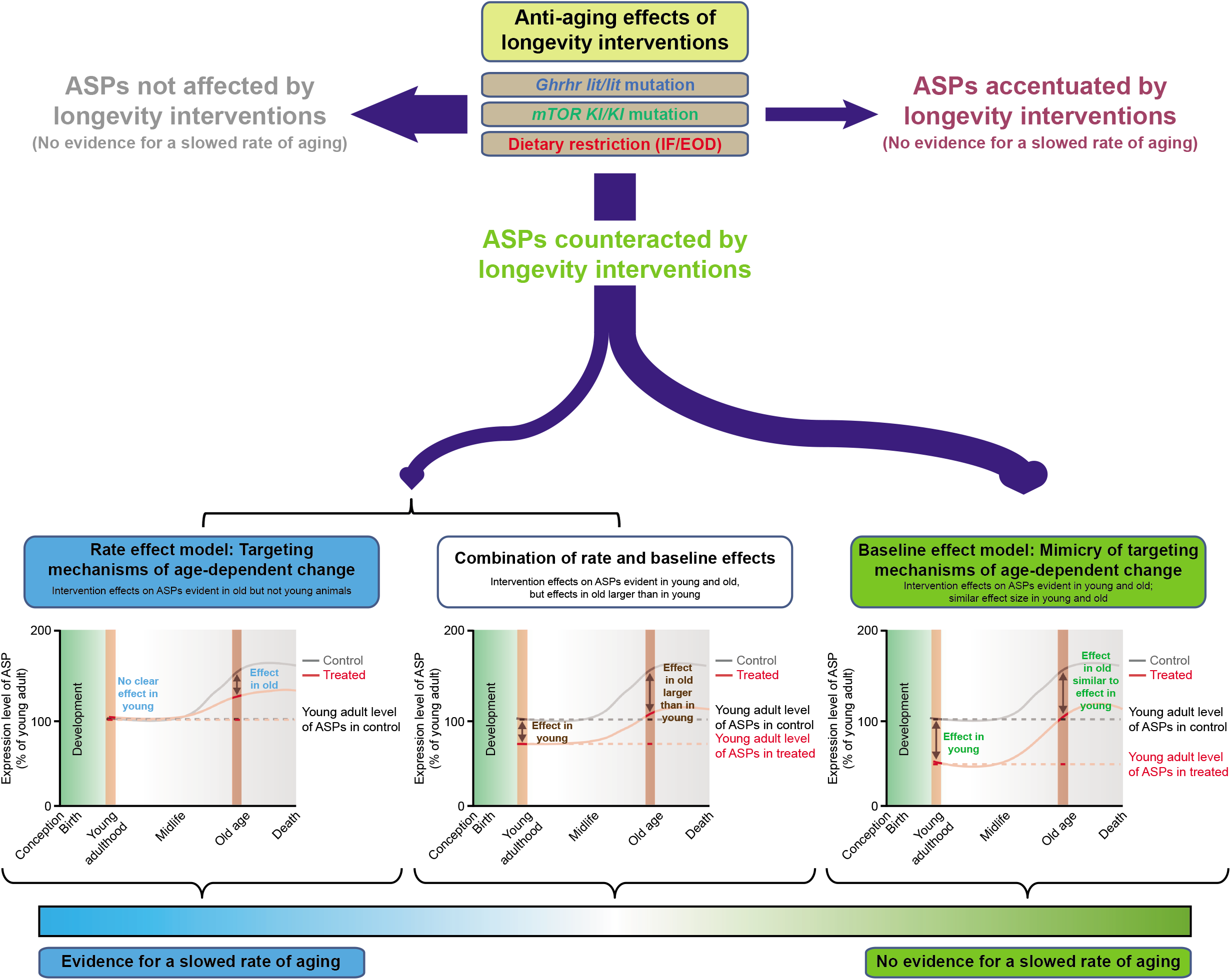
‘Anti-aging’ effects were frequently age-independent in nature. The schematic illustrates major scenarios by which PAAIs could influence aging phenotypes. First, interventions could have no measurable effect on a set of phenotypes or even accentuate age-dependent phenotypic change. ASPs countered by an intervention could be influenced in ways consistent with a targeting of the mechanisms underlying age-dependent phenotypic change: In this case, PAAI effects should become apparent only after the onset of aging-associated phenotypic change, but not at younger ages (rate effect). PAAI effects at a young age (prior to the age when age-dependent phenotypic change becomes first detectable) indicate that it is not the age-dependent change that is being targeted (baseline effect). Although our studies revealed examples of both rate and baseline effects, many ‘anti-aging’ effects fell into the latter category (age-independent effects that do not provide evidence for a slowed aging pace). Ignoring this distinction would lead to a substantial overestimation of the extent by which PAAIs slow the aging process.

In fact, the molecular and cellular mechanisms underlying aging-associated phenotypic changes, examined in the present study, are currently still poorly defined; they are likely complex and may vary from phenotype to phenotype ^15, 54^. Importantly, our approach does not require knowledge of the mechanisms underlying age-dependent change. In the absence of this knowledge, using the approach outlined in this paper, we are still able to address whether PAAIs may act by targeting age-dependent phenotypic change. It will be an important challenge for future research to define the underlying mechanisms. Our findings also suggest that a reexamination of the ‘hallmarks of aging’ processes ^4^, using large-scale phenotyping with the controls outlined in this paper, is warranted to address whether these processes indeed broadly regulate aging or are primarily the ‘hallmarks of lifespan’ with more limited roles in aging.

Our observations are consistent with early considerations by Richardson & Carter who noted that, among a handful of age-sensitive phenotypes responsive to caloric restriction (CR), CR effects were seen in young animals as well, indicating that in these cases CR shifted the level of the process and did not affect the rate of age-dependent change ^19^. Our deep phenotyping approach places this notion on a solid foundation (we analyzed hundreds of phenotypes) and extends this early consideration to IF-based models of dietary restriction as well as central genetic models of longevity, targeting mTOR and growth hormone signaling pathways. Our findings, therefore, raise the possibility that the age-independent nature of ‘anti-aging’ effects may be common among longevity interventions. This would imply that many ‘anti-aging’ effects are unlikely to arise from targeting the causal factors driving aging.

Our findings are in agreement with the notion that genetic effects tend not to be strictly age-specific but mostly affect the organism across its lifespan ^54^. Accordingly, given that our PAAIs often had effects in young individuals, which were frequently of similar effect size as those in old subjects, relatively shorter-term exposure to treatment may to some extent be sufficient to induce sizeable PAAI effects at a young age. Therefore, an important next step is to address whether PAAIs can also induce sizeable treatment effects when old individuals with established phenotypic change are subjected to short-term interventions. This may substantially simplify the development of therapeutics because of shortened treatment periods and the possibility that treatment may, at least in some cases, come with therapeutic benefit even after the onset of age-dependent change. These treatments would be considered symptomatic, not causal, in nature but may still provide valuable alleviation of a subset of aging phenotypes.

Our results also imply that future research should not only identify the molecular and cellular mechanisms underlying age-dependent change in ASPs but also compare them to those by which PAAIs affect ASPs in young animals (i.e., prior to the onset of age-dependent change). For instance, we have found here that age-related alteration in nociceptive function was ameliorated by *Ghrhr^lit/lit^* genotype (Fig. 3c). The *Ghrhr*^lit^ allele increased thermal sensitivity in old mice as well as in young animals (Fig. 3c). The age-dependent loss of thermal sensitivity has been previously linked to a reduced expression, on the protein but not the mRNA level, of Trpv1 in dorsal root ganglion cells in 15-month and 2-year old mice compared to 6-week old controls ^55^. Growth hormone deficiency has been shown to induce thermal hypersensitivity in neonatal mice and this phenotype has been suggested to be linked to a transcriptional upregulation of *Igfr1*, *P2×3*, *Piezo2*, *Trpv1* and *P2y1* in the neonatal mutants ^56^. These observations confirm that thermal hypersensitivity is influenced by growth hormone deficiency long before age-dependent changes in this ASP develop. They also suggest that aging and growth hormone deficiency act on this ASP via separable mechanisms. Given that the ASPs examined in the present study tend to be complex phenotypes, being shaped through a plethora of molecular and cellular regulators, we put forth the testable prediction that, based on probability, it is unlikely that there is a large common base between the mechanisms underlying PAAI effects on ASPs in young mice and those driving age-dependent change of the same phenotypes.

Our study has some limitations. We did not use a longitudinal study design to infer aging trajectories and intervention effects. Rather, we employed a cross-sectional study design that compared different sets of mice to extract age and intervention effects. While longitudinal analyses have their strengths (i.e., being able to take repeated measurements on the same animal, at least for the sets of phenotypes where repeated measurements are possible), they are certainly not without complications and have their own sets of limitations: An important complication of performing assays within longitudinal designs are order effects, referring to the phenomenon that having been tested once or more times has effects on the outcome of subsequent tests (e.g., in behavioral assays). Another important issue inextricably linked to longitudinal designs is that measurements cannot be taken at the same time but instead require comparison of data collected at different points in time (which would be up to 2 years apart in our study). Hence, although there may be exceptions depending on what parameters one specifically considers, longitudinal data may not necessarily generate more robust estimates of aging trajectories in mouse populations than population estimates derived from cross-sectional data collected at the same time, by the same person and under the same well-controlled conditions. Please also note that a considerable number of parameters we measured was collected in the context of terminal procedures, precluding repeated data collection from the same animal. In line with all of these considerations, it is common practice to infer aging trajectories from studies using cross-sectional designs ^57–62^.

One goal of our analyses was to estimate, using pairwise comparisons against the 3-month old group, when mouse phenotypes show first measurable departures from our young adult baseline (i.e., from the 3-month old group). Note that the sensitivity to detect differences between these groups (i.e., the limit of detection) depends on sample size. Increasing sample size could almost arbitrarily lower the limit of detection and potentially reveal additional smaller-sized group differences if they exist.

We identified a number of ASPs that appeared to be affected by a combination of a baseline and a rate effect (i.e. ASPs were affected in both young and old mice; the effect size was larger in old mice than in young mice). One possible interpretation is that age-dependent phenotypic change is in fact slowed in these mice (additionally to an age-independent effect on these ASPs). However, an alternative explanation is that shorter intervention exposure times in young mice account for smaller effects relative to old mice that were exposed to the intervention for a longer period. The shorter-term exposure to intermittent fasting in young mice (IF was initiated 4 weeks prior to the commencement of analyses), for instance, may have led to similar but smaller effects on some ASPs than the more long-term exposure (animals were on IF for 19 months prior to starting the analysis) in our aged cohort of animals. Future experiments should help distinguish between these scenarios by varying exposure time during a period in life where ASPs are relatively stable.

In our intervention studies, we examine cross-sectionally age-dependent changes in phenotypes between two time points (young adult vs. ca. 20 months old). As a consequence, our rate of change estimates refer to an overall change across this time period. A PAAI could potentially not only alter the rate of change but also modify the onset of age-dependent change (e.g., an intervention could lead to an earlier onset of change from baseline associated with a slower rate of change). We cannot address this possibility with our present dataset because that would require a study design with a number of age groups in between the young adult and aged group. Note though that rate of change estimates based on our two time points are well suited to measure the overall change on a population level that accumulates between young adulthood and 20 months of age.

We chose a large set of parameters for our analyses in mice to cover a broad range of age-related changes across numerous physiological systems and tissue contexts, spanning across multiple levels of biological organization (molecular, cellular, tissue, organismal). Our prior analyses established that a large and diverse number of ASPs can be captured using this approach ^10, 16^. However, despite the comprehensive nature of our analysis it is important to note that our conclusions are based on the specific sets of parameters included in our assessment of age-related changes in mice. It will be important for the field to complement our analyses with assessments of additional measures and extend our observations beyond the one genetic background (C57BL/6J) and sex (male) examined here.

In all our intervention studies, the aged groups of mice were ca. 20 months old when our analyses were started. Although this leaves unexamined older age groups with potentially additional aging-associated changes, we have chosen to examine 20-month old mice (and not older ones) to avoid interpretational issues that may arise from differential survival. For instance, if a longevity intervention does not improve age-sensitive phenotypes in 30-month old mice, this observation is difficult to interpret: Possibly, the intervention has truly no effects on age-sensitive phenotypes but, alternatively, differential survival could confound this analysis (e.g. controls with the poorest aging outcomes may have been eliminated from the population and, hence, cannot be analyzed, while mice with poor, but not yet detrimental aging outcome may still be existing in the intervention group, thereby resulting in a biased estimate of aging outcomes across these two populations). Assessing aged mice at 20 months largely eliminated differential survival as a confounding factor from our analysis (at 20 months, there has not yet been appreciable population attrition in the controls; see Fig. 1a).

The determination of when aging begins to manifest is ultimately a matter of one’s viewpoint and depends on the parameters one chooses to assess the consequences of aging ^63^. We applied a set of parameters that are relatively stable over at least some months in younger adult animals. If one intends to assess PAAIs in the context of parameters other than the ones used in the present study, it may be necessary to establish these parameters’ individual lifetime profiles and to adjust the age of the young treated reference group, so that the PAAI can be restricted to a period prior to the onset of detectable age-dependent change.

Our conclusions are based on the specific set of PAAIs that were investigated in our current study (loss of function of *Ghrhr*, *mTOR*; intermittent fasting-based version of dietary restriction; see also **Supplementary Discussion**). As outlined above, our longevity mouse lines were chosen to represent important and central genetic/environmental lifespan-extending interventions established by prior research. mTOR signaling and growth hormone signaling are not only among the most well-established pathways in lifespan regulation, they also feature many links to cellular processes thought to generally play important roles in aging, such as proteostasis, nutrient sensing, inflammation and others ^4, 41, 64^. Similarly, dietary restriction regimens are thought to broadly influence a range of cellular processes linked to aging ^4, 49^. Thus, our findings have some generalizability beyond these three interventions. Nevertheless, how manipulations of other longevity-associated pathways interact with age-dependent change needs to be addressed in further studies examining additional genetic mutants and/or environmental manipulations.

A number of studies have applied multidimensional analytical approaches to measure organismal changes accompanying the aging process ^65–69^. An important feature of our study is that it uses multidimensional phenotypic data, covering multiple levels of complexity from molecular markers to complex physiological and tissue functions, towards testing for a possible modulatory effect of genetic and environmental intervention effects on the aging process. Unlike studies in invertebrate models, our approach in mice captures mammalian physiology. In comparison to studies in humans, the experiments in mice facilitate causal and invasive experimentation (such as the genetic and fasting studies as well as the collection of many invasively obtained phenotypes that would be impossible to get in humans); they also facilitate the lifelong (or almost lifelong) exposure to experimental regimes (such as genetic and environmental manipulations that cover much of the organismal lifespan of a mouse).

Besides phenotypic analyses, we carried out transcriptome studies to test for age-dependent changes in tissue level gene expression. In addition to the full set of differentially expressed genes, we provide results derived from pathway analyses that were performed to detect whether specific pathways were enriched among the sets of differentially expressed genes. Note that the results for some of these annotated processes may seem counterintuitive, although they are expected. For instance, based on gene expression changes associated with advanced age, Ingenuity pathway analysis (IPA) derived predictions of reduced “organismal death” and “morbidity/mortality” in old mice. While these predictions appear to be inconsistent with the actually increased mortality rate associated with aging, they are expected ^70^ and are based on gene expression changes well known to be associated with aging (increased expression of inflammation-related genes in the context of aging ^71^) that lead IPA to predict a downregulation of the processes “organismal death” and “morbidity/mortality”. We also note that some of the IPA categories (such as many of the canonical pathways) contain a small number of member genes and/or may intersect with a small number of differentially expressed genes which can render these categories susceptible to variable results. These findings should not be over-interpreted without further study and corroboration.

Although there are few studies employing multi-point profiling of gene expression changes in rodent models, recent reports based on various rat and mouse tissues ^59, 72^, indicated that most of the differentially expressed genes were first detectable during middle-ages (12-15 months) with progressive gradual changes till advanced age. However, there were notable differences across tissues, with the spleen for instance exhibiting more robust age-associated transcriptomic changes and in contrast, the brain being more resistant to aging ^59^. Moreover, pathway analysis of the differentially expressed genes associated with aging in these studies suggested the presence of both organ specific and global molecular signatures ^59, 72^. These findings are consistent with our observations from spleen and brain bulk RNA-seq datasets (Fig. 2g–j, **Extended Data Fig. 3**).

In the present study, many age-sensitive phenotypes did not provide support for a slowing of age-dependent change in the PAAI groups, indicating that PAAIs did not exert their effects by inhibiting the accumulation of aging-associated damage (at least with respect to the ASPs examined). There are interesting parallels between these results and earlier observations: Dietary restriction (DR) in *Drosophila melanogaster* was found to reduce mortality risk entirely via short-term effects; flies that were transiently food-restricted and then switched to free access to food quickly adopted the mortality risk of flies that had free access to food throughout their lives ^73^, indicating that DR had no sustained influence on mortality risk (as one might expect if DR were to reduce mortality risk by slowing the accumulation of aging-associated damage).

Analyses of survival curves in pro-longevity mouse models indicated that the rate at which mortality risk increases with advancing age (captured in the Gompertz function parameter “G”) was unaffected ^74^; instead, pro-longevity interventions in mice shifted the age at which mortality risk started to increase (captured in the Gompertz function parameter “A”) to older ages, without changing the rate of increase thereafter. Although it remains unclear how specifically these findings relate to ours, this observation has been interpreted as evidence that changes in the “rate of aging” (based on a lack of effect on the parameter “G”) may not underlie the pro-longevity influences in mice, but rather changes in “baseline vulnerability” to adverse effects of disease and environmental factors (based on an effect on the parameter “A”) ^74^. In fact, resistance to the development of lethal neoplastic disease could represent such a change in “baseline vulnerability” in long-lived mutant mouse lines.

It is important to complement lifespan analyses with additional measures capturing age-dependent change, if one wishes to make statements about aging. This is particularly relevant in cases where it is unclear how well lifespan reflects broad changes across a range of physiological systems. In mice, for instance, lifespan is well known to be limited by the development of lethal neoplastic disease, indicating that it is determined by a rather narrow set of pathologies and hence cannot reflect aging-associated changes across a broader set of physiological systems ^5–10^. It will be an important task to further define the sets of biological processes that limit lifespan in other organisms, specifically in those that heavily informed the biology of aging via studies of lifespan (such as *C. elegans* and *D. melanogaster*) ^75–78^.

In the present work, we used age-dependent phenotypic change across a range of molecular, cellular, physiological and pathological markers as a proxy for biological aging. This does not imply that all aging-associated change is necessarily adverse. In fact, it has been pointed out that aging-associated alterations can have adaptive and beneficial effects for the aged organism ^79, 80^. Future work will need to address which of the phenotypes employed in our present study design may serve adaptive purposes vs. may have detrimental effects for aging mice.

A unique strength of the current study is that we included young adult treated (mutant/fasted) groups as well as an assessment of onset of measurable age-dependent phenotypic change in our comprehensive large-scale analyses of phenotypic and molecular alterations. This permitted us to separate age-independent PAAI effects from interactions of PAAI with age-dependent phenotypic change. Any study not considering these controls is bound to substantially overestimate the extent by which PAAIs slow the aging process.

Aging is a multi-faceted process that transforms young adult into aged organisms. We believe that the identification of molecular regulators influencing aging will require approaches that measure many aspects of this organismal transformation directly, rather than relying on single or a small number of proxy markers. The isolated focus on lifespan, for instance, bears the risk of bias by putting center stage the subset of aging processes directly linked to lifespan (such as specific pathologies), but with potentially limited relevance for other facets of aging that do not *per se* determine the end of life. Much of what we currently think we know about molecular regulators of aging (summarized, e.g., in the “hallmarks of aging” ^4^) has been derived from studies utilizing such proxy markers of aging. We therefore look forward to seeing more studies attempting to validate these regulators in the context of multidimensional phenotypic studies.

In conclusion, the PAAIs examined (i.e. *mTOR* loss of function, *Ghrhr* loss of function, intermittent fasting-based version of dietary restriction) often influenced age-sensitive traits in a direct way and not by slowing age-dependent change. These findings have important implications regarding the extent to which the aging process can be modulated. Having said that, we also identified ASPs predominantly influenced in the old groups of mice, indicating that, on a subset of ASPs, PAAIs exerted their effects by slowing the rate of age-dependent change. Our analytical approach provides a valuable resource as well as an important framework for future research aimed at parsing how genetic and environmental factors interact with the mammalian aging process.

## Abbreviations

ASP: age-sensitive phenotype
PAAI: putative anti-aging intervention

## Acknowledgements

This work was supported by a grant from the Helmholtz Future Topic AMPro (Aging and Metabolic Programming), the German Federal Ministry of Education and Research (Infrafrontier grant 01KX1012), the German Center for Neurodegenerative Diseases (DZNE) and the German Center for Diabetes Research (DZD). Immune phenotyping in the Rhineland Study is partially supported by the Deutsche Forschungsgemeinschaft (DFG, German Research Foundation) under Germany’s Excellence Strategy – EXC2151 – 390873048. DL was supported by a scholarship from China Scholarship Council. We acknowledge use of the Genotype-Tissue Expression (GTEx) resource supported by the Common Fund of the Office of the Director of the National Institutes of Health, and by NCI, NHGRI, NHLBI, NIDA, NIMH, and NINDS. The data used for the analyses described in this manuscript were obtained from: the GTEx Portal (www.gtexportal.org, accessed on 05/30/20 and 07/15/20). We thank Sach Mukherjee and Gerhard Ehninger for valuable discussions and feedback on an earlier version of the manuscript. We acknowledge Kristian Händler and Joachim L. Schultze for providing access to the PRECISE genomics platform at the DZNE. Figures were created with BioRender.com and/or by adapting from BioRender.com templates.

## Author contributions

D.E. conceived and initiated the study; K.X., H.F., V.G-D. and D.E. planned and prepared the study; K.X., E.S., D.L., A.A., J.A.A.-P., O.V.A., L.B., P.S-B., J.C-W.,Y.C., Y.D., A.C.E., L.G., C.G., R.G., S.M.H., T.K-R., D.L., P.M-K., L.L.N., C.O., B.L.P., B.R., J.Ro., K.S., N.S., A.S-M., I.T. and D.E. performed the experiments; K.X. E.S., D.L., A.A., J.A.A.-P., O.V.A., L.B., P.S-B., J.C-W.,Y.C., L.G., R.G., S.M.H., M.K., T.K-R., D.L., P.M-K., M.A.O., B.R., J.Ro., J.Ru., N.S., A.S-M., I.T. and D.E. analyzed the data; K.X., S.L., C.S. and D.E. did project coordination; D.B., D.H.B., J.G., M.K., T.K., B.A.M., P.S., C.S.-W., M.W., E.W., W.W., V.G.-D., M.M.B.B., H.F., M.H.A. and D.E. provided oversight and resources; K.X., E.S., D.L., A.A., B.L.P., D.B. and D.E. wrote the manuscript.

## Material and methods

### Mice, age-sensitive parameters and interventions

We used wild-type C57BL/6J mice to determine the trajectories of ASPs in mice. All the analyses described in the paper were restricted to male mice only. Animals for the quantification of ASP trajectories were assessed at 3, 5, 8, 14, 20 and 26 months of age: We had one cohort of mice for all deep phenotyping as well as RNA-seq analyses (data shown in **Supplementary Data 1-4**) and a separate cohort from which we harvested tissues for the molecular analyses presented in **Supplementary Data 5**. Three major longevity mouse lines were included in our analysis to address to what extent the corresponding interventions may or may not delay aging trajectories: These featured either a loss-of-function mutation in the *Ghrhr* gene (*Ghrhr^lit^* mutation), were engineered to carry a hypomorphic *mTOR* mutant allele or were subjected to almost lifelong intermittent fasting. *Ghrhr^lit^* mutants ^28, 29^, also known as B6-little, were obtained from the Jackson Laboratory (stock no. 000533) (data summarized in **Supplementary Data 6 and 10**). Founder animals of the hypomorphic mTOR mouse line ^25^, bearing a knock-in sequence replacing exon 12 within the murine mTOR gene, were kindly provided by Dr. Wendy DuBois (Laboratory of Cancer Biology and Genetics, Center for Cancer Research, National Cancer Institute, Bethesda, MD, US) (data summarized in **Supplementary Data 7, 8 and 11**). After arrival at our facility, mice carrying the hypomorphic mTOR allele were crossed with wild-type C57BL/6J mice for five generations prior to use of the mice in the current analyses. The intermittent fasting (IF) cohort has been previously described and was generated using an every-other-day feeding paradigm in group-housed male wild-type C57BL/6J mice ^10^ (current analysis summarized in **Supplementary Data 9 and 12**). In brief, intermittent fasting (IF) animals were subjected to alternating 24h-cycles of free access to food (Altromin 1314 standard rodent chow) and complete food deprivation, starting at the age of 8 weeks and continued throughout life of the animals. The Altromin 1314 chow came in solid pellets. Pilot experiments showed that mice did not crumble these pellets. Accordingly, removing the pellets on the restriction days was sufficient to fully deprive the animals of food (no cage change required). IF mice were compared to controls with *ad libitum* (AL) access to food throughout the course of the study. To evaluate whether these interventions’ effects on ASPs are primarily due to either altering the rate of age-dependent change in ASPs or due to age-independent effects on ASPs, we analyzed aged mutant/fasted animals/wildtype littermate controls (if not stated otherwise, ∼20 months old at the commencement of the deep phenotyping analysis) side-by-side with young mutant/fasted mice/wildtype littermate controls (∼3 months old at the commencement of the deep phenotyping analysis).

Our deep phenotyping approach covered a wide range of analyses, including assessments of anatomical, physiological, metabolic, neuropsychiatric, cardiovascular, immunological, sensory, molecular, cellular and histopathological parameters. The following analyses were carried out in the order listed and were completed within a period of 11 weeks, if not stated otherwise ^81, 82^: modified SHIRPA (week 1), open field (week 1), grip strength (week 1), rotarod (week 2), acoustic startle response and pre-pulse inhibition (week 2), clinical chemistry after fasting (week 3), hot plate test (week 4), body surface temperature (week 4), transepidermal water loss (week 4), indirect calorimetry (week 5), body composition analysis (week 5), glucose tolerance test (week 6), electrocardiography (week 7), echocardiography (week 7), Scheimpflug imaging (week 8), optical coherence tomography (week 8), laser interference biometry (week 8), virtual drum vision test (week 8), auditory brain stem response (week 9), bone densitometry (week 9), clinical chemistry (week 11), hematology (week 11), FACS-based analysis of blood leukocyte populations (week 11), immunoglobulins and plasma biomarkers (week 11), lymphocyte proliferation assay (week 11) and pathology (week 11). In general, animals, within their home cage, were allowed to habituate to the test room for a period of at least 15 min prior to the start of experimental procedures, if not stated otherwise. In all cases, experiments were carried out according to the IMPReSS (International Mouse Phenotyping Resource of Standardised Screens) workflow in compliance with the International Mouse Phenotyping Consortium (IMPC) (https://www.mousephenotype.org/) and the metadata was recorded. Analyses and procedures were always balanced across experimental groups. The number of animals examined depended on the specific experiment and assay used and is indicated in the data provided (for details, see **Supplementary Data 1**, **5-7, 9**). Of note, assessments in IF mice were performed after a feeding day to avoid measuring acute hunger effects.

All mice were housed in individually ventilated cages (IVC) under specific pathogen-free conditions (according to FELASA guidelines) in groups of 2-5 mice. All mice were fed with Altromin 1314 standard rodent chow (composition: 5.1% fat (equivalent to 14% of total metabolizable energy), 22.5% protein (equivalent to 27% of total metabolizable energy) and 40.4% carbohydrates (equivalent to 59% of total metabolizable energy)). Husbandry conditions included a constant temperature of 22 °C, a 12h:12h light/dark cycle as well as *ad libitum* access to food (except for IF animals as described above) and water. Local and federal regulations regarding animal welfare were followed. In accordance with the German Animal Welfare Act, the present study was approved by the “Landesamt für Natur, Umwelt und Verbraucherschutz Nordrhein-Westfalen” (Recklinghausen, Germany) as well as the “Regierung von Oberbayern” (Munich, Germany).

### Modified SHIRPA

The modified SHIRPA protocol was designed as a rapid semi-quantitative screen to detect phenotypic anomalies in mice ^83^. Our SHIRPA test battery started with the inspection-based assessment of the animals’ general appearance as well as their undisturbed behavior as observed in a transparent glass cylinder (11 cm in diameter). Mice were then transferred to a transparent Perspex box (420 mm x 260 mm x 180 mm), marked with a grid on the floor, for an assessment of general neurological status. A set of 17 tests were performed including assessments of the acoustic startle reflex, biting behavior, body position, contact righting reflex, defecation, gait, head bobbing, limb grasping, locomotor activity, pinna reflex, tail elevation, touch escape, transfer arousal, tremor, trunk curl, urination and vocalization.

### Open field

To measure general locomotor activity in a novel environment, we subjected animals to an open field assay using a transparent and infrared light-permeable acrylic test box (45.5 cm x 45.5 cm x 39 cm inner dimensions) equipped with evenly spaced infrared light beams along the x- and y-axis and a rearing indicator covering the z-axis (ActiMot, TSE, Bad Homburg, Germany). Illumination in the center of the test box was set to ∼200 lux; light intensity in the corners was ∼150 lux. Animals were transferred to an area immediately adjacent to the test room where they were left undisturbed for a period of 30 min prior to commencement of open field analysis. In the open field assay, mice were allowed to freely explore the novel environment (the open field box) for 20 min. Our analysis included the following parameters: distance traveled within the first 5 min, total distance traveled, number of rearings within the first 5 min, total number of rearings, percentage of distance traveled in the center of the box within the first 5 min, percentage of total distance traveled in the center of the box, percentage of the first 5 min spent in the center of the box, percentage of total time spent in the center of the box, average velocity.

### Grip strength

Grip strength was measured using a grip strength meter system (Bioseb, Pinellas Park, FL, US). The animal, held by its tail, was allowed to grab a metal grid with either two or four paws and was then pulled back horizontally. The maximum force applied to the grid, just prior to the animal losing grip, was recorded as the peak tension by a force sensor. Three trials were given to each mouse over the course of 1 min. Two-paw/four-paw grip strength was calculated by averaging the animals’ performance over three consecutive trials.

### Rotarod

Motor coordination and balance were evaluated on an accelerating rotarod (Bioseb, Pinellas Park, FL, US) ^84^. After the mouse was placed on the apparatus, the rod was subjected to linear acceleration from 4 to 40 rpm over the course of the 5 min test period. The trial was terminated once the animal fell off the rod, displayed passive cycling or the 5 min time period had elapsed, whichever came first. Latency to fall was averaged across three trials given to each mouse with inter-trial intervals of 15 min.

### Acoustic startle response and pre-pulse inhibition

Acoustic startle reflex and pre-pulse inhibition were assessed using a startle apparatus (Med Associates, Fairfax, VT, US) equipped with four identical sound attenuating chambers (inner dimensions: 55.88 cm x 34.29 cm x 36.83 cm). Animals were left undisturbed in an area adjacent to the testing room for 30 min prior to the start of the experiment. Next, each mouse was habituated to the test compartment (a mouse restrainer) over a period of 5 min. Background noise was set to 65 dB. Bursts of white noise (40 ms in duration) were used as startle pulses. The protocol applied began with a 5 min acclimation period succeeded by five leader startle pulses at 110 dB that were excluded from analysis. Trial types for pre-pulse inhibition included four different pre-pulse intensities (67, 69, 73 and 81 dB) and generally preceded the startle pulse (110 dB) by 50 ms. Each trial type was presented 10 times in random order, organized in 10 blocks, each trial type occurring once per block.

### Hot plate test

The hot plate test was carried out using an Analgesia Meter Hot Plate apparatus (TSE, Bad Homburg, Germany). In brief, the animal was placed on a metal surface maintained at 52 ± 0.2 °C which was surrounded by a cylindric plexiglas restrainer (20 cm high, 18 cm diameter) in order to restrict movement of the animal. The trial was terminated once we observed one of three typical indications of pain (hind paw licking, hind paw shake/flutter or jumping); the mouse was then immediately removed from the metal plate. Latencies and response type (hind paw licking, hind paw shake/flutter or jumping) were recorded. The maximum duration of this test was limited to 30 s in order to avoid tissue damage.

### Body surface temperature

Body surface temperature was assessed using infrared thermovision. Specifically, we determined maximal, average and minimal body surface temperature using a FLIR A655sc camera system (FLIR Systems, Wilsonville, OR, US).

### Transepidermal water loss

Transepidermal water loss via diffusion or evaporation was assessed non-invasively with an AquaFlux AF200 evaporimeter (Biox Systems, London, UK). To quantify the amount of transepidermal water loss, the probe was placed on the skin of the mouse for a period of 60 – 90 s.

### Indirect calorimetry

Metabolic turnover was analyzed via indirect calorimetry. Mice were single-housed in individually ventilated metabolic cages (Phenomaster, TSE, Bad Homburg, Germany) at 23 °C and were maintained on a 12h:12h light dark cycle (lights on at 6 am, light off at 6 pm). Metabolic cages were constantly supplied with fresh air and changes in O_2_ and CO_2_ levels were recorded by high precision sensors in every cage. Additionally, locomotion and rearing activity were detected by infrared sensors surrounding the cages. Animals were habituated to the metabolic cages for a period of 2 hours prior to starting measurements. Recordings were generally carried out over a period of 24 hours, except for the study of IF mice in which we measured for 47 hours to accommodate metabolic analyses during both feeding and fasting days. All mice were granted free access to food and water with the exception of IF mice which were only fed during the first 24 hours. Relevant parameters determined by indirect calorimetry included minimal O_2_ consumption, average O_2_ consumption, maximal O_2_ consumption, minimal respiratory exchange rate (RER, computed as volume of CO_2_ generated (VCO_2_)/volume of O_2_ consumed (VO_2_)), average RER, maximal RER, ΔRER (maximal RER – minimal RER), minimal heat production (calculated using the formula: heat production (mW) = (4.44 + 1.43 x RER) x VO_2_ (ml/h)), average heat production, maximal heat production, cumulative food intake, cumulative water consumption, total distance traveled, cumulative number of rearings and cumulative number of fine movements. In addition, body weight was measured at the beginning and at the end of the metabolic assessments.

### Body composition analysis

Time Domain Nuclear Magnetic Resonance (TD-NMR)-based body composition analysis was carried out by placing animals into a Minispec Whole Body Composition Analyzer (Bruker, Billerica, MA, US). Measures of interest were lean mass, fat mass, and total amount of free body fluid.

### Glucose tolerance test

We carried out an intraperitoneal glucose tolerance test (IpGTT) subsequent to six hours of food deprivation. Body weight was recorded before and after the fasting period. The tip of the tail was scored and a small drop of blood was used to determine the base glucose level via an Accu-Chek Aviva glucose analyzer (Roche, Basel, Switzerland). After 2 g glucose per kg body weight was injected intraperitoneally, blood glucose levels were sequentially measured at four additional time points (15, 30, 60 and 120 min after injection).

### Electrocardiography

Non-invasive electrocardiographs (ECG) were recorded using ECGenie (Mouse Specifics, Framingham, MA, US) in conscious animals ^85^ in order to avoid effects of anaesthesia on cardiac function ^86^. Cardiac electrical activity was measured through the paws of the animal staying on a shielded acquisition platform. Intervals and amplitude from at least 15 consecutive ECGs were averaged. Heart rate was calculated from peak detections. P, Q, R, S and T waves were analyzed such that unfiltered noise or motion artifacts were excluded. The corrected QT interval (QTc) was calculated by dividing the QT interval by the square root of the preceding RR interval. QT dispersion represents inter-lead variability between QT intervals. QTc dispersion was calculated as the rate QTc dispersion. Relevant parameters included were: duration of the P wave, PR interval, QRS interval, QT interval, QT dispersion, QTc, QTc dispersion and RR interval.

### Echocardiography

We performed transthoracic echocardiography in awake animals using the Vevo 2100 Imaging System (Visual Sonics, Toronto, Canada) equipped with a 30 MHz transducer. Anatomic structure and cardiac function of the left ventricle were analyzed by dual mode imaging. Specifically, left ventricular parasternal short- and long-axis views were imaged in B-mode and left ventricular parasternal short-axis imaging was performed in M-mode at the papillary muscle level. Short-axis M-mode images derived from three consecutive heart beats were used to measure the following anatomic parameters: left ventricular end-diastolic internal diameter (LVIDd), left ventricular end-systolic internal diameter (LVIDs), diastolic septal wall thickness (IVSd), systolic septal wall thickness (IVSs), thickness of the left ventricle posterior wall during diastole (LVPWd) and thickness of the left ventricle posterior wall during systole (LVPWs). Corrected mass of the left ventricle (LV mass corr) was computed as LV mass corr = 0.8 x (1.053 x ((LVIDd + LVPWd + IVSd)^3^ – LVIDd^3^)). Additionally, we determined ejection fraction (ES), fractional shortening (FS) as well as stroke volume (SV). EF was calculated as EF% = 100 x ((LVvolD – LVvolS)/LVvolD) with LVvol = ((7.0/(2.4 + LVID) x (LVID^3^). The formula FS% = ((LVIDd – LVIDs/LVIDd) x 100 was used to calculate FS. The difference between the end-diastolic and the end-systolic blood volumes during one heartbeat was defined as SV. We also measured heart and respiration rates during echocardiographic assessment.

### Scheimpflug imaging

Optical density and morphology of lens as well as cornea were assessed in a contact-free manner using the Pentacam system (Oculus, Wetzlar, Germany). For the acquisition phase, the mouse was placed on a platform in front of the apparatus and the eye was positioned towards a vertical light source (LEDs, 475 nm). The distance between eye and the Pentacam device was adjusted automatically by the software to gain optimal focus.

### Optical coherence tomography

The posterior segment of the eye, including retina and fundus, were evaluated in anaesthetized mice using a Spectralis OCT device (Heidelberg Engineering, Heidelberg, Germany). After 1% atropine was administered to widen the pupils, a small amount of Methocel 2% (OmniVision, Puchheim, Germany) was used to carefully place a contact lens of 10 mm focal length (Roland Consult, Brandenburg an der Havel, Germany) onto the eye. For measurements, the animal was placed on an elevated platform in order to optimally position the eye in front of the transducer of the recording unit. Images were taken as described previously ^87^. We measured thickness and morphology of the retina, as well as morphology of the optical disk, fundus pigmentation and the number of main blood vessels.

### Laser interference biometry

Eye size measurements were performed using the AC Master system (Carl Zeiss Meditec, Jena, Germany). Briefly, anaesthetized mice were placed on a platform. Proper positioning of the animals was supported by light signals from six infrared LEDs arranged in a circle that must be placed in the center of the pupil. Eyes were treated with 1% atropine to ensure pupil dilation. Central measurements of axial eye length were performed as described elsewhere^88^.

### Virtual drum vision test

Visual acuity was tested using OptoMotry, a virtual optomotor system (Cerebral Mechanics, Westchester County, New York, US) as described previously ^89^. Prior to the start of the assessments, animals were placed on an elevated platform surrounded by four computer monitors. Animals were then allowed to track a virtual rotating cylinder comprised of a sine wave grating and their movements were recorded by an overhead camera. A lack of compensatory head and neck movements countering the motion of the sine wave grating indicates an inability to discern the displayed visual pattern. Rotation speed and contrast of this test was set to 12 d/s and 100%, respectively.

### Auditory brain stem response

Non-invasive assessments of hearing sensitivity were performed using an auditory brain stem response (ABR) test (Industrial Acoustics Company, North Aurora, IL, US). After anaesthetizing the animals via i.p. injection of a ketamine/xylazin mixture, mice were placed on a heated blanket (37-38 °C) and were then transferred to the acoustic chamber. Subsequently, three electrodes were placed subcutaneously (COM-electrode behind the right ear, G1-electrode on top of the skull and G2-electrode underneath the left ear). Auditory brain stem responses were induced using different acoustic stimuli, including clicks of 0.01 ms duration and beeps of a given frequency (6, 12, 18, 24 and 30 kHz; 5 ms duration, 1 ms rise/fall time). Hearing threshold levels for the respective stimuli were determined by gradually elevating sound intensity from 5 to 85 dB SPL in 5 dB steps.

### Bone densitometry

Bone densitometry was performed non-invasively using either dual-energy X-ray absorptiometry (DXA) or micro-CT imaging in anesthetized animals. DXA analyses were performed using an UltraFocus DXA system (Faxitron Bioptics, LLC). Relevant parameters examined included bone area, bone mineral content, bone mineral density and volumetric bone mineral density of the whole animal excluding the skull. Micro-CT analysis of tibiae of IF mice and their controls was processed using a SkyScan 1172 micro-CT scanner (Bruker, Billerica, MA, US). A resolution of 7.88 µm pixel was achieved at 80 kV voltage, 100 mM current and using a 0.5 mm aluminum filter. Images acquired from cross-sectional slices of the distal tibia (197 µm from 25 slices halfway between the tibia-fibula junction and the distal end of the tibia) were reconstructed by the SkyScan volumetric NRecon reconstruction software (Bruker, Billerica, MA, US) and further analysis was performed using the CT Analyser software (CTAn, v.1.15) (Bruker, Billerica, MA, US). Relevant parameters were: total tissue area, bone area, marrow area, cortical thickness and polar moment of inertia.

### Blood sampling

For blood sample collection, the animals’ retrobulbar sinus was punctured using non-heparinized glass capillaries (1 mm diameter) after inducing anesthesia with isoflurane. To assess metabolic parameters in the fasted state, blood specimens were collected in heparinized tubes from animals after an overnight 16-hour food withdrawal. Blood collected from animals in the fed state were distributed into three portions. While the larger fraction was collected in heparinized tubes, we also collected two portions in EDTA-coated tubes. To ensure homogeneous distribution of the anticoagulant, tubes were inverted five times after blood collection. Samples were then stored at room temperature for 1-2 h. Next, heparinized tubes were centrifuged for 10 min at 4 °C and 4200 g to pellet the blood cells for the collection of plasma. Plasma from the heparinized blood samples were used in part for the immunoglobulin and plasma biomarker measurements. Another fraction was used for clinical chemistry-based measures in the fasted and fed state. Hematological analyses were performed using one portion of EDTA blood samples derived from animals in the fed state. Another portion of EDTA blood samples was used for FACS-based quantification of leukocyte populations.

### Clinical chemistry

Assessments of clinical chemistry parameters were performed using an AU480 Automated Chemistry Analyzer (Beckman Coulter, Brea, CA, US) and specific kits for free fatty acids (Wako Chemicals, Neuss, Germany), glycerol (Randox Laboratories, Crumlin, UK) as well as all other parameters covered by the present study (Beckman Coulter, Brea, CA, US) ^90^. A set of 21 parameters consisting of specific metabolite levels, electrolyte concentrations and enzyme activities was measured in fed mice: albumin, α-amylase, alkaline phosphatase (AP), aspartate-aminotransferase (ASAT/GOT), alanine-aminotransferase (ALAT/GPT), calcium, cholesterol, chloride, creatinine, fructosamine, glucose, iron, lactate, lactate dehydrogenase (LDH), phosphate, potassium, sodium, total protein, triglycerides, urea and unsaturated iron-binding capacity. In addition, a selection of parameters (cholesterol, glucose, glycerol, HDL-cholesterol, non-esterified fatty acids (NEFA), non-HDL cholesterol and triglycerides) was measured in fasted animals after 4 (Ghrhr study), 6 (IF study) or 16-18 hours (mTOR study, C57BL/6J baseline study) of food deprivation. Plasma insulin levels were measured using a commercial kit based on immuno-electrofluorescence (Mesoscale Discovery, Rockville, Maryland, USA) or ELISA technology (Mercodia, Uppsala, Sweden).

### Hematology

EDTA-blood samples were diluted 1:5 in Sysmex Cell-Pack buffer (Sysmex, Kobe, Japan) prior to performing blood cell counts via a Sysmex XT2000iV device (Sysmex, Kobe, Japan). Parameters analyzed included red blood cell count (RBC), hematocrit (HCT), hemoglobin concentration (HBG), mean corpuscular hemoglobin content (MCH), mean corpuscular hemoglobin concentration (MCHC), mean corpuscular volume (MCV), red blood cell width distribution (RDW), platelet count (PLT), mean platelet volume (MPV), platelet distribution width (PDW), platelet large cell ratio (PLCR, >12 fl) and total white blood cell count (WBC).

### FACS-based analysis of peripheral blood leukocytes

Whole blood samples were incubated with Fc block (clone 2.4G2) for 5 min at 4–10 °C. Subsequently, blood leukocytes were stained using a mixture of fluorescence-conjugated monoclonal antibodies (BD Biosciences, Franklin Lakes, NJ, US) for 1 h at 4–10 °C. After lysis of erythrocytes and a formalin-based fixation, samples were analyzed using a Gallios ten-color flow cytometer (Beckman Coulter, Brea, CA, US) combined with an IntelliCyt HyperCyt sampler (Sartorius, Göttingen, Germany). The acquisition threshold (trigger) was set on the CD45-channel ^91^. A total number of 10,000-50,000 leukocytes per sample was examined. Frequencies of leukocyte populations were determined by software-based analysis (Flowjo, TreeStar Inc, USA; and SPICE ^92^). Gates for each parameter were based on formerly performed ‘fluorescence minus one’ (FMO) controls ^93^. Surface antigens used to define leukocyte populations were: B220, CD3, CD4, CD5, CD8, CD11b, CD11c, CD19, CD25, CD44, CD62L, Ly6C, Ly6G, NK1.1, NKp46 and gamma delta T cell receptor (gdTCR). Detailed information regarding the definition of relevant cell subpopulations is provided in **Extended Data Table 1** as well as **Extended Data Fig. 17 and 18**. The list of antibodies applied is provided in **Extended Data Table 2**.

### Analysis of cytokine and immunoglobulin abundance in plasma samples

The abundance of several plasma biomarkers was determined using MULTI-ARRAY technology MSD (Meso Scale Discovery, Rockville, MD, US). MSD-Mouse Isotyping Panel (IgA, IgM, IgG1, IgG2a, IgG2b, IgG3), MSD-Mouse Proinflammatory Panel (IFN-γ, IL-1β, IL-2, IL-4, IL-5, IL-6, IL-10, IL-12p70, KC/GRO) and MSD-U-PLEX Custom (IgE, Insulin, IL-6, and TNF-a) were quantified side by side on MULTI-SPOT plates.

### Lymphocyte proliferation assay

For monitoring T and B cell proliferation rates, single cell suspensions were prepared from spleens of mice at different ages. Cells were then stimulated *in vitro* as described below. Assessment of Class Switch Recombination was performed using stimulation with a mixture of anti-CD40 antibody and IL-4 (applied at concentrations of either 1 μg/ml anti-CD40 + 5 ng/ml IL-4 or 5 μg/ml anti-CD40 + 10 ng/ml IL-4) followed by a cultivation period of 7 days. For the assessment of T cell proliferation, cells were treated with a mixture of anti-CD3 antibody and IL-2 (at 1 µg/ml anti-CD3 + 1 ng/ml IL2 or 5 µg/ml anti-CD3 + 10 ng/ml IL-2) followed by cultivation for 3 days. The CellTiter-Glo (Promega) Luminescent Cell Viability Assay kit was used for proliferation measurements following the manufacturer’s instructions. The luminescence signal was read using a Microplate Reader (TECAN Infinite M200).

### Pathology

During necropsy, mice were examined morphologically. Body and organ weights/length measurements as well as any tissue lesions were documented by experienced necropsy technical personnel using an annotation approach developed for high-throughput mouse phenotyping (https://www.mousephenotype.org/impress/procedures/14). The following organs were collected, fixed in 4 % neutral buffered formalin and embedded in paraffin: abdominal aorta, adipose tissue (brown and white), adrenal gland, brain, bone (femur), epididymis, heart, intestine, kidney, liver, lung, pancreas, reproductive organs, skeletal muscle, skin, spleen, stomach, thymus, thyroid gland and urinary bladder. For histological examination, we generated 2-µm sections from the respective organ samples and stained them with either Haematoxylin-Eosin (HE), Periodic Acid Schiff (PAS), Van Gieson or Movat Pentachrome. Digital scans of stained slides were processed using a NanoZoomer HT2.0 slide scanning system (Hamamatsu Photonics, Hamamatsu, Japan) and were analyzed by two experienced pathologists.

### Organ harvest and processing for molecular analyses

After animals were sacrificed with CO_2_, organs (brain, lung and spleen) were harvested quickly, snap-frozen in liquid nitrogen and stored at −80 °C. Frozen tissue samples were then pulverized in liquid nitrogen using a porcelain mortar and pestle (MTC Haldenwanger, Waldkraiburg, Germany) maintained on dry ice. Several aliquots of tissue powder were made and stored at −80 °C until further use.

### RNA extraction, RNA-seq and data processing

Total RNA isolation from mouse tissues was performed with TRI-Reagent (Merck, Darmstadt, Germany). Briefly, 1 ml TRI-Reagent was added to an aliquot of frozen tissue powder followed by solubilization via ten passages through a 24-gauge needle. Further processing steps were performed according to the manufacturer’s recommendations. Total RNA was purified with the Monarch RNA Cleanup kit (New England Biolabs, Ipswich, MA, US).

A previously described protocol ^94^ was used for mRNA isolation and cDNA library preparation with a few modifications. Briefly, mRNA was isolated from purified 1 µg total RNA using oligo-dT beads (New England Biolabs, Ipswich, MA, US) and fragmented in reverse transcription buffer by incubating at 85 °C for 7 min, before cooling on ice. SmartScribe reverse transcriptase (Taraka Bio, Kusatsu, Japan) with a random hexamer oligo (HZG883: CCTTGGCACCCGAGAATTCCANNNNNN) was used for cDNA synthesis. Samples were then treated with RNase A and RNase H to remove RNA, followed by purification of cDNA on Agencourt AMPure XP beads (Beckman Coulter, Brea, CA, US). The single stranded cDNA was ligated with a partial Illumina 5’ adaptor (HZG885:/5phos/AGATCGGAAGAGCGTCGTGTAGGGAAAGAGTGTddC) using T4 RNA ligase 1 (New England Biolabs, Ipswich, MA, US) and incubated overnight at 22 °C. Ligated cDNA was purified on AMPure XP beads and amplified by 20 cycles of PCR using FailSafe PCR enzyme (Epicenter Technologies, Thane, India) and oligos that contain full Illumina adaptors (LC056: AATGATACGGCGACCACCGAGATCTACACTCTTTCCCTACACGACGCTCTTCCGATCT and unique index primers: CAAGCAGAAGACGGCATACGAGATnnnnnnnnnnGTGACTGGAGTTCCTTGGCACCCGAG AATTCCA, where nnnnnnnnnn indicates index nucleotides) for each sample. The resulting cDNA libraries were purified on AMPure XP beads, size selected using SPRIselect beads (Beckman Coulter, Brea, CA, US), and quantified by Qubit dsDNA HS Assay Kit (Thermo Fisher Scientific, Waltham, MA, US) prior to pooling. The pooled library was run on an Agilent High Sensitivity DNA chip (Agilent Technologies, Santa Clara, CA, US) with an Agilent 2100 Bioanalyzer instrument (Agilent Technologies, Santa Clara, CA, US) to check the quality and average fragment size.

Pooled indexed cDNA libraries were sequenced on an Illumina NovaSeq 6000 system (Illumina, San Diego, CA, US) with a single 111 bp read and 10 bp index read. Demultiplexing and data transformation to generate fastq files was done using bcl2fastq2 (v2.20). Sequencing reads were trimmed using CutAdapt (https://usegalaxy.org/) to remove adapter sequences. Trimmed reads were mapped to the mouse transcriptome (GRCm38, mm10) using HISAT2 (v2.1.0) in Galaxy (https://usegalaxy.org/) with forward strand information and default settings. Bam files were indexed using Samtools and count matrices generated by Genomic Alignments in R. Gene count matrices were generated using annotation information from a Mus_musculus.GRCm38.102.chr.gtf file imported with the rtracklayer ^95^ package into R. All downstream analyses were performed using R (Version 3.5.1, https://cran.r-project.org/). Library normalization and expression differences between samples were quantified using the DESeq2 package ^96^. A false discovery rate (FDR) < 0.05 was used as a cutoff in differential expression analyses. Assessment of the significance of the overlap of mTOR-and age-sensitive genes was performed based on exact hypergeometric probability.

Sample sizes for RNA-seq analyses were as follows: 3 months, n=7 mice; 5 months, n=9 mice; 8 months, n=8 mice; 14 months, n=9 mice; 20 months, n=7 mice; 26 months, n=5 mice (**Supplementary Data 2**, **3**); young/old mTOR mutant mice and WT littermate controls, each n=3 mice per group (**Supplementary Data 8**).

### Real-time quantitative PCR

Total RNA was reverse-transcribed by the iScript cDNA Synthesis Kit (Bio-Rad Laboratories, Hercules, CA, US). Real-time quantitative PCR based on the SYBR Green method was performed using the PowerUP SYBR Green Master Mix (Thermo Fisher Scientific, Waltham, MA, US) on a StepOnePlus Real-Time PCR System (Thermo Fisher Scientific, Waltham, MA, US). The threshold cycle value (Ct) of each target gene was normalized to the corresponding Ct value of β-actin (*Actb*). Primer sequences used are provided in **Extended Data Table 4**.

### Protein isolation

To generate tissue homogenates suitable for measuring levels of lipid peroxidation, 20S proteasome activity as well as reactive oxygen species production, 300 µl Tris-Buffered Saline (TBS, pH 7.6) and 1% Triton X-100 (Merck, Darmstadt, Germany) were added to a tissue powder aliquot and homogenization was performed using ten consecutive passages through a 24-gauge needle on ice. After a 30 min incubation step on ice, samples were centrifuged at 15000 x g for 30 min at 4 °C. The supernatant was aliquoted into new tubes and stored at −80 °C until use. To generate protein homogenates suitable for western blot analysis, 300 µl Tris-Buffered Saline (TBS, pH 7.6) + 1% Triton X-100 (Merck, Darmstadt, Germany) supplemented with 1x Protease Inhibitor Cocktail (Roche, Basel, Switzerland) and 1x PhosSTOP Phosphatase Inhibitor Cocktail (Roche, Basel, Switzerland) was added to a tissue powder aliquot. Further processing steps were analogous to the ones described above.

### Lipid peroxidation

Levels of thiobarbituric acid reactive substances (TBARS), a byproduct of lipid peroxidation, were determined using a TBARS Assay Kit (Cayman Chemical, Ann Arbor, MI, US) following the manufacturer’s instructions.

### Proteasome activity

The 20S proteasome activity was assessed *in vitro* using a protocol published elsewhere ^97^. Developing buffer (50 mM Tris (pH7.5), 150 mM NaCl, 5 mM MgCl_2_) containing 30 µg protein in a total volume of 94 µl was loaded onto a black 96-well plate (Sarstedt, Nümbrecht, Germany). Two µl 50 mM ATP-Mg^2+^ (Merck, Darmstadt, Germany), 2 µl 5 mM Suc-LLVY-AMC substrate (Cayman Chemical, Ann Arbor, MI, US) and 2 µl 1% SDS (Carl Roth, Karlsruhe, Germany) were added immediately prior to commencing the experiment. Reaction suspensions were incubated in the dark at 37 °C for 30 min and fluorescent signals (excitation 380 nm, emission 460 nm) were acquired using a Tecan Infinite M200 Pro plate reader (Tecan, Männedorf, Switzerland).

### Reactive oxygen species

Production of reactive oxygen species (ROS) in mouse tissues was measured *in vitro* using a previously described protocol ^98^ with a few modifications. All chemicals used in this assay were purchased from Merck (Darmstadt, Germany). We mixed 45 µl protein samples (30 µg protein) with 50 µl ice-cold 2x Locke’s buffer (308 mM NaCl, 11.2 mM KCl, 7.2 mM NaHCO_3_, 4 mM CaCl_2_, 20 mM D-glucose, 10 mM HEPES (pH7.4)) and loaded this mixture onto a black 96-well plate (Sarstedt, Nümbrecht, Germany). We then added 5 µl 200 µM 2’,7’-dichlorodihydrofluorescein diacetate (DCFH-DA) and the reaction mixture was incubated in the dark on an orbital shaker for 30 min at room temperature and 50 rpm. Fluorescent intensities (excitation 485 nm, emission 530 nm) were acquired using a Tecan Infinite M200 Pro plate reader (Tecan, Männedorf, Switzerland).

### Western blot

All chemicals and reagents used in this procedure were purchased from Merck (Darmstadt, Germany), if not specified otherwise. We mixed 30 mg protein samples with the appropriate amount of 4x loading buffer (240 mM Tris-HCl (pH 6.8), 8% SDS, 5% beta-mercaptoethanol, 40% glycerol and 0.04% bromophenolblue) and ran this mixture through handcast Tris-glycine gels prior to blotting onto nitrocellulose membranes with 0.1 µm pore size (GE Healthcare, Chicago, IL, US). Subsequently, the membranes were incubated with Phosphate-Buffered Saline (PBS) + 10% skim milk powder for 1h at room temperature to reduce background noise. Primary antibody solutions, diluted in PBS + 1% milk, were applied overnight at 4 °C. A detailed list of primary antibodies used is provided in **Extended Data Table 5**. Secondary antibodies, either goat anti-rabbit (Promega, Madison, WI, US) at 1:3000 dilution or goat anti-mouse (Agilent Technologies, Santa Clara, CA, US) at 1:10000 dilution, were used at room temperature with an incubation time of 90 min. Immunoreactivity was visualized via enhanced chemiluminescence (Advansta, Menlo Park, CA, US) and band densities were quantified using ImageJ software (version 1.50e, National Institute of Health). Phosphorylated proteins were normalized to the respective total protein band on the same lane. In all other cases, normalization was carried out using the Actin signal derived from the same lane. Whenever multiple protein targets were determined on the same membrane, signals derived from preceding visualizations were erased by adding a sodium azide incubation step in between protein target measurements.

### Aging-related (endo)phenotypes in humans

The human data were collected in the context of the Rhineland Study, which is an ongoing, large-scale, single-center, population-based prospective cohort study among people aged 30 years and above in Bonn, Germany. The only exclusion criterion is insufficient command of the German language to provide informed consent. Persons living in the recruitment areas are predominantly German from Caucasian descent. One of the Rhineland Study’s primary objectives is to identify determinants and markers of healthy aging, utilizing a deep-phenotyping approach. Approval to undertake the study was obtained from the ethics committee of the University of Bonn, Medical Faculty. We obtained written informed consent from all participants in accordance with the Declaration of Helsinki.

For these analyses, we used baseline data of the first 3034 participants of the Rhineland Study who had both phenotype and genotype data available. Fifty-four aging-related (endo)phenotypes representing ten physiological functional domains, including body composition (n=5), body fat distribution based on magnetic resonance imaging (MRI) (n=4), cardiology (n=7), clinical chemistry (n=11), hematology (n=9), inflammation (n=4), immunology (n=6), muscle strength (n=2), ophthalmology (n=2) and physical activity (n=4), were included. Further details of the study have been described previously ^30, 99, 100^.

### Statistics and data analysis

Phenotypic and molecular data were analyzed across age groups using one-way ANOVAs with the between-subjects factor age, followed by Fisher’s LSD posthoc analyses if appropriate (using base R version 3.6.1 and the package ‘agricolae’ version 1.3-1). We analyzed non-parametric data across age-groups using Kruskal-Wallis tests, followed by Dunn tests where appropriate (using base R version 3.6.1 and the package ‘FSA’ version 0.8.26). Count-based data (histopathology) were analyzed using Fisher’s exact test across all age groups, followed by pairwise comparisons against the 3 months old reference group if appropriate (using base R version 3.6.1 and the package ‘rcompanion’ version 2.3.7). Throughout the manuscript, we report two-tailed p-values. Age-sensitivity of parameters was determined by evaluating the p-value (p<0.05) of the global comparison (i.e., ANOVA, Kruskal-Wallis or Fisher’s exact test). The age at first detected phenotypic change was determined by assessing the results of the posthoc tests vs. the 3 months old reference group. For instance, we considered a phenotype to be age-sensitive and to feature an age at first detected change of 8 months if the global test (i.e., ANOVA, Kruskal-Wallis or Fisher’s exact test) for an age effect was significant (p<0.05) and the posthoc analyses vs. the 3 months old group were not significant for the comparison 3 vs. 5 months, but significant for all other tests (3 vs. 8, 3 vs. 14, 3 vs. 20, 3 vs. 26) with the maximum possible exception of one comparison. For PCA of phenotypic data, all continuous variables available were included in the analysis. PCA was performed using base R version 3.6.1 and the packages ‘FactoMineR’ (version 2.0) and ‘factoextra’ (version 1.0.6). Multivariate imputation by chained equations was used for imputation in case of missing values (using base R version 3.6.1 and the package ‘mice’ version 3.13.0).

Phenotypic data from our intervention studies (*mTOR*, *Ghrhr*, IF) were analyzed using two-way ANOVAs with the between-subjects factors age and intervention (genotype or diet). We used a full ANOVA model, including an intervention × age interaction term. These analyses were followed by Fisher’s LSD posthoc tests where appropriate (using base R version 3.6.1 and the package ‘agricolae’ version 1.3-1). Non-parametric data (SHIRPA outcomes, data from auditory brain stem responses) were analyzed using Aligned Rank Transform with the factors age and intervention (including an intervention × age interaction term), followed by Mann-Whitney U tests for posthoc analyses if appropriate (using base R version 3.6.1 and the package ‘ARTool’ version 0.10.6). Count-based data (histopathology) were analyzed using Fisher’s exact test across age and intervention groups, followed by pairwise comparisons if appropriate (using base R version 3.6.1 and the package ‘rcompanion’ version 2.3.7). We visualized the relationship of age and intervention effects using Venn diagrams (shown in panels c of Figures 3-5). These show the numbers of age-sensitive phenotypes (parameters with a main effect of age, p<0.05), genotype/diet-sensitive phenotypes (parameters with a main effect of genotype/diet), the number of parameters with a significant genotype/diet × age interaction (p<0.05), as well as the intersection between these parameter sets. Sunburst diagrams in panels c of Figures 3-5 focus on all age-sensitive phenotypes (ASPs) shown in the Venn diagram and dissect these into ASPs not influenced by genotype/diet, ASP accentuated by genotype/diet and ASP ameliorated by genotype/diet. To address whether age-sensitive phenotypes in the intersection (associated with both age and diet) are ameliorated or accentuated by the intervention (genotype/diet) we evaluated effect sizes (Cohen’s d; computed using the R package effsize version 0.7.6) of age, as well as intervention in the old group: If Cohen’s d values pointed in the same direction, ASPs were considered to be accentuated, if they were in opposing directions, ASPs were considered to be ameliorated by the intervention. ASPs were not considered in cases where Cohen’s d values could not be computed (because of 0 values in the denominator). For ASPs ameliorated by genotype/diet, the sunburst diagrams also provide the proportion of ASPs influenced via either a genotype/diet main effect and/or via a genotype/diet × age interaction (inner ring of the sunburst diagrams). The outer ring of the sunburst diagrams shows the proportional distribution of the age at first detected phenotypic change of ASPs (based on the parameters for which this information had been collected in the context of our aging trajectories baseline study). We used an analogous approach for the analysis of gene expression data: In this case, the Venn diagram shows the number of age-sensitive genes (genes with a main effect of age, FDR<0.05), genotype-sensitive genes (genes with a main effect of genotype, FDR<0.05), genes with a significant genotype × age interaction (FDR<0.05), as well as the intersection between these parameter sets. Instead of Cohen’s d values, we used log2 fold changes for the comparisons of age and genotype effects but otherwise the proceeding was analogous to the approach used for the assessment of phenotypic data described above. Linear regression analyses of effect size estimates were carried out using GraphPad Prism version 8 (La Jolla, CA, US). We performed intraclass correlation analyses using base R version 3.6.1 and the package ‘psych’ (version 2.0.7). Statistical comparison of Cohen’s d effect sizes for individual phenotypes was carried out as described in ^78^ (for further details, see also our analysis code available at https://github.com/ehningerd/Xie_et_al-longevity_regulators).

For the statistical analyses in humans, characteristics of the study participants were reported as means (standard deviations (SD) and ranges) for continuous variables and numbers and percentages for categorical variables. The eQTL dosage was coded as GG=0, AG=1, and AA=2 for *GHRHR*, and as GG=0, CG=1, and CC=2 for *MTOR*. All variables representing (endo)phenotypes were standardized before further analyses in order to enable better comparison of the effect sizes across different physiological and functional domains. Given the low rate of missingness (<5%), all analyses were based on cases with complete data. Multiple linear regression analyses were applied to quantify the association between eQTL dosage (determinant) and each (endo)phenotypic measure (outcome). Models were adjusted for age, sex and population stratification using the first ten genetic principal components. In addition, inflammation markers were adjusted for batch information, MRI-based fat measurements were adjusted for height of the segmented region as described previously ^30, 99, 100^, and physical activity variables were adjusted for examination season and the number of valid recoding days. For (endo)phenotypes where a *GHRHR*- and/or an *MTOR*-genotype effect was observed, we further assessed the interaction between age and each eQTL to evaluate age-(in)dependency of the effect estimates. All standardized effect estimates are reported together with their 95% confidence intervals (CIs).

Statistical analyses were carried out using R version 3.6.1, including the packages FactoMineR_2.0, factoextra_1.0.6, ggplot2_3.2.1, psych_2.0.7, ARTool_0.10.6, mice_3.13.0, effsize_0.7.6, car_3.0-5, Rmisc_1.5, agricolae_1.3-1, rcompanion_2.3.7, FSA_0.8.26, lattice_0.20-38, plyr_1.8.5, and GraphPad Prism version 8 (La Jolla, CA, US). Pathway enrichment analyses were performed using Ingenuity Pathway Analysis version 01-18-06 (Ingenuity Systems, Redwood City, CA, US).

### Data and code availability

Raw phenotypic and molecular data from Fig. 2, 3, 4 and 5 were deposited on Mendeley at https://data.mendeley.com/datasets/ypz9zyc9rp/draft?a=09b16f74-4581-48f7-94af-469e01757949. Raw sequencing data from Fig. 2, **Extended Data Fig. 3 and 9** are available through GEO datasets at accession number GSE168068. Analysis code is available at https://github.com/ehningerd/Xie_et_al-longevity_regulators.

## Supplementary results

We sought to extract from our dataset ASPs sensitive to PAAI-mediated amelioration specifically in the old group (but not the young group) by selecting phenotypes with an overall significant main effect of age (on the 2-way ANOVA) and a significant difference on the posthoc test between the old intervention group and the old control group, but not on the comparison young intervention group vs. young control group (see **Supplementary Data 6,7,9** for full information on results from statistical analyses which these analyses are based upon). This would be ASPs corresponding to the “rate effect model” introduced in Fig. 1b.

The analysis of our *Ghrhr^lit/lit^* dataset revealed that 7.3% of all ASPs (corresponding to 7 ASPs) followed this pattern (i.e., showed a significant difference between mutant and control in old but not young mice) (**Extended Data Fig. 7a**). Statistical comparison of *Ghrhr^lit/lit^* effect sizes in young vs. old mice also identified one of these ASPs as significantly different between age groups (activity of Alkaline Phosphatase in the blood plasma; **Extended Data Fig. 7b**).

In the case of our *mTOR^KI/KI^* cohort, 15.4% of all ASPs (corresponding to 18 ASPs) showed a significant effect of genotype in the old but not the young group based on the posthoc tests (**Extended Data Fig. 7a**). The effect size plot in **Extended Data Fig. 7c** examines how this subset of ASPs was influenced by genotype in the old vs. the young group. This analysis confirms that, based on statistical comparison of Cohen’s d effect sizes, several ASPs were differentially ameliorated by *mTOR^KI/KI^* genotype in the old vs. the young group (p<0.05; hemoglobin, hematocrit, plasma triglyceride concentration, subpopulations of CD4+ T cells). However, many of these ASPs appeared to show similar effect sizes in the young vs. the old group of animals (**Extended Data Fig. 7c**). Intraclass correlation analyses of effect sizes in young vs. old mice for this set of ASPs revealed an overall significant correlation (ICC=0.49, p=0.01; **Extended Data Fig. 7c**), suggesting that our strategy to extract ASPs of interest (i.e., ASPs selectively ameliorated in old mice) based on the pattern of posthoc results may generate some false positives.

The analysis of our IF cohort revealed that 22.5% of all ASPs (corresponding to 23 ASPs) followed this pattern (**Extended Data Fig. 7a**). Several of these ASPs were also corroborated by comparison of effect sizes in young vs. old mice, such as plasma insulin concentration, plasma urea concentration, respiratory exchange ratio and the abundance of NKT cells (**Extended Data Fig. 7d**). However, we again noted that in a number of cases diet effect sizes appeared to be similar in young and old mice (despite the posthoc test not revealing a difference between the young IF and young control group upon selection of these ASPs) with an overall significant intraclass correlation of diet effect sizes in young vs. old mice in this set of ASPs (ICC=0.44; p= 0.01; **Extended Data Fig. 7d**).

In conclusion, while these analyses were able to identify ASPs whose selective amelioration in the old group of mice is convincing (see examples discussed above; see also yellow datapoints in effect size plots shown in **Extended Data Fig. 7b-d**), it also suggested some ASPs that are likely false positives (given that effect sizes in the young group were similar to those in the old group). Based on these analyses, the upper bound of our estimate of ASPs following the pattern of selective amelioration in the old group is the one shown in **Extended Data Fig. 7a**. A lower bound may be derived from the number of ASPs with a significant effect size difference between young and old mice (i.e., the yellow datapoints in **Extended Data Fig. 7b-d**); this would suggest that about 1% of all ASPs in the *Ghrhr^lit/lit^* dataset, 4.3% of all ASPs in the *mTOR^KI/KI^* cohort and 5.9% of all ASPs in the IF dataset correspond to ASPs selectively ameliorated in old mice but not young mice (i.e., ASPs corresponding to the “rate effect model” introduced in Fig. 1b).

## Supplementary discussion

Our analyses generated a large dataset on phenotypes associated with *Ghrhr* loss of function in mice. Novel findings in *Ghrhr^lit/lit^* mice included, for instance, a higher auditory sensitivity, reduced visual acuity as well as an electrocardiographic shortening of the PR interval that may predispose for arrhythmias. In other cases, we confirmed previously reported effects of growth hormone deficiency, such as reduced bone mineral density ^101^, higher nociceptive sensitivity ^102^, as well as changes in body composition and metabolism ^103^, which we found across age groups in *Ghrhr^lit/lit^* mice. Our observation of reduced activity levels in young and old *Ghrhr^lit/lit^* mice, notable across different assays employed (open field, SHIRPA and metabolic phenotyping) is in contrast to a prior report of increased locomotor activity in *Ghrh* (encoding growth hormone releasing hormone) mutant mice ^104^.

Previous work had established that hypomorphic mTOR mutant mice feature a ca. 20% extension of median lifespan which was associated with a reduced incidence of neoplastic diseases in the mutants ^26^. Lifespan studies using the oral mTOR inhibitor rapamycin in mice had yielded median lifespan extensions ranging from 4-26%, depending on dose, age at onset of treatment, sex and site of investigation ^9, 40, 105, 106^. A large number of the phenotypic effects we observed in mTOR mutants were similar to effects seen under chronic treatment with the pharmacological mTOR inhibitor rapamycin ^16, 39^: For instance, both the genetic and pharmacological manipulations were associated with age-independent increases in exploratory locomotor activity, red blood cell counts, naïve CD4^+^-and CD8^+^-T-cell counts as well as age-independent decreases in hepatic microgranulomas, bronchus-associated lymphatic tissue and unsaturated iron binding capacity. Moreover, both were also associated with a prevention of age-related cardiac hypertrophy and a reduced cancer incidence in old mice and shared adverse effects, such as testicular degeneration, impaired glucose tolerance and an exacerbation of the age-related decrease in NK cells.

However, we also noted a number of effects seen in the mTOR mutants, which we did not observe in mice under chronic rapamycin treatment ^16^. For instance, while the specific rapamycin treatment approach we employed previously ^16^ did not have consistent effects on body and organ weights across treatment cohorts (heart, liver, spleen, brain, kidney; an exception was testis with dramatically reduced weights due to testicular degeneration), the mTOR mutant allele led to clear reductions in body mass, organ weights (brain, heart, kidney, liver, lung, muscle, pancreas, spleen and testis) and reduced retinal thickness. Additional phenotypic effects restricted to the mTOR mutants included a protection against age-related glomerular pathology and elevations in white blood cell and platelet counts. While some of these differential effects may be a matter of rapamycin dosage (e.g., body weight reductions were also seen with higher rapamycin doses ^105^), others may not (e.g., chronic oral rapamycin was associated with renal toxicity ^16^; mTOR mutants, in contrast, were protected against age-related glomerular pathology and showed no signs of renal toxicity). One limitation of the hypomorphic mTOR mutant mouse model is that it is associated with some degree of embryonic lethality ^25–27^. Advantages, relative to (oral) pharmacological approaches, include the specific targeting of mTOR (due to the genetic nature of the manipulation) as well as the fact that mTOR inhibition is independent of food intake (which typically declines in old mice).

## Extended Data Figure legends

**Extended Data Figure 1: Schematic illustration of the analytical workflow of the current study.** The figure summarizes our analytical approach. We performed large-scale phenotypic analyses in 3-month, 5-month, 8-month, 14-month, 20-month and 26-month old C57BL/6J mice to identify age-sensitive phenotypes (ASPs) and estimate their aging trajectories. To identify ASPs, we performed one-way ANOVA with the between-subjects factor age (or Kruskal-Wallis test in the case of non-parametric data). For each ASP, we used posthoc analyses to determine at which age phenotypes first differed significantly from the 3-month reference group (results are presented in Fig. 2a–e and fully described in **Supplementary Data 1**). We carried out PCA to visualize how these six age groups differed from each other when extracting the first 2 principal components from this multidimensional dataset (results are presented in Fig. 2f).

We examined three pro-longevity interventions for their effects on age-dependent phenotypic change. For each intervention, we carried out large-scale phenotypic analyses using a study design that included a young control group, a young intervention group, an aged control group and an aged intervention group.

To visualize overall age and intervention effects in our multidimensional dataset, we carried out PCA on all continuously distributed phenotypes. We provide, for each animal, the values of the first 2 principal components in a scatter plot (results are presented in Fig. 3–5b; compare to schematics outlined in Fig. 1b).

On the level of individual phenotypes, we used two-way ANOVAs with the between-subject factors age and intervention (or aligned rank transform in the case of non-parametric data) to extract main effects of age, main effects of intervention as well as intervention × age interactions (**Extended Data Fig. 5**; **Supplementary Data 6**, **7** and **9**). These analyses help to differentiate, on the level of individual phenotypes, between the “rate effect” model as well as “combination of rate effect and baseline effect” model on the one hand (Fig. 1b, left and middle panels; ASPs with a significant interaction term) and the “baseline effect model” on the other hand (Fig. 1b, panels to the right; ASPs without a significant interaction term). We show Venn diagrams featuring the number of phenotypes with main effects and/or an interaction (Fig. 3–5c). We further examine phenotypes with a main effect of age (age-sensitive phenotypes, ASPs): Sunburst charts show the proportion of ASPs opposed (effect sizes of age and intervention are in opposing directions; in green), accentuated (effect sizes of age and intervention are in the same direction; in magenta) or not influenced by an intervention (in grey) (Fig. 3–5c). For ASPs ameliorated by an intervention, the inner circle of the sunburst chart shows the proportion of ASPs that features a significant main effect of intervention and/or a significant intervention × age interaction. The outer circle of the sunburst chart shows at which age changes in the corresponding ASPs were first detected based on data available from our baseline study. We carried out posthoc analyses in an attempt to identify ASPs opposed by intervention only in the old but not in the young group of mice (**Extended Data Fig. 7**); these analyses were meant to identify ASPs consistent with the “rate effect” model (Fig. 1b, left panels).

To show how intervention effect sizes in young mice relate to intervention effect sizes in aged mice (overall and on the level of individual phenotypes), we provide effect size plots for different subsets of phenotypes: 1) ASPs countered by intervention (i.e., ASPs with a significant main effect of intervention and/or a significant intervention × age interaction and effect sizes of age and intervention that go in opposing directions; this corresponds to the central green section of the sunburst chart) (Fig. 3–5g). 2) ASPs accentuated by intervention (i.e., ASPs with a significant main effect of intervention and/or a significant intervention × age interaction and effect sizes of age and intervention that go in the same direction; this corresponds to the central magenta section of the sunburst chart) (Fig. 3–5h). 3) Phenotypes featuring a main effect of intervention and/or an intervention × age interaction but not a main effect of age (Fig. 3–5i).

We examined, for each phenotype individually, whether intervention effects differed significantly between young and old mice (phenotypes with significant differences are highlighted in the effect size plots) (Fig. 3–5f–i). These analyses were also used to help differentiate, on the level of individual phenotypes, between the “rate effect” model as well as “combination of rate effect and baseline effect” model on the one hand (Fig. 1b, left and middle panels; ASPs with a significant difference in intervention effect size when comparing young vs. old mice) and the “baseline effect model” on the other hand (Fig. 1b, ASPs without a significant difference in intervention effect size when comparing young vs. old mice). We performed linear regression to test how well effects in young and old mice are correlated across these sets of phenotypes (Fig. 3–5f–i). Additionally, we performed intraclass correlation analyses which reflect not only the degree of correlation but also the agreement between measures in the young and old group (Fig. 3–5f–i). These analyses were performed to help differentiate between the models outlined in Fig. 1b.

**Extended Data Figure 2: Pathological findings in aging C57BL/6J mice.** The graphs show the relative proportion of animals in each age group affected by inflammation in the accessory glands (**a**, scale bar: 250 µm), inflammatory infiltrates in the epididymides (**b**, scale bar: 500 µm), heart fibrosis (**c**, scale bar: 1 mm), chronic progressive nephropathy (**d**, scale bar: 250 µm), perivascular infiltrates in the kidneys (**e**, scale bar: 500 µm), tubular regeneration in the kidneys (**f**, scale bar: 250 µm), lateral meniscus tissue structure changes in the knees (**g**, scale bar: 1 mm), Russel bodies in the spleen (**h**, scale bar: 250 µm), adenoma (**i**, scale bar: 250 µm) or goiter (**j**, scale bar: 250 µm) of the thyroid gland. Representative examples of histopathological findings in older mice (alongside healthy tissue in younger mice) are shown in the images accompanying the graphs. Data are based on n=5 mice per age group. For further details, see **Supplementary Data 1**.

**Extended Data Figure 3: RNA-seq-based transcriptome analysis captures gene expression changes in the brain across the lifespan in male C57BL/6J mice.** Ingenuity Pathway Analysis shows top canonical pathways, diseases and biological functions as well as predicted upstream regulators of genes differentially expressed in the brain relative to the 3-month old group (FDR<0.05). Positive z-scores (in orange) indicate activating effects, while negative z-scores (in blue) indicate inhibitory effects on corresponding processes.

**Extended Data Figure 4: Western-blot-based quantification of proteins linked to hallmarks of aging.** Representative band densities are shown for proteins detected in brain (**a**), lung (**b**) and spleen (**c**).

**Extended Data Figure 5: PAAIs - systematic analysis of main effects of age, main effects of intervention and intervention × age interactions. a**,**d**,**g**: These plots show, for all 3 PAAIs examined in the present paper, cumulative frequencies of −log10(p-values) for age effects, intervention effects and intervention × age interactions for all phenotypes analyzed via two-way ANOVA or aligned rank transform. The vertical dotted line marks the significance threshold (p<0.05; corresponding to ∼1.3 after the log transformation and multiplication with - 1). **b**,**e**,**h**: These scatter plots show, for all PAAIs assessed, −log10(p-values) for intervention main effects plotted vs. −log10(p-values) of intervention × age interactions for all phenotypes analyzed via two-way ANOVA or aligned rank transform. The vertical and horizontal dotted lines mark the significance threshold (p<0.05). The graphs also show regression lines, correlation coefficients and p-values derived from linear regression analyses. **c**,**f**,**i**: These scatter plots show, for all PAAIs assessed, −log10(p-values) for intervention main effects plotted vs. −log10(p-values) of intervention × age interactions for age-sensitive phenotypes countered by intervention. The vertical and horizontal dotted lines mark the significance threshold (p<0.05). The graphs also show regression lines, correlation coefficients and p-values derived from linear regression analyses.

**Extended Data Figure 6: Survival and pathological analyses in *Ghrhr^lit/lit^* and hypomorphic *mTOR^KI/KI^* mice.** The figure shows provisional survival data as well as summarizes histopathological analyses for aging *Ghrhr^lit/lit^* (**a**-**g**) and *mTOR^KI/KI^* (**h**-**p**) mice and the corresponding WT littermate controls. **a**,**h**: Provisional survival curves were established based on cases of natural deaths in *Ghrhr^lit/lit^* and *mTOR^KI/KI^* cohorts aged in our facility (p-values shown are based on analyses via Log-rank (Mantel-Cox) test). **b**-**d**, **i**-**j**: These panels show the percentage of aged mutant vs. WT animals affected by the pathological findings specified in the graphs (p-values are based on analysis via Fisher’s exact test). *p<0.05, **p<0.01, ***p<0.001, ****p<0.0001. BALT: bronchus-associated lymphoid tissue; KALT: kidney-associated lymphoid tissue. **e**-**g**, **k**-**p**: The images show representative examples of histopathological findings in aged mice as well as the corresponding healthy intact tissue in young animals. **e**, Lipofuscin deposits in the adrenal gland; scale bar: 100 µm. **f**, Bronchus-associated lymphoid tissue (BALT); scale bar: 500 µm. **g**, Thyroid gland adenoma; scale bar: 250 µm. **k**, Lymphoid infiltrates in the liver; scale bar: 500 µm. **l**, Microgranulomas in the liver; scale bar: 50 µm. **m**, Bronchus-associated lymphoid tissue (BALT); scale bar: 500 µm. **n**, Glomerular lesions in the kidney; scale bar: 50 µm. **o**, Kidney-associated lymphoid tissue (KALT); scale bar: 500 µm. **p**, Tubular degeneration in the testis; scale bar: 100 µm. Additional information is available in **Supplementary Data 6** and **7**.

**Extended Data Figure 7: Analysis of ASPs sensitive to PAAI-mediated effects specifically in the old groups of mice. a**, Percentage of ASPs that feature a significant intervention effect in the old group (posthoc test old invention group vs. old control group, p<0.05), but not the young group of animals (posthoc test young intervention group vs. young control group, p>0.05). **b**-**d**, Effect size plots show Cohen’s d effect sizes of intervention (**b**: *Ghrhr^lit/lit^* vs. WT; **c**: *mTOR^KI/KI^* vs. WT; **d**: IF vs. AL) in the young group vs. the old group of animals. To assess overall relationships between phenotypic intervention effect sizes in young vs. old animals, we performed linear regression (see correlation coefficient R, p-value, linear regression equation; black line: regression line; blue line: line through origin with slope 1) and intraclass correlation (see ICC, p-value) analyses. The graphs also show whether individual phenotypes had significantly different effect sizes in young vs. old mice (phenotypes with significant differences are identified by their abbreviated name; see **Supplementary Data 6, 7** and **9** for full description).

**Extended Data Figure 8: Effect sizes of *GHRHR* and *mTOR* eQTLs on gene expression levels.** Violin plots of *GHRHR* expression levels (**a**), stratified by the cis-eQTL at SNP rs11772180 (chr7: 30810998_A_G), as well as *MTOR* expression levels (**b**), stratified by the cis-eQTL at SNP rs2295079 (chr1: 11262508_C_G), as obtained from the Genotype Tissue Expression portal (genome build 38). Note that eQTLs for *GHRHR* have thus far only been assessed in human liver tissue. The *MTOR* eQTL has been validated in a wide range of human tissues, including brain, heart, skin, muscle and various gastrointestinal tissues (**b** shows expression levels in blood). The numbers below the horizontal axes indicate the number of samples assessed for each genotype for estimating gene expression levels. The shaded regions represent the density distributions of the samples for each genotype. The box plots indicate the interquartile ranges (black) and the median value (white lines) of gene expression for each genotype.

**Extended Data Figure 9: RNA-seq-based transcriptome analysis of spleen in young and old *mTOR^KI/KI^* mice as well as wildtype littermate controls.** This figure summarizes the results of an RNA-seq-based differential expression analysis comparing gene expression in the spleen of young and old *mTOR^KI/KI^* mice as well as WT littermate controls (n=3 per group). **a**, Venn diagram shows the number of age-sensitive genes (FDR<0.05), genotype-sensitive genes (FDR<0.05), genes with an interaction (FDR<0.05) as well as the intersection of these sets. **b**, Sunburst chart shows the number of age-sensitive genes either unaltered (in grey), countered (in green) or accentuated (in magenta) by the *mTOR^KI/KI^* genotype. For age-sensitive genes (ASGs) countered by *mTOR^KI/KI^* genotype, the inner ring shows the proportion of genes with a main effect of genotype (in dark green), a genotype × age interaction (in violet) or both a main effect and an interaction (in yellow). The outer ring shows when changes in the corresponding ASGs were first detected based on data available from our baseline study.

**Extended Data Figure 10: Cluster analysis of phenotypes in *Ghrhr^lit/lit^* cohort.** The figure shows results of hierarchical clustering applied to the phenotypic data obtained in the context of our *Ghrhr^lit/lit^* cohort. In order to be able to see what relationships might exist between phenotypes within our young control group, we performed hierarchical clustering on the young WT animals only (hence, yielding phenotype clusters that are independent of age- and genotype-associated phenotypic variation); the resulting clusters and distances between them can be extracted from the dendrogram shown in the figure. The heatmap to the left demonstrates standardized phenotype values for all phenotypes and animals (including young mutant mice and the old groups). How many clusters one identifies depends on the distance at which the dendrogram is cut. Analyses of genotype influences on clusters derived from different ways to cut the dendrogram (based on different minimal inter-cluster distances) are summarized in **Supplementary Data 10**.

**Extended Data Figure 11: Cluster analysis of phenotypes in *mTOR^KI/KI^* cohort.** The figure shows results of hierarchical clustering applied to the phenotypic data obtained from our *mTOR^KI/KI^* cohort. In order to be able to see what relationships exist between phenotypes within our young control group, we performed hierarchical clustering on the young WT animals only (hence, yielding phenotype clusters that are independent of age- and genotype-associated phenotypic variation); the resulting clusters and distances between them can be extracted from the dendrogram shown in the figure. The heatmap to the left demonstrates standardized phenotype values for all phenotypes and animals (including young mutant mice and the old groups). How many clusters one identifies depends on the distance at which the dendrogram is cut. Analyses of genotype influences on clusters derived from different ways to cut the dendrogram (based on different minimal inter-cluster distances) are summarized in **Supplementary Data 11**.

**Extended Data Figure 12: Cluster analysis of phenotypes in IF cohort.** The figure shows results of hierarchical clustering applied to the phenotypic data obtained from our IF cohort. In order to be able to see what relationships exist between phenotypes within our young control group, we performed hierarchical clustering on the young AL animals only (hence, yielding phenotype clusters that are independent of age- and diet-associated phenotypic variation); the resulting clusters and distances between them can be extracted from the dendrogram shown in the figure. The heatmap to the left demonstrates standardized phenotype values for all phenotypes and animals (including young IF mice and the old groups). How many clusters one identifies depends on the distance at which the dendrogram is cut. Analyses of diet influences on clusters derived from different ways to cut the dendrogram (based on different minimal inter-cluster distances) are summarized in **Supplementary Data 12**.

**Extended Data Figure 13: Deep phenotyping analyses of age-dependent changes in tumor-free C57BL/6J mice**. **a**–**d**, Deep phenotyping results in wildtype tumor-free C57BL/6J mice. **a**, Principal component analysis of deep phenotyping data (number of mice: 3-month old, n=15; 5-month old, n=14; 8-month old, n=15; 14-month old, n=13; 20-month old, n=15; 26-month old, n=13). **b**, Relative proportion of age-sensitive phenotypes among all phenotypes examined. **c**,**d**, Age at first detectable change (**c**) and age at full manifestation (**d**) of age-sensitive phenotypes (ASPs) shown as proportion of all ASPs.

**Extended Data Figure 14: Anti-aging effects induced by the *Ghrhr^lit/lit^* mutation; analysis restricted to tumor-free mice**. **a**, Principal component analysis of deep phenotyping data (number of mice: young WT, n=30; young *Ghrhr^lit/lit^*, n=20; old WT, n=25; old *Ghrhr^lit/lit^*, n=29). **b,** Venn diagram shows the number of age-sensitive phenotypes, genotype-sensitive phenotypes, phenotypes with a genotype × age interaction and their intersection. **c**, Sunburst chart shows the number of age-sensitive phenotypes either unaltered (in grey), counteracted (in green) or accentuated (in magenta) by the *Ghrhr^lit/lit^* mutation. For age-sensitive phenotypes counteracted by the *Ghrhr^lit/lit^* mutation, the inner ring shows the proportion of phenotypes with a main effect of genotype (in dark green), a genotype × age interaction (in violet) or both a main effect and an interaction (in yellow). The outer ring shows when changes in the corresponding ASPs were first detected based on data available from our baseline study. **d**– **g**, Scatter plots show the effect size of *Ghrhr^lit/lit^* genotype in young mice plotted vs. the effect size of *Ghrhr^lit/lit^* genotype in old mice for different sets of phenotypes: **e**, ASPs counteracted by genotype via a main effect and/or an interaction (i.e., corresponding to the central green section of the sunburst chart in **c**); green dots denote phenotypes in which genotype effects in young and old mice did not differ significantly; orange denotes phenotypes in which anti-aging effects of genotype were significantly larger in old mice than in young mice; red denotes phenotypes in which anti-aging effects of genotype were significantly larger in young mice than in old mice. **f**, ASPs accentuated by genotype. **g**, Phenotypes featuring a main effect of *Ghrhr^lit/lit^* genotype and/or a genotype × age interaction but not a main effect of age; blue dots denote phenotypes in which the genotype effect did not differ significantly between young and old mice. Yellow denotes phenotypes in which the genotype effect size differed significantly between young and old mice. **d**, all phenotypes shown in **e**-**g** collapsed into one panel. ICC = intraclass correlation. For further details, see **Supplementary Data 6**.

**Extended Data Figure 15: Anti-aging effects induced by a hypomorphic mTOR mutation; analysis restricted to tumor-free mice**. **a**, Principal component analysis of deep phenotyping data (number of mice: young WT, n=27; young *mTOR^KI/KI^*, n=21; old WT, n=18; old *mTOR^KI/KI^*, n=19). **b,** Venn diagram shows the number of age-sensitive phenotypes, genotype-sensitive phenotypes, phenotypes with a genotype × age interaction and their intersection. **c**, Sunburst chart shows the number of age-sensitive phenotypes either unaltered (in grey), counteracted (in green) or accentuated (in magenta) by the *mTOR^KI/KI^* mutation. For age-sensitive phenotypes counteracted by the *mTOR^KI/KI^* mutation, the inner ring shows the proportion of phenotypes with a main effect of genotype (in dark green), a genotype × age interaction (in violet) or both a main effect and an interaction (in yellow). The outer ring shows when changes in the corresponding ASPs were first detected based on data available from our baseline study. **d**–**g**, Scatter plots show the effect size of *mTOR^KI/KI^* genotype in young mice plotted vs. the effect size of *mTOR^KI/KI^* genotype in old mice for different sets of phenotypes: **e**, ASPs counteracted by genotype via a main effect and/or an interaction (i.e., corresponding to the central green section of the sunburst chart in **c**); green dots denote phenotypes in which genotype effects in young and old mice did not differ significantly; orange denotes phenotypes in which anti-aging effects of genotype were significantly larger in old mice than in young mice; red denotes phenotypes in which anti-aging effects of genotype were significantly larger in young mice than in old mice. **f**, ASPs accentuated by genotype. **g**, Phenotypes featuring a main effect of *mTOR^KI/KI^* genotype and/or a genotype × age interaction but not a main effect of age; blue dots denote phenotypes in which the genotype effect did not differ significantly between young and old mice. Yellow denotes phenotypes in which the genotype effect size differed significantly between young and old mice. **d**, all phenotypes shown in **e**-**g** collapsed into one panel. ICC = intraclass correlation. For further details, see **Supplementary Data 7**.

**Extended Data Figure 16: Anti-aging effects induced by every-other-day fasting; analysis restricted to tumor-free mice**. **a**, Principal component analysis of deep phenotyping data (number of mice: young AL, n=16; young IF, n=16; old AL, n=22; old IF, n=22). **b,** Venn diagram shows the number of age-sensitive phenotypes, diet-sensitive phenotypes, phenotypes with a diet × age interaction and their intersection. **c**, Sunburst chart shows the number of age-sensitive phenotypes either unaltered (in grey), counteracted (in green) or accentuated (in magenta) by IF. For age-sensitive phenotypes counteracted by IF, the inner ring shows the proportion of phenotypes with a main effect of diet (in dark green), a diet × age interaction (in violet) or both a main effect and an interaction (in yellow). The outer ring shows when changes in the corresponding ASPs were first detected based on data available from our baseline study. **d**–**g**, Scatter plots show the effect size of IF in young mice plotted vs. the effect size of IF in old mice for different sets of phenotypes: **e**, ASPs counteracted by diet via a main effect and/or an interaction (i.e., corresponding to the central green section of the sunburst chart in **c**); green dots denote phenotypes in which diet effects in young and old mice did not differ significantly; orange denotes phenotypes in which anti-aging effects of diet were significantly larger in old mice than in young mice; red denotes phenotypes in which anti-aging effects of diet were significantly larger in young mice than in old mice. **f**, ASPs accentuated by diet. **g**, Phenotypes featuring a main effect of diet and/or a diet × age interaction but not a main effect of age; blue dots denote phenotypes in which the diet effect did not differ significantly between young and old mice. Yellow denotes phenotypes in which the diet effect size differed significantly between young and old mice. **d**, all phenotypes shown in **e**-**g** collapsed into one panel. ICC = intraclass correlation. For further details, see **Supplementary Data 9**.

**Extended Data Figure 17: Gating strategy of peripheral blood FACS analysis.**

**Extended Data Figure 18: Gating strategy of peripheral blood FACS analysis.**

## References

1. Tacutu, R., et al. Human Ageing Genomic Resources: new and updated databases. Nucleic Acids Res 46, D1083–D1090 (2018).

2. Barardo, D., et al. The DrugAge database of aging-related drugs. Aging Cell 16, 594–597 (2017).

3. Miller, R.A. Biology of Aging and Longevity. in Hazzard’s Geriatric Medicine and Gerontology (eds. Halter, J.B., et al.) (McGraw Hill, 2009).

4. Lopez-Otin, C., Blasco, M.A., Partridge, L., Serrano, M. & Kroemer, G. The hallmarks of aging. Cell 153, 1194–1217 (2013).

5. Blackwell, B.N., Bucci, T.J., Hart, R.W. & Turturro, A. Longevity, body weight, and neoplasia in ad libitum-fed and diet-restricted C57BL6 mice fed NIH-31 open formula diet. Toxicol Pathol 23, 570–582 (1995).

6. Pettan-Brewer, C. & Treuting, P.M. Practical pathology of aging mice. Pathobiology of Aging & Age-related Diseases 1, 7202 (2011).

7. Brayton, C.F., Treuting, P.M. & Ward, J.M. Pathobiology of aging mice and GEM: background strains and experimental design. Vet Pathol 49, 85–105 (2012).

8. Lipman, R., Galecki, A., Burke, D.T. & Miller, R.A. Genetic loci that influence cause of death in a heterogeneous mouse stock. J Gerontol A Biol Sci Med Sci 59, 977–983 (2004).

9. Miller, R.A., et al. Rapamycin, but not resveratrol or simvastatin, extends life span of genetically heterogeneous mice. J Gerontol A Biol Sci Med Sci 66, 191–201 (2011).

10. Xie, K., et al. Every-other-day feeding extends lifespan but fails to delay many symptoms of aging in mice. Nat Commun 8, 155 (2017).

11. Rose, M.R. Evolutionary biology of aging, (Oxford University Press, Oxford, 1991).

12. Rockstein, M., Chesky, J.A. & Sussman, M. Comparative biology and evolution of aging. in Handbook of the biology of aging 3-34 (Van Nostrand Reinhold Company, New York, 1977).

13. Aspinall, R. Aging of the Organs and Systems, (Kluwer Academic Publishers, 2003).

14. Abdulla, A. & Rai, G.S. The biology of ageing and its clinical implications, (Radcliffe Publishing, London, 2013).

15. Freund, A. Untangling Aging Using Dynamic, Organism-Level Phenotypic Networks. Cell Syst 8, 172–181 (2019).

16. Neff, F., et al. Rapamycin extends murine lifespan but has limited effects on aging. J Clin Invest 123, 3272–3291 (2013).

17. Bellantuono, I., et al. A toolbox for the longitudinal assessment of healthspan in aging mice. Nat Protoc 15, 540–574 (2020).

18. Ehninger, D., Neff, F. & Xie, K. Longevity, aging and rapamycin. Cell Mol Life Sci 71, 4325–4346 (2014).

19. Richardson, A. & McCarter, R. Mechanism of food restriction: change of rate or change of set point. in The potential for nutritional modulation of aging processes (eds. Ingram, D.K., Baker, G.T. & Shock, N.W.) 177–192 (Food & Nutrition Press, Inc., 1992).

20. Meszaros, L., Hoffmann, A., Wihan, J. & Winkler, J. Current Symptomatic and Disease-Modifying Treatments in Multiple System Atrophy. Int J Mol Sci 21(2020).

21. Hampel, H., et al. Biomarkers for Alzheimer’s disease: academic, industry and regulatory perspectives. Nat Rev Drug Discov 9, 560–574 (2010).

22. Cummings, J. & Fox, N. Defining Disease Modifying Therapy for Alzheimer’s Disease. J Prev Alzheimers Dis 4, 109–115 (2017).

23. Espay, A. & Stecher, B. Symptomatic vs. Disease-Modifying Therapies. in Brain Fables: The Hidden History of Neurodegenerative Diseases and a Blueprint to Conquer Them 87–93 (Cambridge University Press, 2020).

24. Lamming, D.W. Extending Lifespan by Inhibiting the Mechanistic Target of Rapamycin (mTOR). in Anti-aging Drugs: From Basic Research to Clinical Practice (ed. Vaiserman, A.M.) 352–375 (The Royal Society of Chemistry, 2017).

25. Zhang, S., et al. Constitutive reductions in mTOR alter cell size, immune cell development, and antibody production. Blood 117, 1228–1238 (2011).

26. Wu, J.J., et al. Increased mammalian lifespan and a segmental and tissue-specific slowing of aging after genetic reduction of mTOR expression. Cell Rep 4, 913–920 (2013).

27. Zhang, S., et al. B cell-specific deficiencies in mTOR limit humoral immune responses. J Immunol 191, 1692–1703 (2013).

28. Eicher, E.M. & Beamer, W.G. Inherited ateliotic dwarfism in mice. Characteristics of the mutation, little, on chromosome 6. J Hered 67, 87–91 (1976).

29. Flurkey, K., Papaconstantinou, J., Miller, R.A. & Harrison, D.E. Lifespan extension and delayed immune and collagen aging in mutant mice with defects in growth hormone production. Proc Natl Acad Sci U S A 98, 6736–6741 (2001).

30. Ward, D.D., et al. Association of retinal layer measurements and adult cognitive function: A population-based study. Neurology 95, e1144–e1152 (2020).

31. McCay, C.M., Crowell, M.F. & Maynard, L.A. The effect of retarded growth upon the length of life span and upon the ultimate body size. J Nutr 10, 63–79 (1935).

32. Goodrick, C.L., Ingram, D.K., Reynolds, M.A., Freeman, J.R. & Cider, N. Effects of intermittent feeding upon body weight and lifespan in inbred mice: interaction of genotype and age. Mech Ageing Dev 55, 69–87 (1990).

33. Someya, S., et al. Age-related hearing loss in C57BL/6J mice is mediated by Bak-dependent mitochondrial apoptosis. Proc Natl Acad Sci U S A 106, 19432–19437 (2009).

34. Henson, S.M. & Aspinall, R. Aging and the Immune System. in Aging of Organs and Systems (ed. Aspinall, R.) 225–242 (Kluwer Academic Publishers, 2003).

35. Linton, P.J. & Dorshkind, K. Age-related changes in lymphocyte development and function. Nat Immunol 5, 133–139 (2004).

36. Dorshkind, K., Montecino-Rodriguez, E. & Signer, R.A. The ageing immune system: is it ever too old to become young again? Nat Rev Immunol 9, 57–62 (2009).

37. Bonda, T.A., et al. Remodeling of the intercalated disc related to aging in the mouse heart. J Cardiol 68, 261–268 (2016).

38. Mason, J.W., et al. Electrocardiographic reference ranges derived from 79,743 ambulatory subjects. J Electrocardiol 40, 228–234 (2007).

39. Wilkinson, J.E., et al. Rapamycin slows aging in mice. Aging Cell (2012).

40. Harrison, D.E., et al. Rapamycin fed late in life extends lifespan in genetically heterogeneous mice. Nature 460, 392–395 (2009).

41. Bartke, A. Growth Hormone and Aging: Updated Review. World J Mens Health 37, 19–30 (2019).

42. Bartke, A. & Quainoo, N. Impact of Growth Hormone-Related Mutations on Mammalian Aging. Front Genet 9, 586 (2018).

43. Garcia, J.M., Merriam, G.R. & Kargi, A.Y. Growth Hormone in Aging. in Endotext (eds. Feingold, K.R., et al.) (South Dartmouth (MA), 2000).

44. Kim, S.S. & Lee, C.K. Growth signaling and longevity in mouse models. BMB Rep 52, 70–85 (2019).

45. Carrie, I., Debray, M., Bourre, J.M. & Frances, H. Age-induced cognitive alterations in OF1 mice. Physiol Behav 66, 651–656 (1999).

46. GTEx-Consortium. The GTEx Consortium atlas of genetic regulatory effects across human tissues. Science 369, 1318–1330 (2020).

47. Shan, T., et al. Adipocyte-specific deletion of mTOR inhibits adipose tissue development and causes insulin resistance in mice. Diabetologia 59, 1995–2004 (2016).

48. Selman, C. Dietary restriction and the pursuit of effective mimetics. Proc Nutr Soc 73, 260–270 (2014).

49. Speakman, J.R. & Mitchell, S.E. Caloric restriction. Mol Aspects Med 32, 159–221 (2011).

50. Bordner, K.A., et al. Parallel declines in cognition, motivation, and locomotion in aging mice: association with immune gene upregulation in the medial prefrontal cortex. Exp Gerontol 46, 643–659 (2011).

51. Sprott, R.L. & Eleftheriou, B.E. Open-field behavior in aging inbred mice. Gerontologia 20, 155–162 (1974).

52. Ikeno, Y., Bronson, R.T., Hubbard, G.B., Lee, S. & Bartke, A. Delayed occurrence of fatal neoplastic diseases in ames dwarf mice: correlation to extended longevity. J Gerontol A Biol Sci Med Sci 58, 291–296 (2003).

53. Alderman, J.M., et al. Neuroendocrine inhibition of glucose production and resistance to cancer in dwarf mice. Exp Gerontol 44, 26–33 (2009).

54. Mueller, L.D., Rauser, C.L. & Rose, M.R. Aging Stops: Late Life, Evolutionary Biology, and Gerontology. in Does Aging Stop? (Oxford University Press, New York, 2011).

55. Wang, S. & Albers, K.M. Behavioral and cellular level changes in the aging somatosensory system. Ann N Y Acad Sci 1170, 745–749 (2009).

56. Ford, Z.K., et al. Systemic growth hormone deficiency causes mechanical and thermal hypersensitivity during early postnatal development. IBRO Rep 6, 111–121 (2019).

57. Petr, M.A., et al. A cross-sectional study of functional and metabolic changes during aging through the lifespan in male mice. Elife 10(2021).

58. Yang, A.C., et al. Physiological blood-brain transport is impaired with age by a shift in transcytosis. Nature 583, 425–430 (2020).

59. Schaum, N., et al. Ageing hallmarks exhibit organ-specific temporal signatures. Nature 583, 596–602 (2020).

60. Tabula Muris, C. A single-cell transcriptomic atlas characterizes ageing tissues in the mouse. Nature 583, 590–595 (2020).

61. Ximerakis, M., et al. Single-cell transcriptomic profiling of the aging mouse brain. Nat Neurosci 22, 1696–1708 (2019).

62. Fischer, K.E., et al. A cross-sectional study of male and female C57BL/6Nia mice suggests lifespan and healthspan are not necessarily correlated. Aging (Albany NY) 8, 2370–2391 (2016).

63. Hayflick, L. When does aging begin? Res Aging 6, 99–103 (1984).

64. Papadopoli, D., et al. mTOR as a central regulator of lifespan and aging. F1000Res 8(2019).

65. Martineau, C.N., Brown, A.E.X. & Laurent, P. Multidimensional phenotyping predicts lifespan and quantifies health in Caenorhabditis elegans. PLoS Comput Biol 16, e1008002 (2020).

66. Zhang, W.B., et al. Extended Twilight among Isogenic C. elegans Causes a Disproportionate Scaling between Lifespan and Health. Cell Syst 3, 333–345 e334 (2016).

67. Rockwood, K. & Mitnitski, A. Frailty in relation to the accumulation of deficits. J Gerontol A Biol Sci Med Sci 62, 722–727 (2007).

68. Fried, L.P., et al. Frailty in older adults: evidence for a phenotype. J Gerontol A Biol Sci Med Sci 56, M146–156 (2001).

69. Bell, C.G., et al. DNA methylation aging clocks: challenges and recommendations. Genome Biol 20, 249 (2019).

70. Xie, K., et al. Epigenetic alterations in longevity regulators, reduced life span, and exacerbated aging-related pathology in old father offspring mice. Proc Natl Acad Sci U S A 115, E2348–E2357 (2018).

71. Franceschi, C. & Campisi, J. Chronic inflammation (inflammaging) and its potential contribution to age-associated diseases. J Gerontol A Biol Sci Med Sci 69 Suppl 1, S4–9 (2014).

72. Shavlakadze, T., et al. Age-Related Gene Expression Signature in Rats Demonstrate Early, Late, and Linear Transcriptional Changes from Multiple Tissues. Cell Rep 28, 3263–3273 e3263 (2019).

73. Mair, W., Goymer, P., Pletcher, S.D. & Partridge, L. Demography of dietary restriction and death in Drosophila. Science 301, 1731–1733 (2003).

74. Hughes, B.G. & Hekimi, S. Different Mechanisms of Longevity in Long-Lived Mouse and Caenorhabditis elegans Mutants Revealed by Statistical Analysis of Mortality Rates. Genetics 204, 905–920 (2016).

75. Hahm, J.H., et al. C. elegans maximum velocity correlates with healthspan and is maintained in worms with an insulin receptor mutation. Nat Commun 6, 8919 (2015).

76. Zhao, Y., et al. Two forms of death in ageing Caenorhabditis elegans. Nat Commun 8, 15458 (2017).

77. Podshivalova, K., Kerr, R.A. & Kenyon, C. How a Mutation that Slows Aging Can Also Disproportionately Extend End-of-Life Decrepitude. Cell Rep 19, 441–450 (2017).

78. Stroustrup, N., et al. The temporal scaling of Caenorhabditis elegans ageing. Nature 530, 103–107 (2016).

79. Cohen, A.A., Levasseur, M., Raina, P., Fried, L.P. & Fulop, T. Is Aging Biology Ageist? J Gerontol A Biol Sci Med Sci 75, 1653–1655 (2020).

80. Le Couteur, D.G. & Simpson, S.J. Adaptive senectitude: the prolongevity effects of aging. J Gerontol A Biol Sci Med Sci 66, 179–182 (2011).

81. Fuchs, H., et al. Mouse phenotyping. Methods 53, 120–135 (2011).

82. Gailus-Durner, V., et al. Systemic first-line phenotyping. Methods Mol Biol 530, 463–509 (2009).

83. Rogers, D.C., et al. Behavioral and functional analysis of mouse phenotype: SHIRPA, a proposed protocol for comprehensive phenotype assessment. Mamm Genome 8, 711–713 (1997).

84. Jones, B.J. & Roberts, D.J. A rotarod suitable for quantitative measurements of motor incoordination in naive mice. Naunyn Schmiedebergs Arch Exp Pathol Pharmakol 259, 211 (1968).

85. Schoensiegel, F., et al. High throughput echocardiography in conscious mice: training and primary screens. Ultraschall Med 32 Suppl 1, S124–129 (2011).

86. Roth, D.M., Swaney, J.S., Dalton, N.D., Gilpin, E.A. & Ross, J., Jr. Impact of anesthesia on cardiac function during echocardiography in mice. Am J Physiol Heart Circ Physiol 282, H2134–2140 (2002).

87. Fischer, M.D., et al. Noninvasive, in vivo assessment of mouse retinal structure using optical coherence tomography. PLoS One 4, e7507 (2009).

88. Schmucker, C. & Schaeffel, F. In vivo biometry in the mouse eye with low coherence interferometry. Vision Res 44, 2445–2456 (2004).

89. Prusky, G.T., Alam, N.M., Beekman, S. & Douglas, R.M. Rapid quantification of adult and developing mouse spatial vision using a virtual optomotor system. Invest Ophthalmol Vis Sci 45, 4611–4616 (2004).

90. Rathkolb, B., et al. Blood Collection from Mice and Hematological Analyses on Mouse Blood. Curr Protoc Mouse Biol 3, 101–119 (2013).

91. Weaver, J.L., Broud, D.D., McKinnon, K. & Germolec, D.R. Serial phenotypic analysis of mouse peripheral blood leukocytes. Toxicol Mech Methods 12, 95–118 (2002).

92. Roederer, M., Nozzi, J.L. & Nason, M.C. SPICE: exploration and analysis of post-cytometric complex multivariate datasets. Cytometry A 79, 167–174 (2011).

93. Baumgarth, N. & Roederer, M. A practical approach to multicolor flow cytometry for immunophenotyping. J Immunol Methods 243, 77–97 (2000).

94. Hou, Z., et al. A cost-effective RNA sequencing protocol for large-scale gene expression studies. Sci Rep 5, 9570 (2015).

95. Lawrence, M., Gentleman, R. & Carey, V. rtracklayer: an R package for interfacing with genome browsers. Bioinformatics 25, 1841–1842 (2009).

96. Love, M.I., Huber, W. & Anders, S. Moderated estimation of fold change and dispersion for RNA-seq data with DESeq2. Genome Biol 15, 550 (2014).

97. Li, Y., Tomko, R.J., Jr. & Hochstrasser, M. Proteasomes: Isolation and Activity Assays. Curr Protoc Cell Biol 67, 3 43 41-43 43 20 (2015).

98. Driver, A.S., Kodavanti, P.R. & Mundy, W.R. Age-related changes in reactive oxygen species production in rat brain homogenates. Neurotoxicol Teratol 22, 175–181 (2000).

99. Aziz, N.A., et al. Seroprevalence and correlates of SARS-CoV-2 neutralizing antibodies: Results from a population-based study in Bonn, Germany. medrxiv 2020.08.24.20181206.(2020).

100. Estrada, S., et al. FatSegNet: A fully automated deep learning pipeline for adipose tissue segmentation on abdominal dixon MRI. Magn Reson Med 83, 1471–1483 (2020).

101. Colao, A., et al. Bone loss is correlated to the severity of growth hormone deficiency in adult patients with hypopituitarism. J Clin Endocrinol Metab 84, 1919–1924 (1999).

102. Leone, S., et al. Increased pain and inflammatory sensitivity in growth hormone-releasing hormone (GHRH) knockout mice. Prostaglandins Other Lipid Mediat 144, 106362 (2019).

103. Berryman, D.E., et al. Two-year body composition analyses of long-lived GHR null mice. J Gerontol A Biol Sci Med Sci 65, 31–40 (2010).

104. Leone, S., et al. Behavioural phenotyping, learning and memory in young and aged growth hormone-releasing hormone-knockout mice. Endocr Connect 7, 924–931 (2018).

105. Miller, R.A., et al. Rapamycin-Mediated Lifespan Increase in Mice is Dose and Sex-Dependent and Appears Metabolically Distinct from Dietary Restriction. Aging Cell 13, 468–477 (2014).

106. Strong, R., et al. Rapamycin-mediated mouse lifespan extension: Late-life dosage regimes with sex-specific effects. Aging Cell 19, e13269 (2020)

