## Extended Data Figures 1-18 for "Deep Phenotyping and Lifetime Trajectories Reveal Limited Effects of Longevity Regulators on the Aging Process in C57BL/6J Mice"

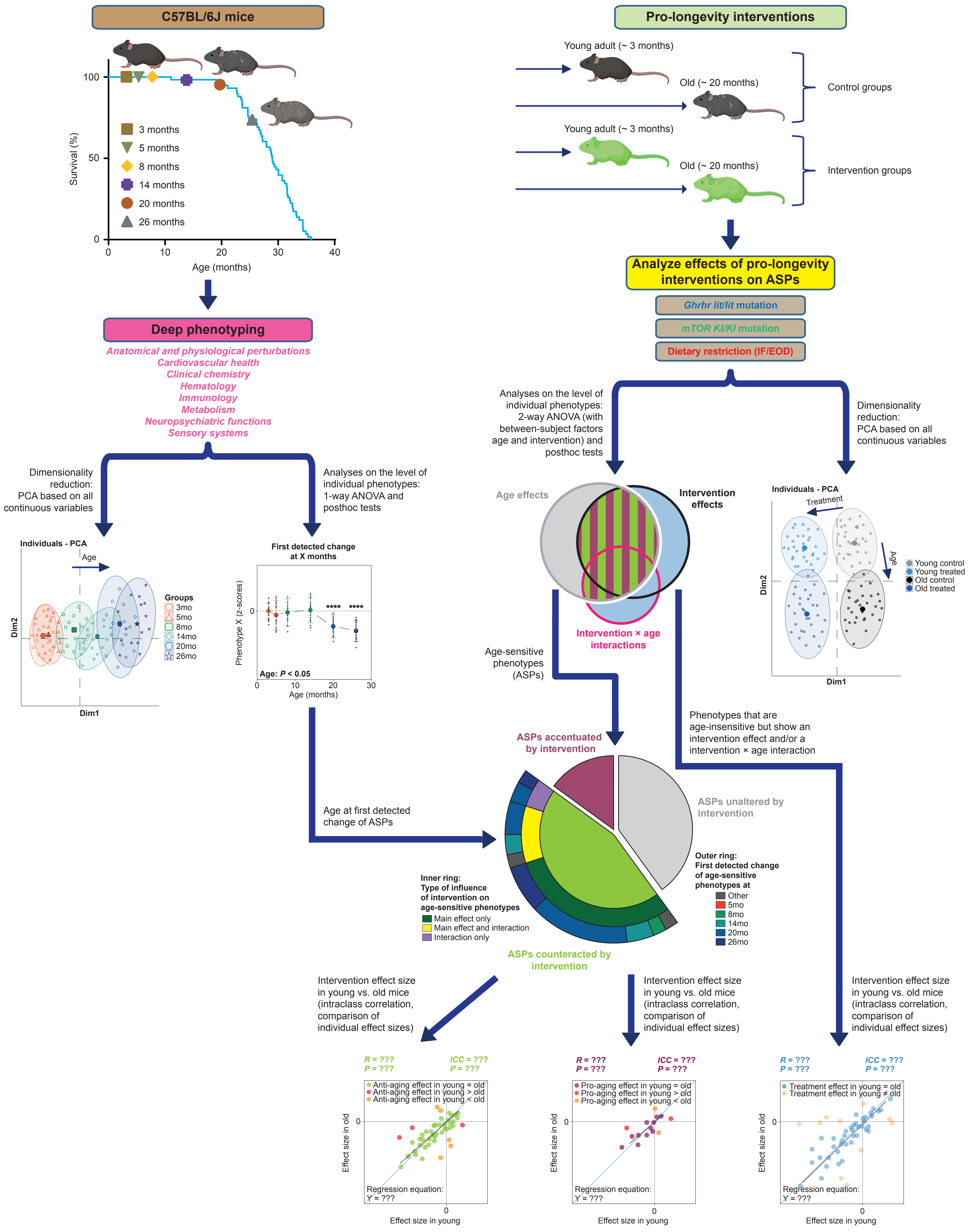


Extended Data Figure 1

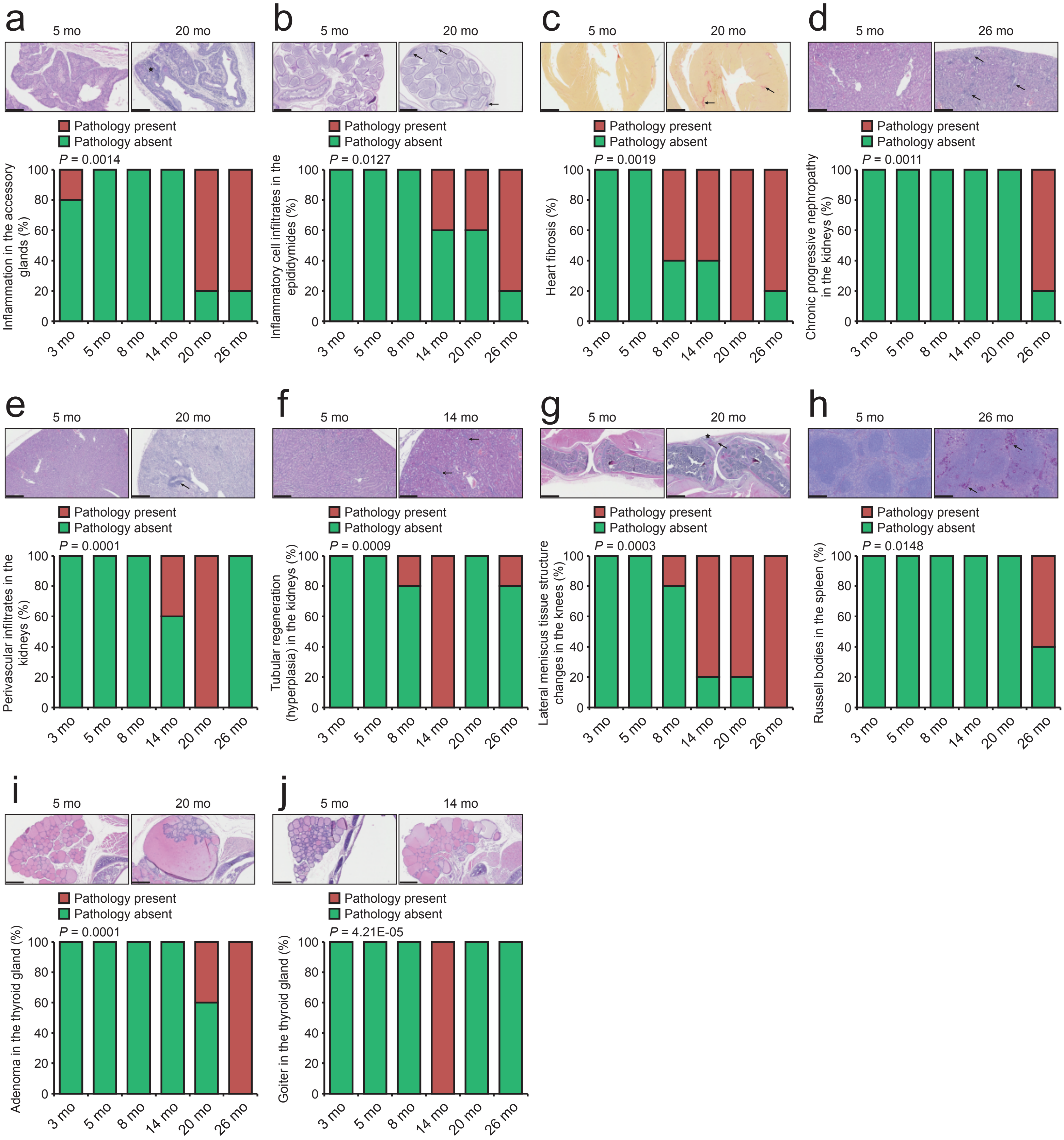


Extended Data Figure 2

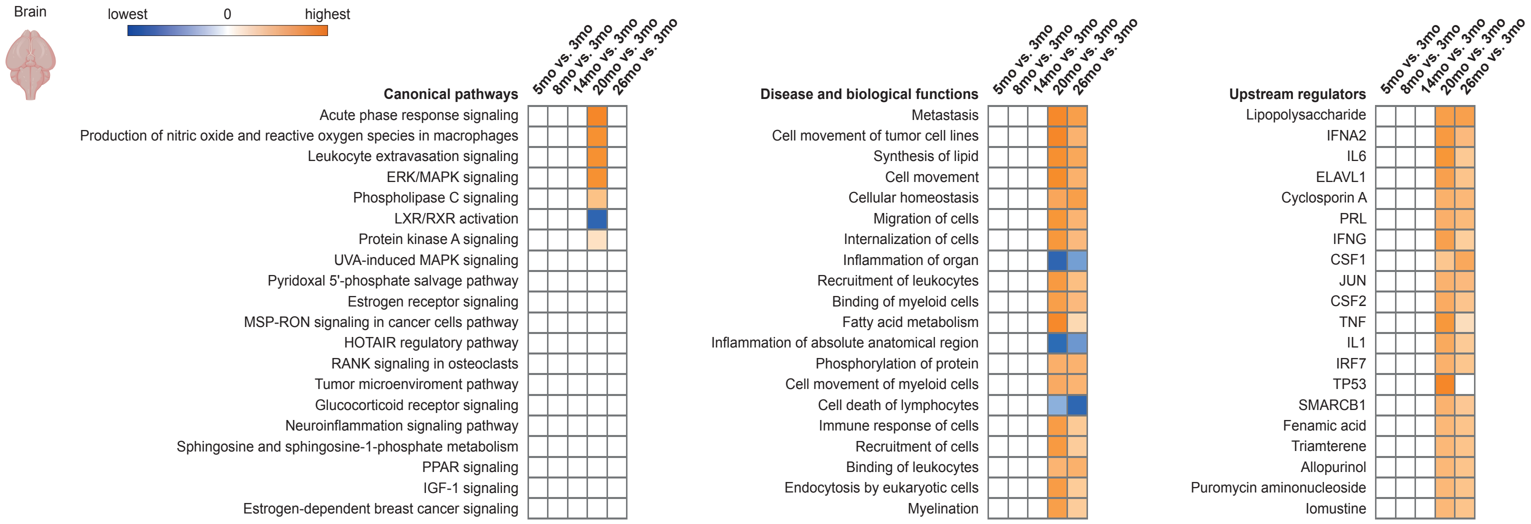


Extended Data Figure 3

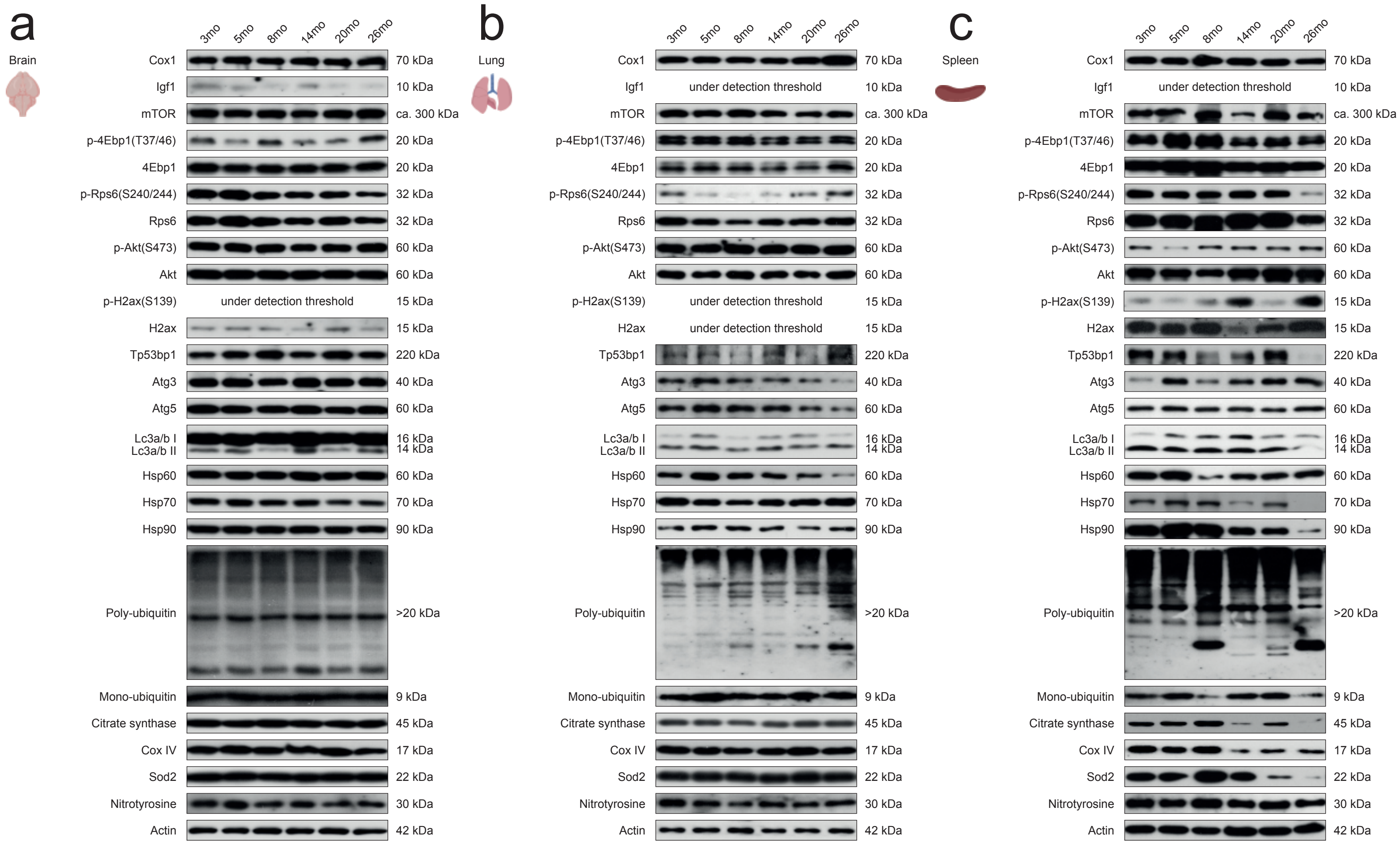


Extended Data Figure 4

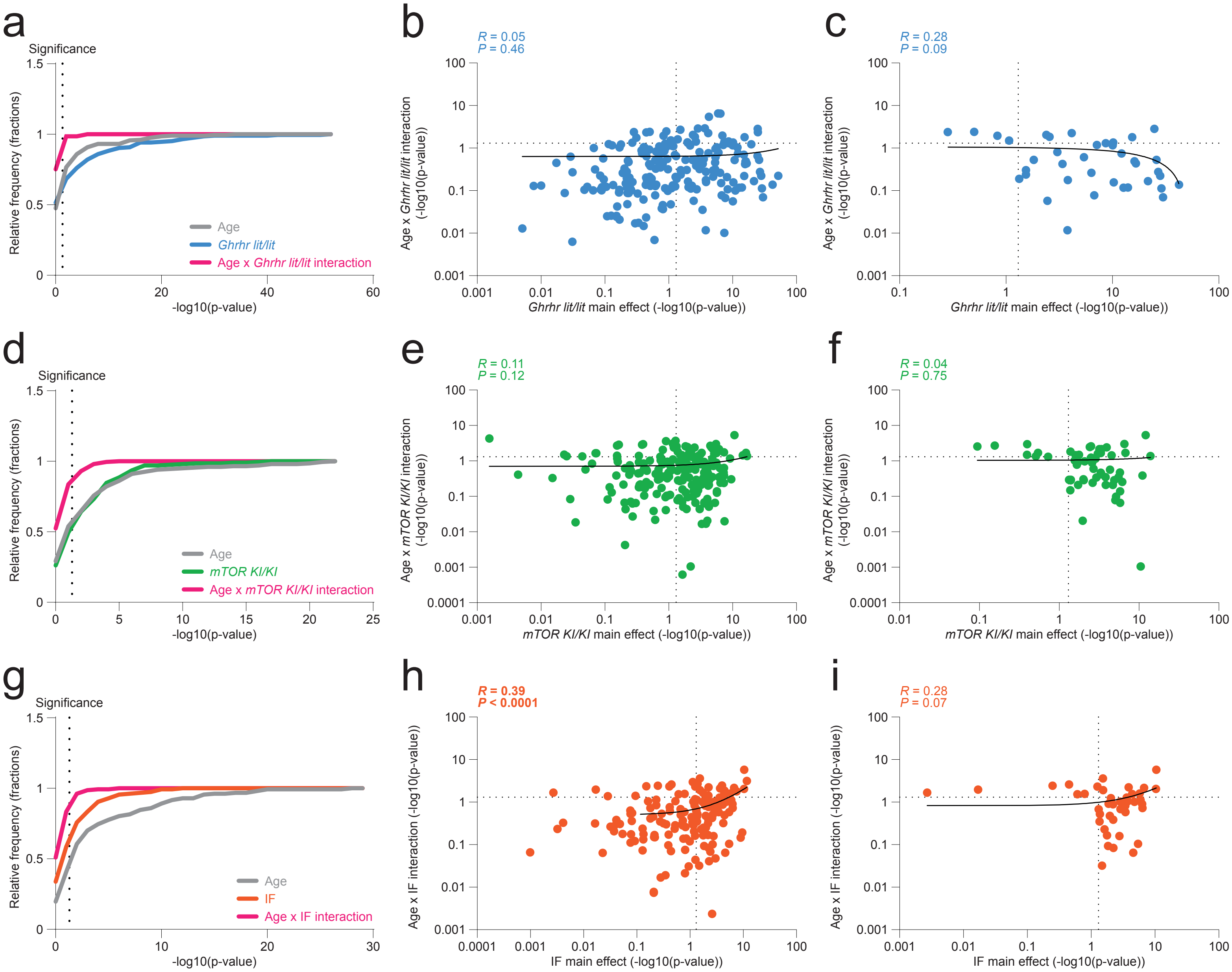


Extended Data Figure 5

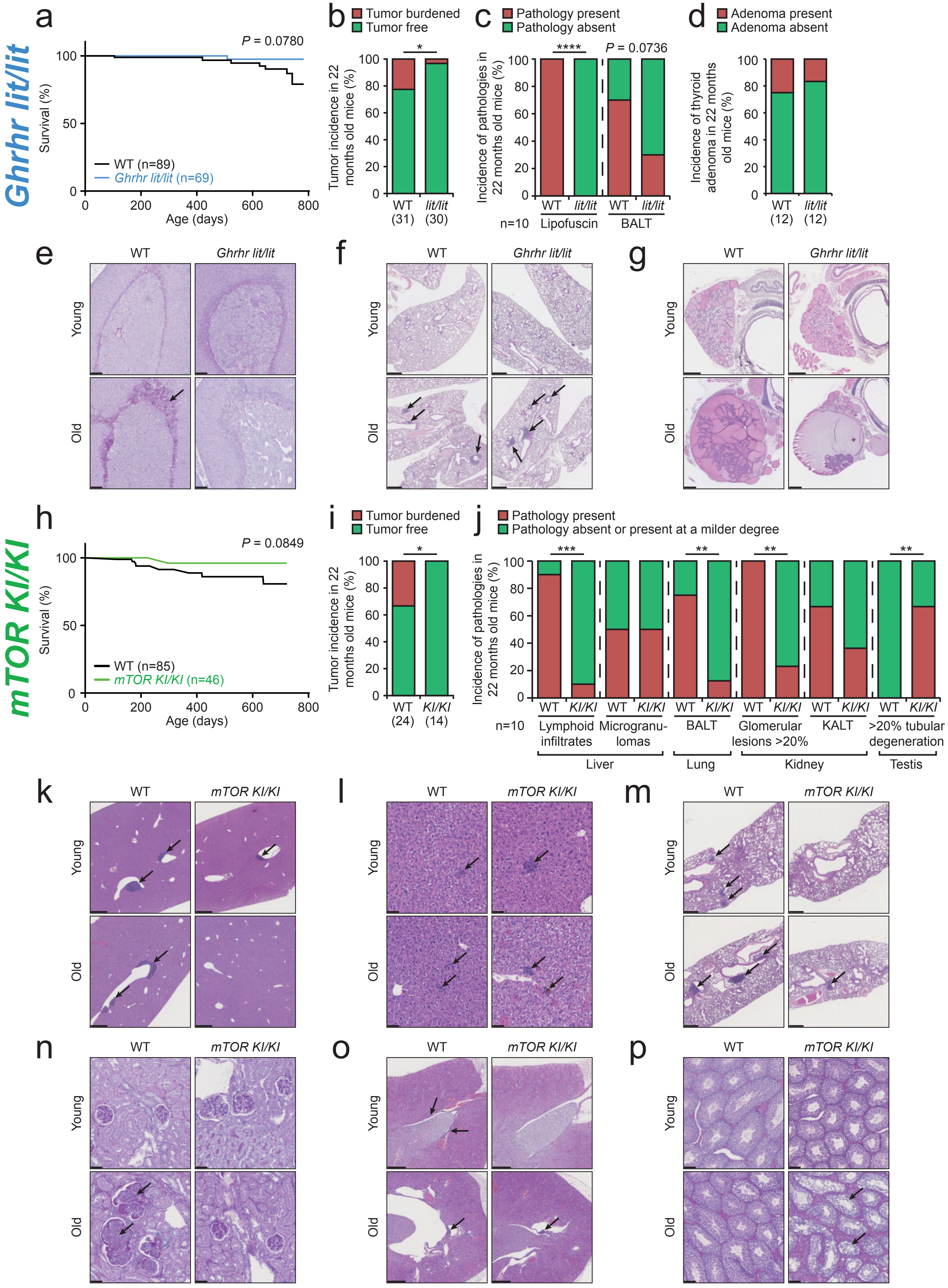


Extended Data Figure 6

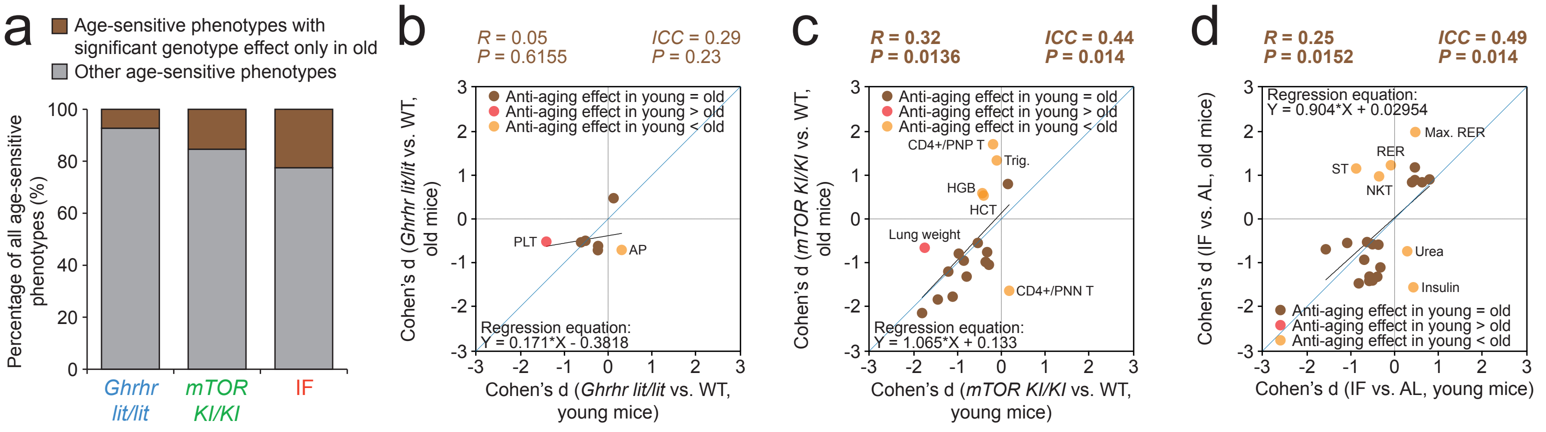


Extended Data Figure 7

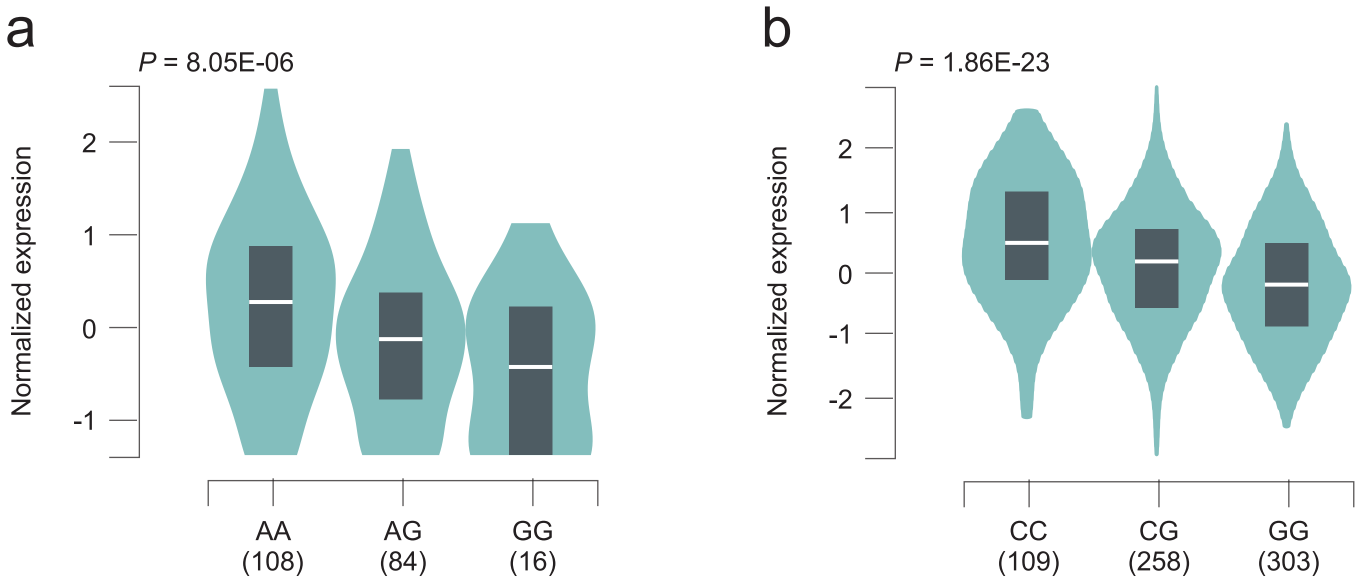


Extended Data Figure 8

a

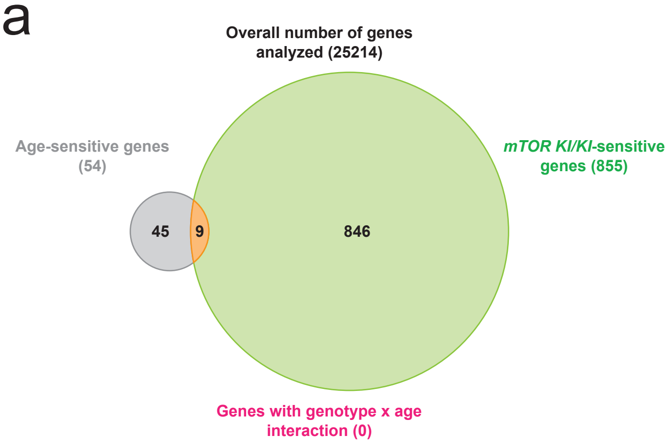


b

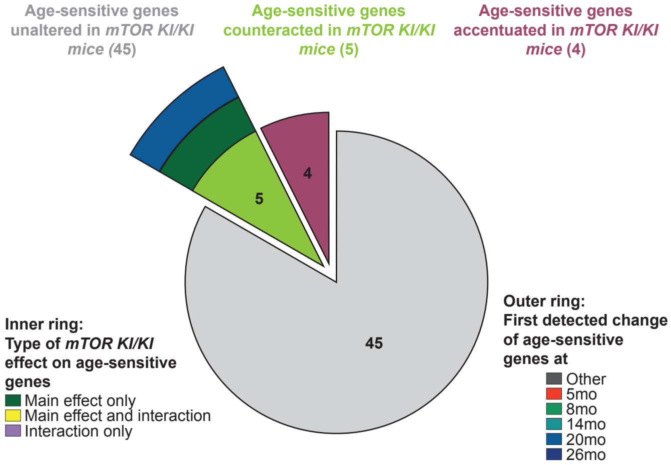


Extended Data Figure 9

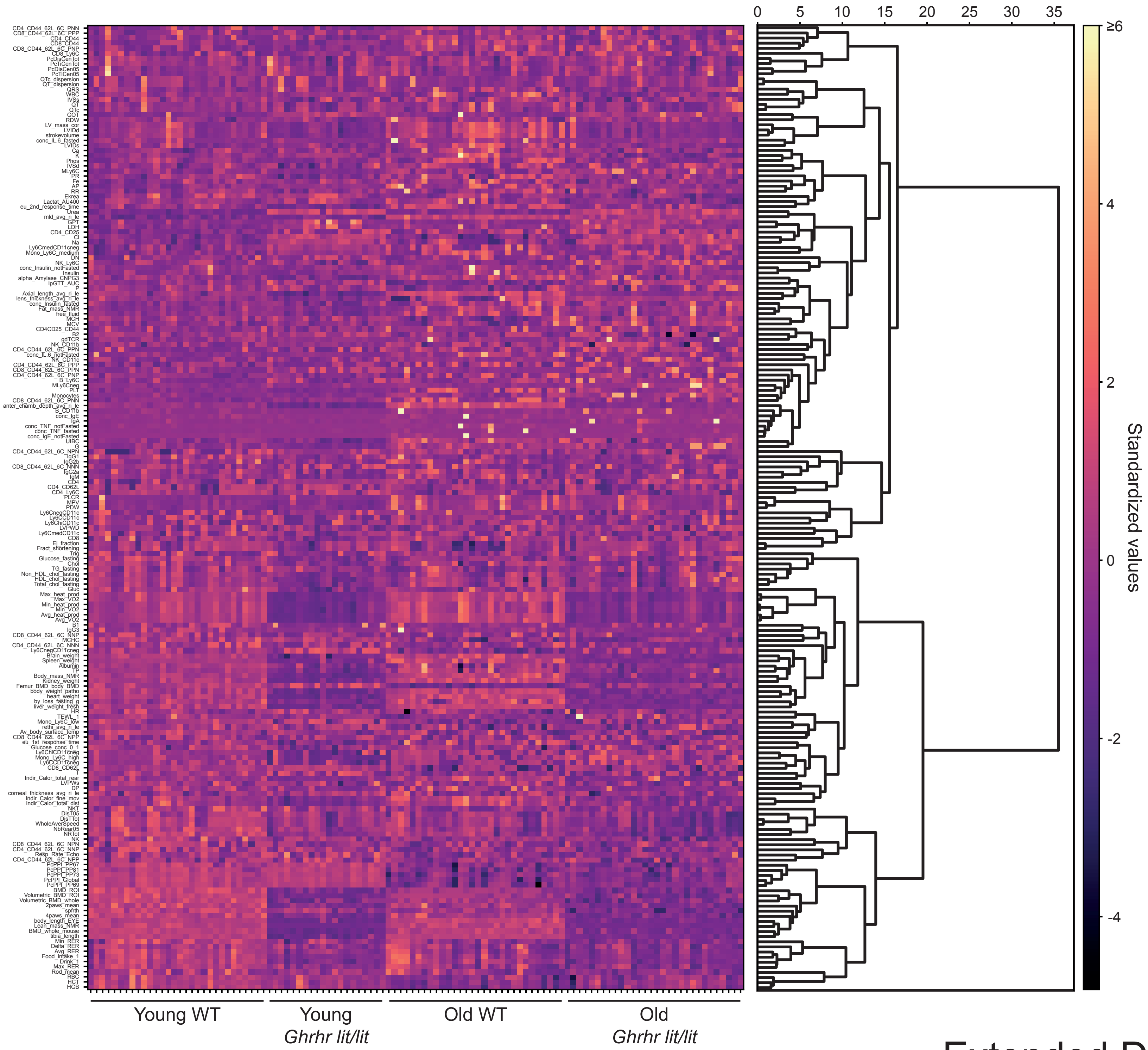


Extended Data Figure 10

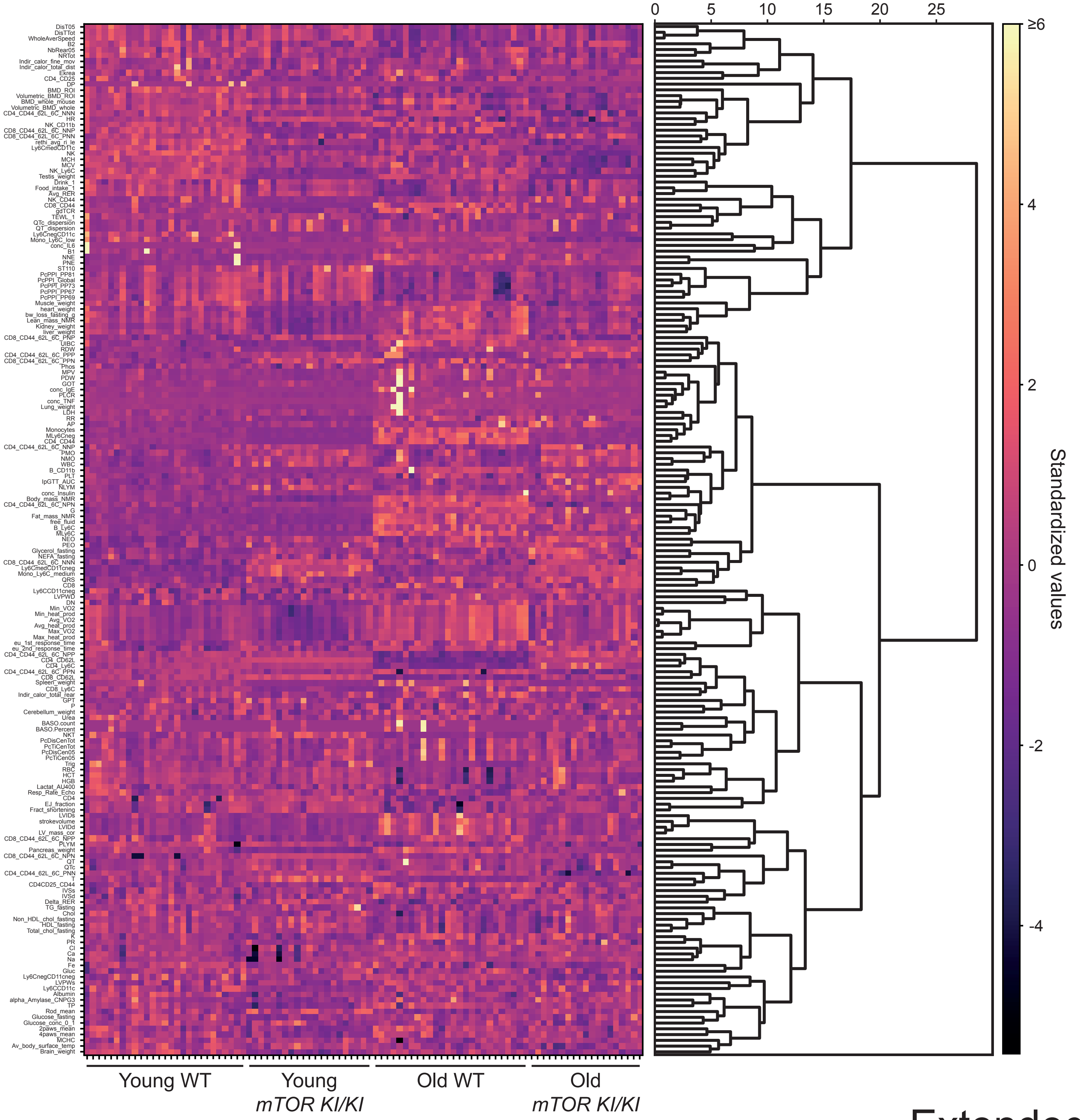


Extended Data Figure 11

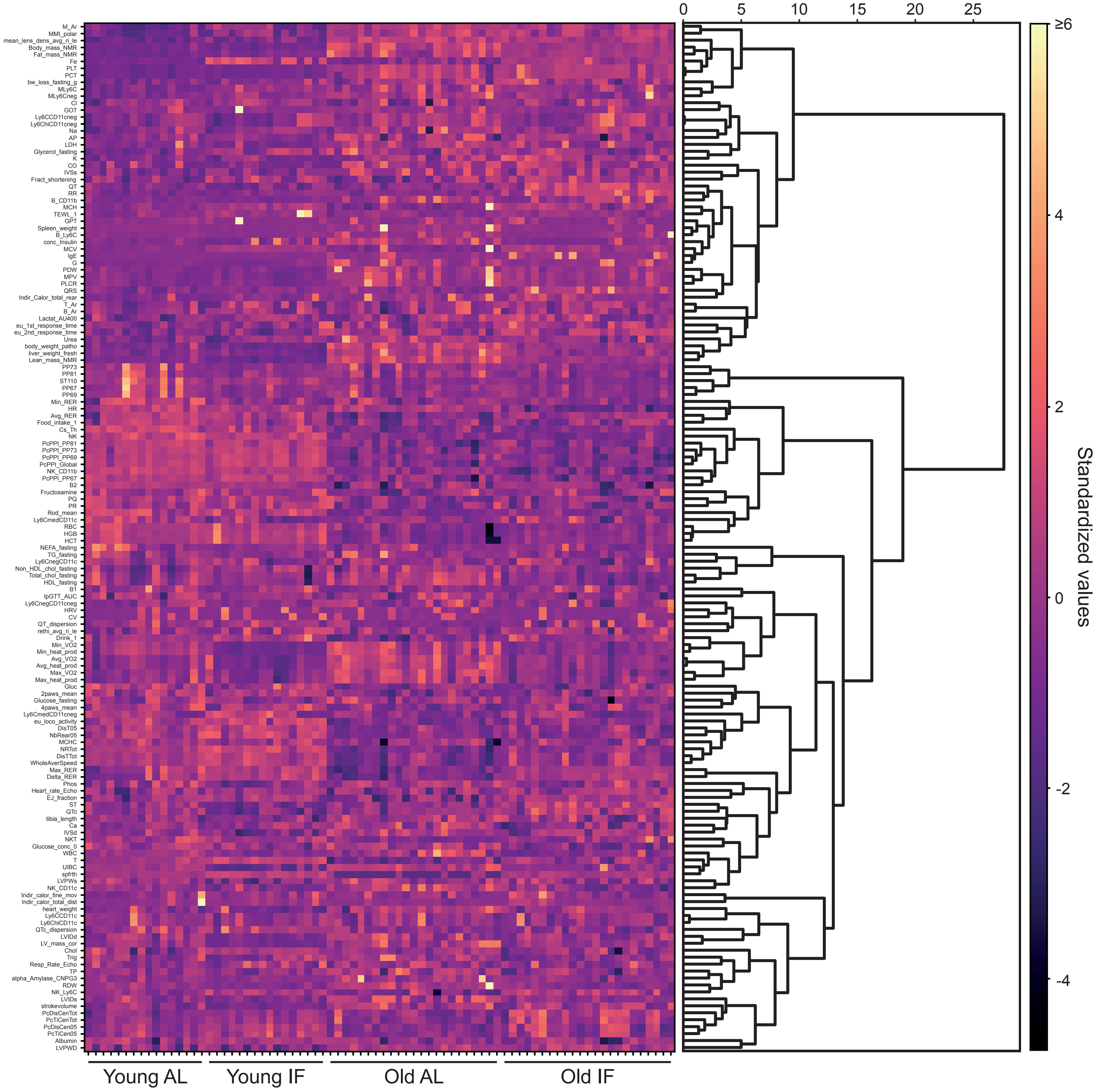


Extended Data Figure 12

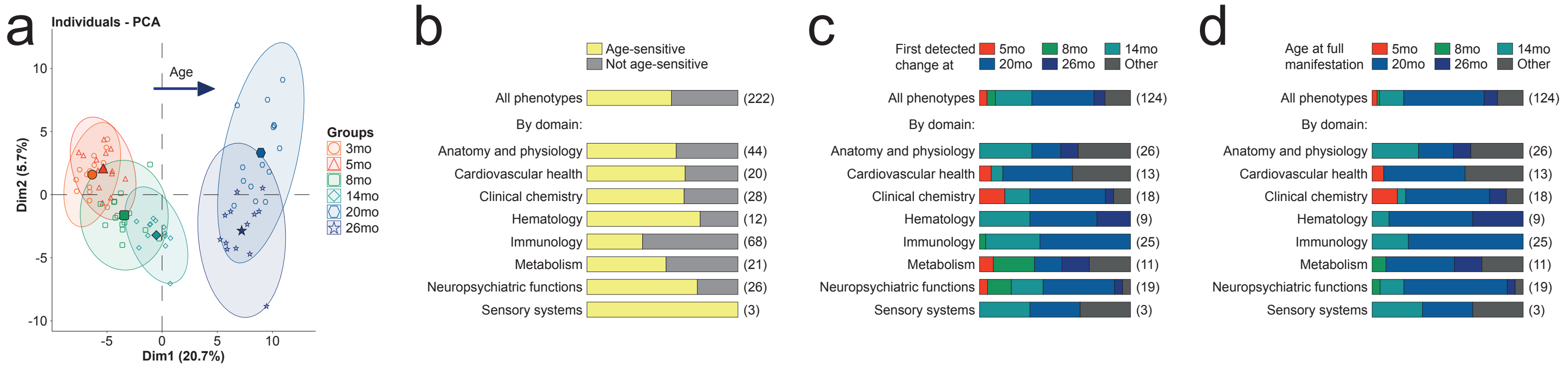


Extended Data Figure 13

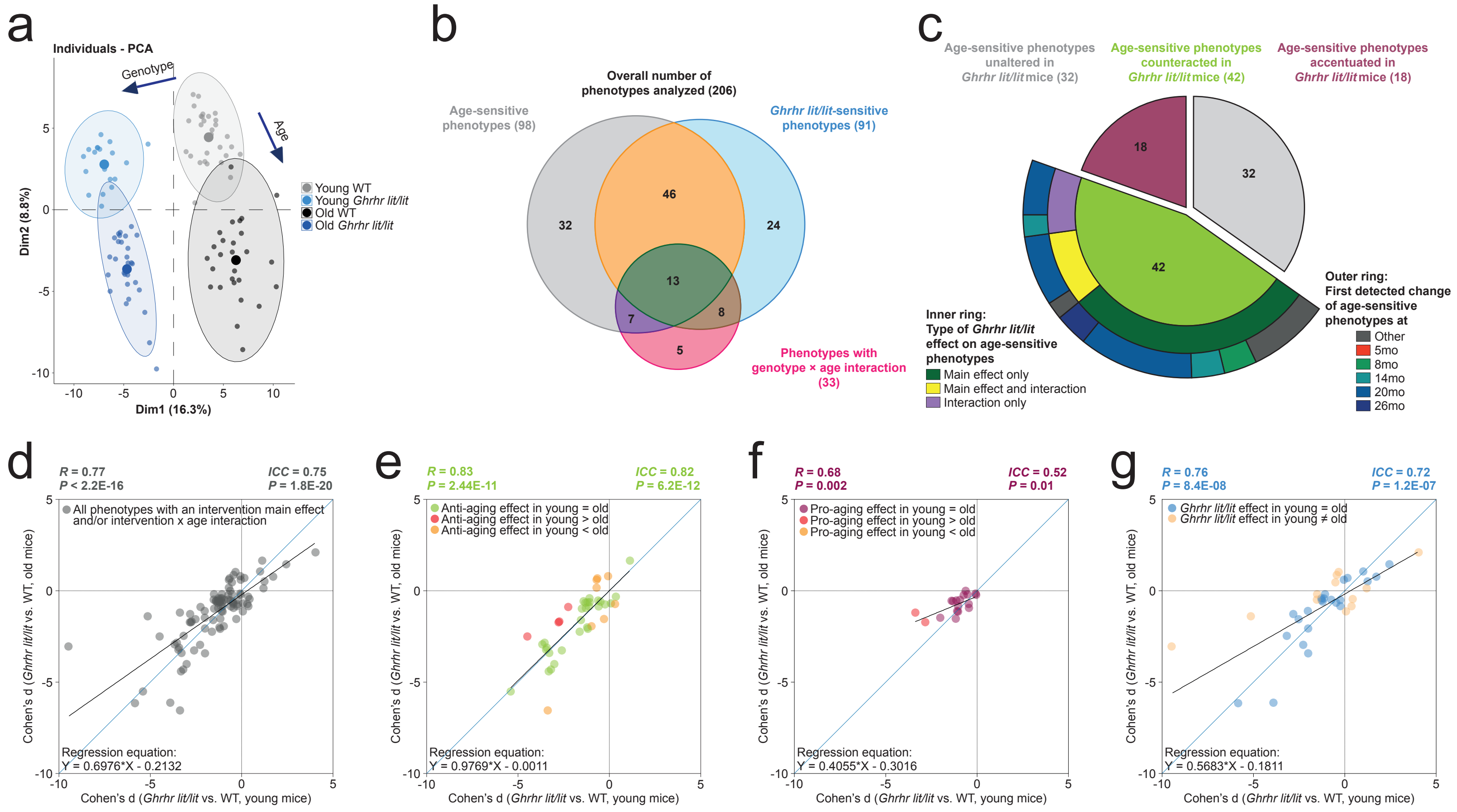


### Extended Data Figure 14

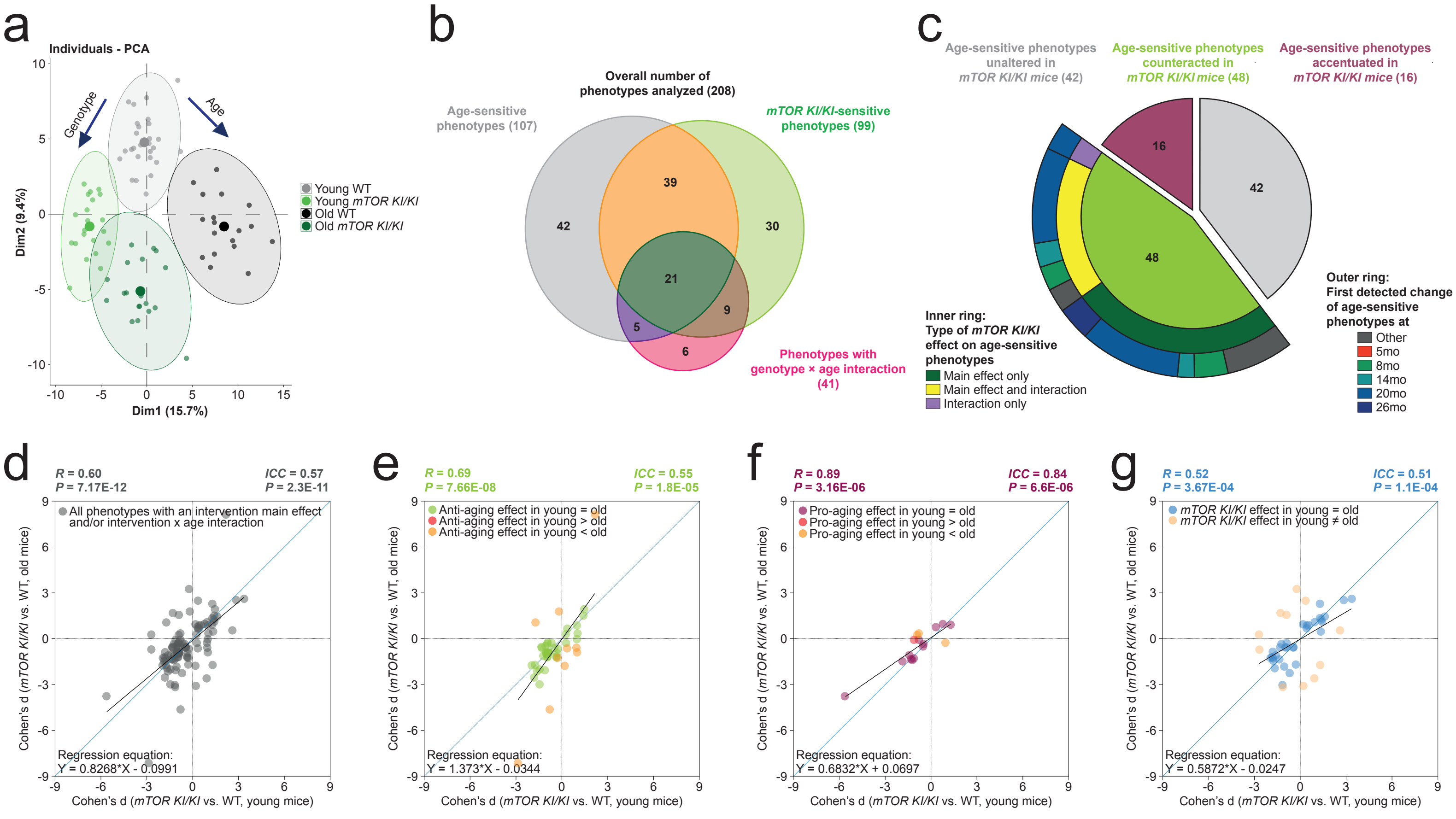


Extended Data Figure 15

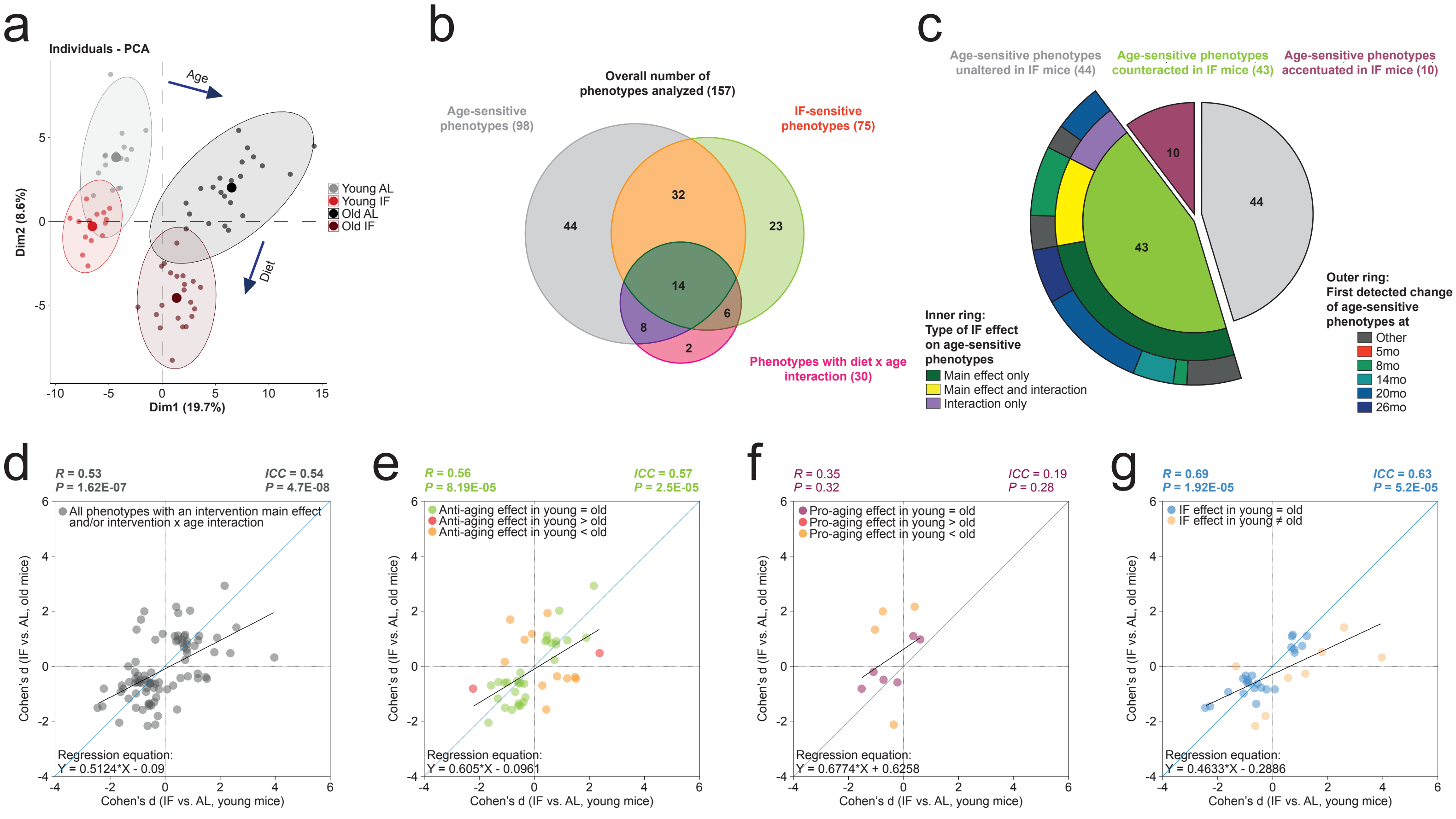


### Extended Data Figure 16

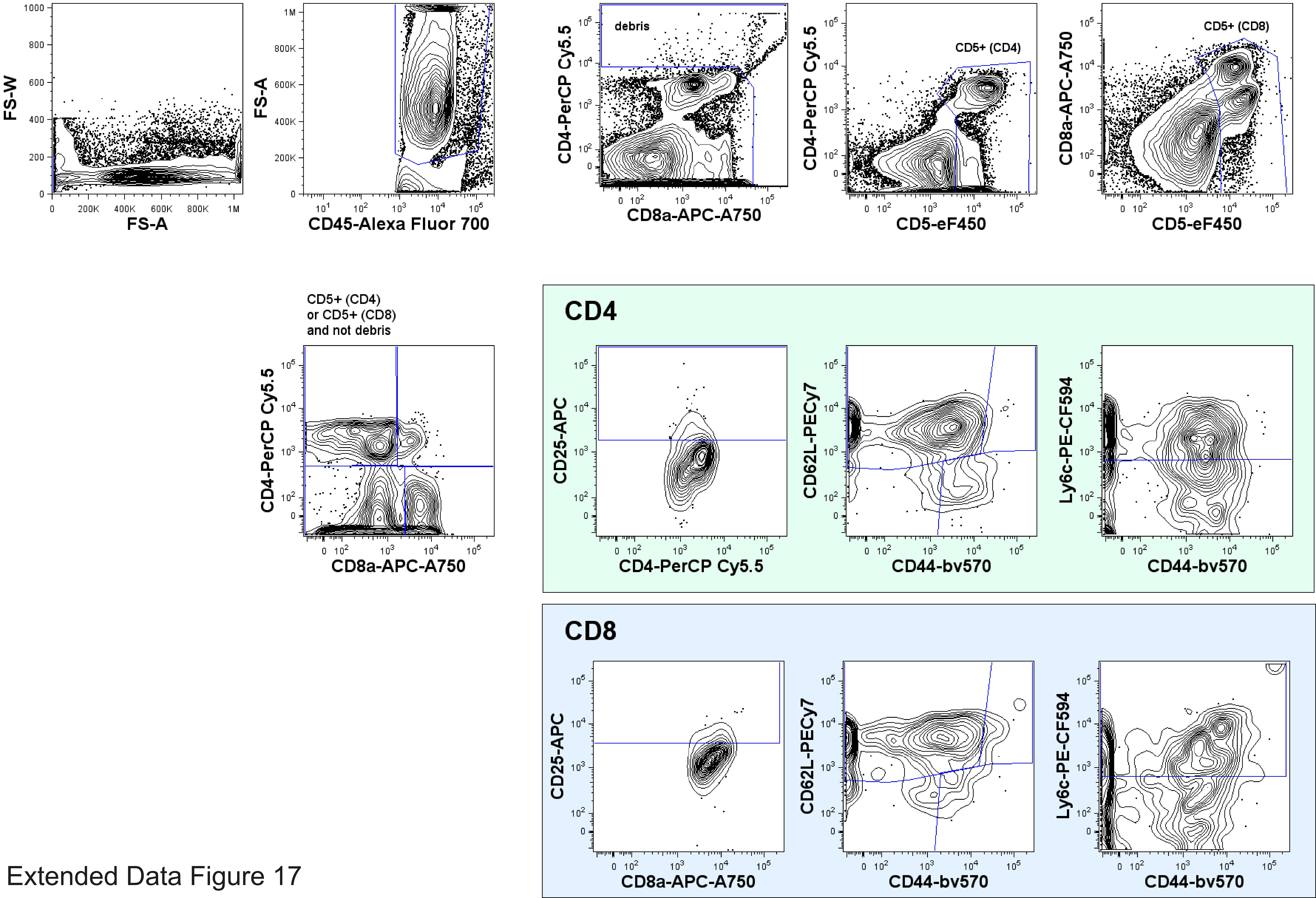


Extended Data Figure 17

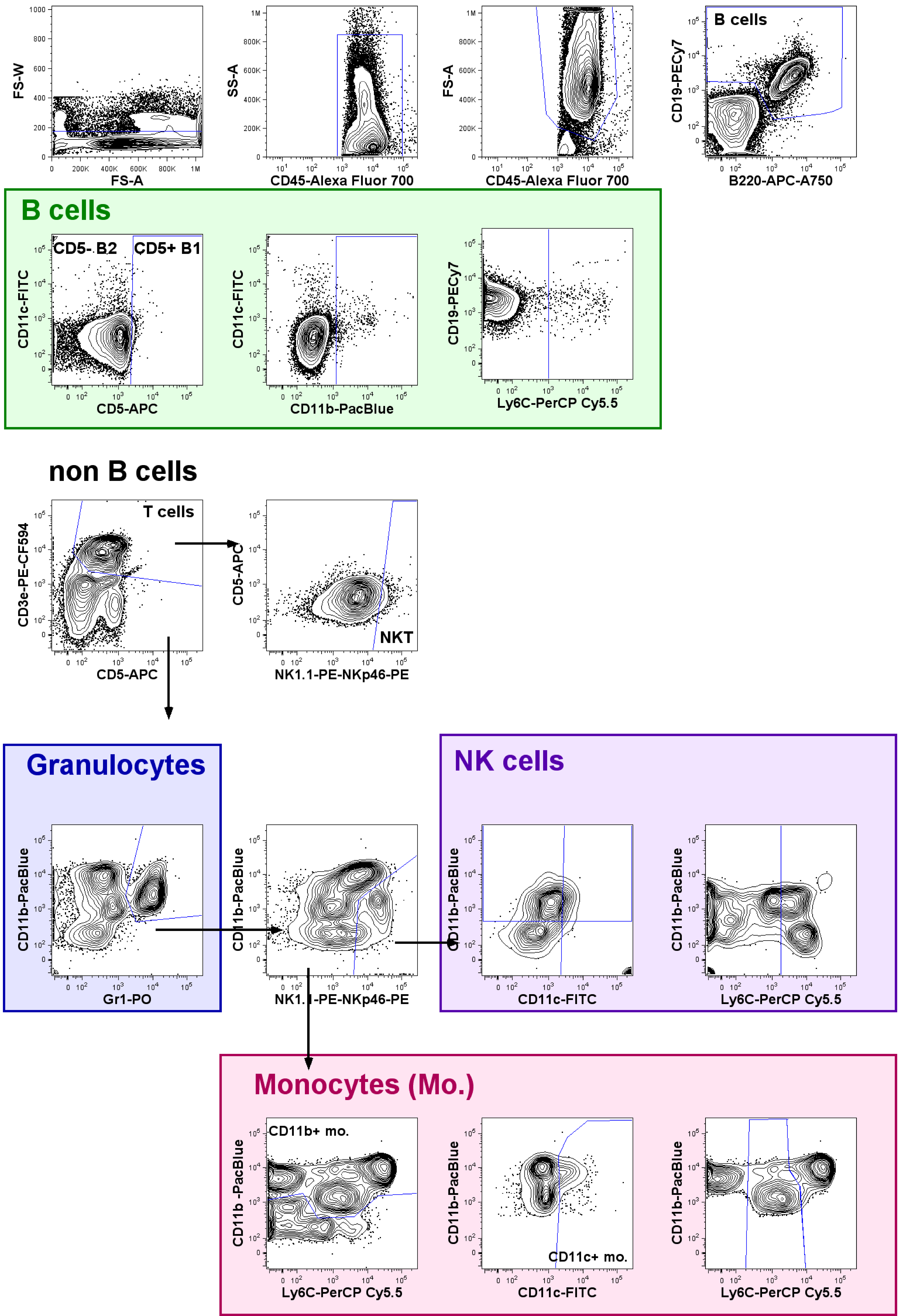


Extended Data Figure 18
