## Extended Data Tables 1-11 for "Deep Phenotyping and Lifetime Trajectories Reveal Limited Effects of Longevity Regulators on the Aging Process in C57BL/6J Mice"

**Extended Data Table 1. Deep phenotyping analyses in C57BL/6J wildtype mice to identify age-sensitive phenotypes including the age at first detected change relative to the young adult (3-month) baseline**

Age-sensitive phenotypes are denoted in bold.

| Domain | Test/method | Phenotype | P-value age | Age effect | First detected change at |
|---|---|---|---|---|---|
| **Anatomy and physiology** | **Body weight** | **Body weight** | **0.0237** | other | **other** |
| **Anatomy and physiology** | **Histology** | **Accessory gland inflammation** | **0.0014** | other | **other** |
| **Anatomy and physiology** | **Histology** | **Adrenal gland amyloid deposition** | **5.49E-06** | ↑ | **14 months** |
| **Anatomy and physiology** | **Histology** | **Adrenal gland lipofuscin pigments** | **0.0486** | ↓ | **14 months** |
| Anatomy and physiology | Histology | Adrenal gland pathology | n.s. | N/A | N/A |
| Anatomy and physiology | Histology | Adrenal subcapslar cell hyperplasia | n.s. | N/A | N/A |
| Anatomy and physiology | Histology | Epididymal epithelium vacuolated | n.s. | N/A | N/A |
| **Anatomy and physiology** | **Histology** | **Epididymal inflammatory cell infiltrates** | **0.0127** | ↑ | **26 months** |
| **Anatomy and physiology** | **Histology** | **Heart fibrosis** | **0.0019** | ↑ | **20 months** |
| **Anatomy and physiology** | **Histology** | **Kidney amyloid deposition** | **5.49E-06** | ↑ | **14 months** |
| **Anatomy and physiology** | **Histology** | **Kidney chronic progressive nephropathy** | **0.0011** | ↑ | **26 months** |
| **Anatomy and physiology** | **Histology** | **Kidney pathology** | **0.0002** | other | **other** |
| **Anatomy and physiology** | **Histology** | **Kidney perivascular infiltrates** | **0.0001** | ↑ | **20 months** |
| Anatomy and physiology | Histology | Kidney pyelonephritis | n.s. | N/A | N/A |
| **Anatomy and physiology** | **Histology** | **Kidney tubular regeneration** | **0.0009** | other | **other** |
| Anatomy and physiology | Histology | Knee inflammatory infiltrates around the joint | n.s. | N/A | N/A |
| **Anatomy and physiology** | **Histology** | **Knee lateral meniscus tissue structure** | **0.0003** | ↑ | **14 months** |
| **Anatomy and physiology** | **Histology** | **Knee pathology** | **2.06E-06** | ↑ | **14 months** |
| **Anatomy and physiology** | **Histology** | **Liver amyloid deposition** | **0.0003** | ↑ | **20 months** |
| Anatomy and physiology | Histology | Liver microgranuloma | n.s. | N/A | N/A |
| **Anatomy and physiology** | **Histology** | **Liver pathology** | **0.0264** | other | **other** |
| Anatomy and physiology | Histology | Liver perivascular infiltrates | n.s. | N/A | N/A |
| Anatomy and physiology | Histology | Lung alveolar histiocytosis | n.s. | N/A | N/A |
| Anatomy and physiology | Histology | Lung alveolar macrophages | n.s. | N/A | N/A |
| Anatomy and physiology | Histology | Lung bronchus-associated lymphoid tissue | n.s. | N/A | N/A |
| Anatomy and physiology | Histology | Lung cystic gland like infoldings | n.s. | N/A | N/A |
| **Anatomy and physiology** | **Histology** | **Lung pathology** | **0.0009** | other | **other** |
| Anatomy and physiology | Histology | Preputial gland duct dilation | n.s. | N/A | N/A |
| Anatomy and physiology | Histology | Seminiferous tubule degeneration | n.s. | N/A | N/A |
| **Anatomy and physiology** | **Histology** | **Spleen amyloid deposition** | **4.99E-07** | ↑ | **14 months** |
| **Anatomy and physiology** | **Histology** | **Spleen pathology** | **5.01E-06** | ↑ | **14 months** |
| **Anatomy and physiology** | **Histology** | **Spleen russle bodies** | **0.0148** | other | **other** |
| **Anatomy and physiology** | **Histology** | **Thyroid adenoma** | **0.0001** | ↑ | **26 months** |

| Domain | Test/method | Phenotype | P-value age | Age effect | First detected change at |
|---|---|---|---|---|---|
| Anatomy and physiology | Histology | Thyroid follicular cell hyperplasia | n.s. | N/A | N/A |
| **Anatomy and physiology** | **Histology** | **Thyroid goiter** | **4.21E-05** | other | **other** |
| Anatomy and physiology | Histology | Thyroid multifocal follicular cell hyperplasia | n.s. | N/A | N/A |
| **Anatomy and physiology** | **Histology** | **Thyroid pathology** | **1.29E-07** | ↑ | **14 months** |
| Anatomy and physiology | Organ weight | Brain weight | n.s. | N/A | N/A |
| **Anatomy and physiology** | **Organ weight** | **Heart weight** | **0.0286** | ↑ | **20 months** |
| **Anatomy and physiology** | **Organ weight** | **Kidney weight** | **4.08E-08** | ↑ | **20 months** |
| **Anatomy and physiology** | **Organ weight** | **Liver weight** | **0.0021** | other | **other** |
| **Anatomy and physiology** | **Organ weight** | **Spleen weight** | **9.39E-07** | ↑ | **14 months** |
| Anatomy and physiology | Organ weight | Tibia length | n.s. | N/A | N/A |
| Anatomy and physiology | TEWL | Transepidermal water loss | n.s. | N/A | N/A |
| **Cardiovascular health** | **Echocardiography** | **Corrected mass of the left ventricle** | **0.0129** | ↑ | **20 months** |
| **Cardiovascular health** | **Echocardiography** | **Diastolic septal wall thickness** | **0.0329** | other | **other** |
| **Cardiovascular health** | **Echocardiography** | **Ejection fraction** | **0.0247** | other | **other** |
| **Cardiovascular health** | **Echocardiography** | **Fractional shortening** | **0.0085** | other | **other** |
| **Cardiovascular health** | **Echocardiography** | **Left ventricular end-diastolic internal diameter** | **0.0072** | ↑ | **20 months** |
| **Cardiovascular health** | **Echocardiography** | **Left ventricular end-systolic internal diameter** | **0.0159** | other | **other** |
| **Cardiovascular health** | **Echocardiography** | **Respiration rate** | **0.0028** | ↓ | **5 months** |
| **Cardiovascular health** | **Echocardiography** | **Stroke volume** | **0.0143** | ↑ | **20 months** |
| Cardiovascular health | Echocardiography | Systolic septal wall thickness | n.s. | N/A | N/A |
| Cardiovascular health | Echocardiography | Thickness of the left ventricle posterior wall during diastole | n.s. | N/A | N/A |
| Cardiovascular health | Echocardiography | Thickness of the left ventricle posterior wall during systole | n.s. | N/A | N/A |
| Cardiovascular health | Electrocardiography | Corrected duration of the QT interval | n.s. | N/A | N/A |
| **Cardiovascular health** | **Electrocardiography** | **Duration of the P interval** | **0.0208** | other | **other** |
| Cardiovascular health | Electrocardiography | Duration of the PR interval | n.s. | N/A | N/A |
| **Cardiovascular health** | **Electrocardiography** | **Duration of the QRS interval** | **1.69E-05** | ↑ | **14 months** |
| **Cardiovascular health** | **Electrocardiography** | **Duration of the QT interval** | **0.0042** | ↑ | **20 months** |
| **Cardiovascular health** | **Electrocardiography** | **Duration of the RR interval** | **1.61E-06** | ↑ | **20 months** |
| **Cardiovascular health** | **Electrocardiography** | **Heart rate** | **7.16E-07** | ↓ | **20 months** |
| Cardiovascular health | Electrocardiography | QT dispersion | n.s. | N/A | N/A |
| Cardiovascular health | Electrocardiography | QTc dispersion | n.s. | N/A | N/A |
| **Clinical chemistry** | **Clinical chemistry** | **Albumin** | **2.76E-07** | ↓ | **14 months** |
| Clinical chemistry | Clinical chemistry | Alanine-aminotransferase | n.s. | N/A | N/A |
| **Clinical chemistry** | **Clinical chemistry** | **Alkaline phosphatase** | **8.99E-12** | ↑ | **20 months** |
| **Clinical chemistry** | **Clinical chemistry** | **Alpha amylase** | **1.05E-05** | ↑ | **20 months** |
| Clinical chemistry | Clinical chemistry | Aspartate-aminotransferase | n.s. | N/A | N/A |
| **Clinical chemistry** | **Clinical chemistry** | **Ca** | **3.32E-13** | ↑ | **20 months** |

| Domain | Test/method | Phenotype | P-value age | Age effect | First detected change at |
|---|---|---|---|---|---|
| Clinical chemistry | Clinical chemistry | Cholesterol | n.s. | N/A | N/A |
| **Clinical chemistry** | **Clinical chemistry** | **Cl** | **0.0076** | ↑ | **26 months** |
| **Clinical chemistry** | **Clinical chemistry** | **Creatinine** | **1.23E-08** | ↑ | **20 months** |
| Clinical chemistry | Clinical chemistry | Fe | n.s. | N/A | N/A |
| **Clinical chemistry** | **Clinical chemistry** | **Glucose** | **6.13E-06** | ↓ | **14 months** |
| **Clinical chemistry** | **Clinical chemistry** | **Glucose after fasting** | **7.86E-07** | ↓ | **20 months** |
| Clinical chemistry | Clinical chemistry | Glycerol after fasting | n.s. | N/A | N/A |
| **Clinical chemistry** | **Clinical chemistry** | **HDL-cholesterol after fasting** | **7.50E-06** | ↓ | **5 months** |
| **Clinical chemistry** | **Clinical chemistry** | **K** | **6.31E-07** | ↑ | **20 months** |
| Clinical chemistry | Clinical chemistry | Insulin | n.s. | N/A | N/A |
| Clinical chemistry | Clinical chemistry | Lactate | n.s. | N/A | N/A |
| Clinical chemistry | Clinical chemistry | Lactate dehydrogenase | n.s. | N/A | N/A |
| **Clinical chemistry** | **Clinical chemistry** | **Na** | **7.97E-05** | ↑ | **20 months** |
| Clinical chemistry | Clinical chemistry | Non-esterified fatty acids after fasting | n.s. | N/A | N/A |
| **Clinical chemistry** | **Clinical chemistry** | **Non HDL-cholesterol after fasting** | **3.35E-05** | ↓ | **5 months** |
| **Clinical chemistry** | **Clinical chemistry** | **Phosphate** | **0.0068** | other | **other** |
| **Clinical chemistry** | **Clinical chemistry** | **Triglycerides after fasting** | **4.85E-06** | ↓ | **14 months** |
| **Clinical chemistry** | **Clinical chemistry** | **Total cholesterol after fasting** | **2.27E-05** | ↓ | **5 months** |
| Clinical chemistry | Clinical chemistry | Total protein | n.s. | N/A | N/A |
| **Clinical chemistry** | **Clinical chemistry** | **Triglycerides** | **0.0046** | other | **other** |
| **Clinical chemistry** | **Clinical chemistry** | **Unsaturated iron binding capacity** | **0.0001** | ↑ | **20 months** |
| **Clinical chemistry** | **Clinical chemistry** | **Urea** | **5.34E-10** | ↑ | **20 months** |
| **Hematology** | **Hematology** | **Hematocrit** | **1.26E-19** | ↓ | **14 months** |
| **Hematology** | **Hematology** | **Hemoglobin concentration** | **9.86E-21** | ↓ | **14 months** |
| **Hematology** | **Hematology** | **Mean corpuscular hemoglobin content** | **0.0149** | ↓ | **20 months** |
| **Hematology** | **Hematology** | **Mean corpuscular hemoglobin concentration** | **0.0260** | ↓ | **26 months** |
| Hematology | Hematology | Mean corpuscular volume | n.s. | N/A | N/A |
| **Hematology** | **Hematology** | **Mean platelet volume** | **0.0072** | ↑ | **26 months** |
| Hematology | Hematology | Platelet distribution width | n.s. | N/A | N/A |
| **Hematology** | **Hematology** | **Platelet large cell ratio** | **0.0259** | ↑ | **26 months** |
| **Hematology** | **Hematology** | **Platelet count** | **3.80E-15** | ↑ | **14 months** |
| **Hematology** | **Hematology** | **Red blood cell count** | **2.76E-13** | ↓ | **20 months** |
| **Hematology** | **Hematology** | **Red blood cell distribution width** | **0.0148** | ↑ | **20 months** |
| **Hematology** | **Hematology** | **White blood cell count** | **0.0005** | ↑ | **14 months** |
| Immune system | ELISA | IgA | n.s. | N/A | N/A |
| Immune system | ELISA | IgE | n.s. | N/A | N/A |
| Immune system | ELISA | IgG1 | n.s. | N/A | N/A |

| Domain | Test/method | Phenotype | P-value age | Age effect | First detected change at |
|---|---|---|---|---|---|
| **Immune system** | **ELISA** | **IgG2a** | **0.0182** | ↑ | **20 months** |
| **Immune system** | **ELISA** | **IgG2b** | **0.0059** | ↑ | **20 months** |
| Immune system | ELISA | IgG3 | n.s. | N/A | N/A |
| **Immune system** | **ELISA** | **Interleukin-6** | **2.70E-07** | ↑ | **20 months** |
| **Immune system** | **ELISA** | **IgM** | **1.08E-05** | ↑ | **14 months** |
| **Immune system** | **ELISA** | **Tumor necrosis factor-α** | **3.49E-06** | ↑ | **20 months** |
| Immune system | Flow cytometry | $CD19^{+}B220^{+}/CD11b^{+}$ % of B cells | n.s. | N/A | N/A |
| Immune system | Flow cytometry | $CD19^{+}B220^{+}/Ly6C^{+}$ % of B cells | n.s. | N/A | N/A |
| Immune system | Flow cytometry | $CD19^{+}B220^{+}/CD5^{+}$ B1 cells % of all leukocytes | n.s. | N/A | N/A |
| **Immune system** | **Flow cytometry** | **$CD19^{+}B220^{+}/CD5^{-}$ B2 cells % of all leukocytes** | **0.0124** | ↓ | **20 months** |
| Immune system | Flow cytometry | $CD3^{+}CD5^{+}$ T cells % of all leukocytes | n.s. | N/A | N/A |
| **Immune system** | **Flow cytometry** | **$CD5^{+}/CD4^{+}$ % of T cells** | **9.90E-05** | ↓ | **14 months** |
| Immune system | Flow cytometry | $CD5^{+}/CD4^{+}CD25^{+}$ % of $CD4^{+}$ T cells | n.s. | N/A | N/A |
| Immune system | Flow cytometry | $CD5^{+}/CD4^{+}CD25^{+}CD44^{++}$ % of $CD5^{+}/CD4^{+}CD25^{+}$ T cells | n.s. | N/A | N/A |
| Immune system | Flow cytometry | $CD5^{+}/CD4^{+}CD44^{++}$ % of $CD4^{+}$ T cells | n.s. | N/A | N/A |
| **Immune system** | **Flow cytometry** | **$CD5^{+}/CD4^{+}CD44^{-}CD62L^{-}Ly6C^{-}$ % of $CD4^{+}$ T cells** | **6.80E-06** | ↑ | **20 months** |
| Immune system | Flow cytometry | $CD5^{+}/CD4^{+}CD44^{-}CD62L^{-}Ly6C^{+}$ % of $CD4^{+}$ T cells | n.s. | N/A | N/A |
| Immune system | Flow cytometry | $CD5^{+}/CD4^{+}CD44^{-}CD62L^{+}Ly6C^{-}$ % of $CD4^{+}$ T cells | n.s. | N/A | N/A |
| **Immune system** | **Flow cytometry** | **$CD5^{+}/CD4^{+}CD44^{-}CD62L^{+}Ly6C^{+}$ % of $CD4^{+}$ T cells** | **0.0435** | ↓ | **14 months** |
| **Immune system** | **Flow cytometry** | **$CD5^{+}/CD4^{+}CD44^{+}CD62L^{-}Ly6C^{-}$ % of $CD4^{+}$ T cells** | **2.99E-08** | ↑ | **14 months** |
| Immune system | Flow cytometry | $CD5^{+}/CD4^{+}CD44^{+}CD62L^{-}Ly6C^{+}$ % of $CD4^{+}$ T cells | n.s. | N/A | N/A |
| Immune system | Flow cytometry | $CD5^{+}/CD4^{+}CD44^{+}CD62L^{+}Ly6C^{-}$ % of $CD4^{+}$ T cells | n.s. | N/A | N/A |
| Immune system | Flow cytometry | $CD5^{+}/CD4^{+}CD44^{+}CD62L^{+}Ly6C^{+}$ % of $CD4^{+}$ T cells | n.s. | N/A | N/A |
| **Immune system** | **Flow cytometry** | **$CD5^{+}/CD4^{+}CD62L^{+}$ % of $CD4^{+}$ T cells** | **1.04E-07** | ↓ | **8 months** |
| **Immune system** | **Flow cytometry** | **$CD5^{+}/CD4^{+}Ly6C^{+}$ % of $CD4^{+}$ T cells** | **0.0008** | ↓ | **20 months** |
| **Immune system** | **Flow cytometry** | **$CD5^{+}/CD8^{+}$ % of T cells** | **0.0003** | ↑ | **14 months** |
| **Immune system** | **Flow cytometry** | **$CD5^{+}/CD8^{+}CD44^{+}$ % of $CD8^{+}$ T cells** | **1.91E-05** | ↑ | **14 months** |
| Immune system | Flow cytometry | $CD5^{+}/CD8^{+}CD44^{-}CD62L^{-}Ly6C^{-}$ % of $CD8^{+}$ T cells | n.s. | N/A | N/A |
| Immune system | Flow cytometry | $CD5^{+}/CD8^{+}CD44^{-}CD62L^{-}Ly6C^{+}$ % of $CD8^{+}$ T cells | n.s. | N/A | N/A |
| **Immune system** | **Flow cytometry** | **$CD5^{+}/CD8^{+}CD44^{-}CD62L^{+}Ly6C^{-}$ % of $CD8^{+}$ T cells** | **0.0022** | ↓ | **14 months** |
| **Immune system** | **Flow cytometry** | **$CD5^{+}/CD8^{+}CD44^{-}CD62L^{+}Ly6C^{+}$ % of $CD8^{+}$ T cells** | **0.0018** | ↓ | **14 months** |
| Immune system | Flow cytometry | $CD5^{+}/CD8^{+}CD44^{++}CD62L^{-}Ly6C^{-}$ % of $CD8^{+}$ T cells | n.s. | N/A | N/A |
| Immune system | Flow cytometry | $CD5^{+}/CD8^{+}CD44^{++}CD62L^{-}Ly6C^{+}$ % of $CD8^{+}$ T cells | n.s. | N/A | N/A |
| **Immune system** | **Flow cytometry** | **$CD5^{+}/CD8^{+}CD44^{++}CD62L^{+}Ly6C^{-}$ % of $CD8^{+}$ T cells** | **0.0290** | other | **other** |
| **Immune system** | **Flow cytometry** | **$CD5^{+}/CD8^{+}CD44^{++}CD62L^{+}Ly6C^{+}$ % of $CD8^{+}$ T cells** | **3.17E-07** | ↑ | **14 months** |
| Immune system | Flow cytometry | $CD5^{+}/CD8^{+}CD62L^{+}$ % of $CD8^{+}$ T cells | n.s. | N/A | N/A |
| **Immune system** | **Flow cytometry** | **$CD5^{+}/CD8^{+}Ly6C^{+}$ % of $CD8^{+}$ T cells** | **0.0003** | ↑ | **20 months** |

| Domain | Test/method | Phenotype | P-value age | Age effect | First detected change at |
|---|---|---|---|---|---|
| Immune system | Flow cytometry | CD5$^{+}$/CD4$^{-}$CD8$^{-}$ % of T cells | n.s. | N/A | N/A |
| Immune system | Flow cytometry | CD5$^{+}$/CD4$^{+}$CD8$^{+}$ % of T cells | n.s. | N/A | N/A |
| Immune system | Flow cytometry | CD5$^{+}$/gamma delta receptor$^{+}$ % of T cells | n.s. | N/A | N/A |
| **Immune system** | **Flow cytometry** | **CD11b$^{+}$Ly6G$^{+}$ granulocytes % of all leukocytes** | **8.20E-05** | ↑ | **20 months** |
| Immune system | Flow cytometry | CD11b$^{+}$/CD11c$^{+}$Ly6C$^{+}$ % of monocytes | n.s. | N/A | N/A |
| Immune system | Flow cytometry | CD11b$^{+}$/CD11c$^{-}$Ly6C$^{+}$ % of monocytes | n.s. | N/A | N/A |
| Immune system | Flow cytometry | CD11b$^{+}$/CD11c$^{+}$Ly6C$^{++}$ % of monocytes | n.s. | N/A | N/A |
| Immune system | Flow cytometry | CD11b$^{+}$/CD11c$^{-}$Ly6C$^{++}$ % of monocytes | n.s. | N/A | N/A |
| Immune system | Flow cytometry | CD11b$^{+}$/CD11c$^{+}$Ly6C$^{(+)}$ % of monocytes | n.s. | N/A | N/A |
| **Immune system** | **Flow cytometry** | **CD11b$^{+}$/CD11c$^{-}$Ly6C$^{(+)}$ % of monocytes** | **0.0361** | other | **other** |
| Immune system | Flow cytometry | CD11b$^{+}$/CD11c$^{+}$Ly6C$^{-}$ % of monocytes | n.s. | N/A | N/A |
| Immune system | Flow cytometry | CD11b$^{+}$/CD11c$^{-}$Ly6C$^{-}$ % of monocytes | n.s. | N/A | N/A |
| **Immune system** | **Flow cytometry** | **CD11b$^{+}$/CD11c$^{+}$Ly6C$^{+}$ monocytes % of all leukocytes** | **0.0017** | ↑ | **20 months** |
| Immune system | Flow cytometry | CD11b$^{+}$/CD11c$^{+}$Ly6C$^{-}$ monocytes % of all leukocytes | n.s. | N/A | N/A |
| **Immune system** | **Flow cytometry** | **CD11b$^{+}$/high Ly6C % of monocytes** | **0.0369** | other | **other** |
| Immune system | Flow cytometry | CD11b$^{+}$/low Ly6C % of monocytes | n.s. | N/A | N/A |
| **Immune system** | **Flow cytometry** | **CD11b$^{+}$/medium Ly6C % of monocytes** | **0.0126** | other | **other** |
| **Immune system** | **Flow cytometry** | **CD11b$^{+}$ monocytes % of all leukocytes** | **0.0173** | ↑ | **20 months** |
| **Immune system** | **Flow cytometry** | **NK1.1$^{+}$/NKp46$^{+}$ NK cells % of all leukocytes** | **0.0016** | ↓ | **20 months** |
| **Immune system** | **Flow cytometry** | **NK1.1$^{+}$/NKp46$^{+}$/CD11b$^{+}$ % of NK cells** | **0.0038** | ↓ | **20 months** |
| Immune system | Flow cytometry | NK1.1$^{+}$/NKp46$^{+}$/CD11c$^{+}$ % of NK cells | n.s. | N/A | N/A |
| **Immune system** | **Flow cytometry** | **NK1.1$^{+}$/NKp46$^{+}$/CD44$^{+}$ of NK cells** | **0.0395** | ↑ | **26 months** |
| Immune system | Flow cytometry | NK1.1$^{+}$/NKp46$^{+}$/Ly6c$^{+}$ % of NK cells | n.s. | N/A | N/A |
| Immune system | Flow cytometry | CD3$^{+}$CD5$^{+}$NK1.1$^{+}$/NKp46$^{+}$ NKT cells % of all leukocytes | n.s. | N/A | N/A |
| Immune system | LPA | Phytohemagglutinin low stimulation | n.s. | N/A | N/A |
| **Immune system** | **LPA** | **Phytohemagglutinin high stimulation** | **0.0179** | ↓ | **14 months** |
| Immune system | LPA | T cells low stimulation | n.s. | N/A | N/A |
| **Immune system** | **LPA** | **T cells high stimulation** | **0.0016** | ↓ | **14 months** |
| **Metabolism** | **BCA** | **Body mass NMR** | **3.56E-11** | ↑ | **8 months** |
| **Metabolism** | **BCA** | **Fat mass NMR** | **7.19E-05** | other | **other** |
| **Metabolism** | **BCA** | **Free fluid NMR** | **2.24E-13** | ↑ | **8 months** |
| **Metabolism** | **BCA** | **Lean mass NMR** | **3.17E-18** | ↑ | **8 months** |
| **Metabolism** | **IpGTT** | **Area under the curve** | **0.0168** | ↓ | **20 months** |
| **Metabolism** | **IpGTT** | **Basal glucose** | **9.22E-05** | ↓ | **20 months** |
| Metabolism | IpGTT | Body weight loss after fasting | n.s. | N/A | N/A |
| **Metabolism** | **Infrared thermovision** | **Average body surface temperature** | **4.66E-08** | ↓ | **5 months** |
| **Metabolism** | **Indirect calorimetry** | **Average heat production** | **0.0489** | ↑ | **26 months** |

| Domain | Test/method | Phenotype | P-value age | Age effect | First detected change at |
|---|---|---|---|---|---|
| **Metabolism** | **Indirect calorimetry** | **Average oxygen consumption** | **0.0340** | ↑ | **26 months** |
| Metabolism | Indirect calorimetry | Average respiratory exchange rate | n.s. | N/A | N/A |
| Metabolism | Indirect calorimetry | Delta respiratory exchange rate | n.s. | N/A | N/A |
| Metabolism | Indirect calorimetry | Food intake | n.s. | N/A | N/A |
| **Metabolism** | **Indirect calorimetry** | **Maximal heat production** | **0.0392** | other | **other** |
| **Metabolism** | **Indirect calorimetry** | **Maximal oxygen consumption** | **0.0296** | other | **other** |
| Metabolism | Indirect calorimetry | Minimal heat production | n.s. | N/A | N/A |
| Metabolism | Indirect calorimetry | Minimal oxygen consumption | n.s. | N/A | N/A |
| Metabolism | Indirect calorimetry | Total distance traveled | n.s. | N/A | N/A |
| Metabolism | Indirect calorimetry | Total number of fine movements | n.s. | N/A | N/A |
| Metabolism | Indirect calorimetry | Total number of rearings | n.s. | N/A | N/A |
| Metabolism | Indirect calorimetry | Water consumption | n.s. | N/A | N/A |
| **Neuropsychiatric functions** | **ASPPI** | **Pre-pulse inhibition global** | **3.67E-12** | ↓ | **14 months** |
| **Neuropsychiatric functions** | **ASPPI** | **Pre-pulse inhibition at 67 dB** | **6.72E-08** | ↓ | **14 months** |
| **Neuropsychiatric functions** | **ASPPI** | **Pre-pulse inhibition at 69 dB** | **2.00E-08** | ↓ | **14 months** |
| **Neuropsychiatric functions** | **ASPPI** | **Pre-pulse inhibition at 73 dB** | **5.31E-11** | ↓ | **20 months** |
| **Neuropsychiatric functions** | **ASPPI** | **Pre-pulse inhibition at 81 dB** | **2.75E-14** | ↓ | **8 months** |
| **Neuropsychiatric functions** | **ASPPI** | **Startle response at 110 dB** | **1.91E-13** | ↓ | **5 months** |
| Neuropsychiatric functions | Grip strength | 2-paws grip strength | n.s. | N/A | N/A |
| **Neuropsychiatric functions** | **Grip strength** | **4-paws grip strength** | **0.0401** | other | **other** |
| **Neuropsychiatric functions** | **Open field** | **Average speed** | **1.79E-08** | ↓ | **20 months** |
| **Neuropsychiatric functions** | **Open field** | **Distance traveled during the first 5 minutes** | **2.10E-09** | ↓ | **20 months** |
| Neuropsychiatric functions | Open field | Distance traveled in the center during the first 5 minutes | n.s. | N/A | N/A |
| **Neuropsychiatric functions** | **Open field** | **Number of rearings during the first 5 minutes** | **9.17E-06** | ↓ | **14 months** |
| Neuropsychiatric functions | Open field | Time spent in the center during the first 5 minutes | n.s. | N/A | N/A |
| **Neuropsychiatric functions** | **Open field** | **Total distance traveled** | **1.82E-07** | ↓ | **20 months** |
| **Neuropsychiatric functions** | **Open field** | **Total distance traveled in the center** | **0.0096** | ↓ | **14 months** |
| **Neuropsychiatric functions** | **Open field** | **Total number of rearings** | **1.35E-07** | ↓ | **8 months** |
| Neuropsychiatric functions | Open field | Total time spent in the center | n.s. | N/A | N/A |
| **Neuropsychiatric functions** | **Rotarod** | **Mean latency to fall** | **4.48E-07** | ↓ | **8 months** |
| Neuropsychiatric functions | SHIRPA | Defecation | n.s. | N/A | N/A |
| **Neuropsychiatric functions** | **SHIRPA** | **Gait** | **4.27E-10** | ↑ | **20 months** |
| **Neuropsychiatric functions** | **SHIRPA** | **Limb grasp** | **2.90E-08** | ↑ | **20 months** |
| **Neuropsychiatric functions** | **SHIRPA** | **Locomotor activity** | **0.0003** | ↓ | **20 months** |
| **Neuropsychiatric functions** | **SHIRPA** | **Startle response** | **7.11E-12** | ↓ | **20 months** |
| Neuropsychiatric functions | SHIRPA | Tail elevation | n.s. | N/A | N/A |
| Neuropsychiatric functions | SHIRPA | Transfer arousal | n.s. | N/A | N/A |

| Domain | Test/method | Phenotype | P-value age | Age effect | First detected change at |
|---|---|---|---|---|---|
| **Neuropsychiatric functions** | **SHIRPA** | **Urination** | **0.0201** | ↑ | **26 months** |
| **Sensory systems** | **Hot plate** | **Time to first response** | **0.0035** | ↑ | **20 months** |
| **Sensory systems** | **Hot plate** | **Time to second response** | **5.01E-08** | ↑ | **14 months** |
| **Sensory systems** | **OCT** | **Average retina thickness** | **0.0008** | other | **other** |

ASPPI = Acoustic startle and pre-pulse inhibition; BCA = Body composition analysis; CD = Cluster of differentiation; ELISA = Enzyme-linked immunosorbent assay; HDL = High density lipoprotein; IHC = Immunohistochemistry; Ig = Immunoglobulin; IpGTT = Intraperitoneal glucose tolerance test; LPA = Lymphocyte proliferation assay; N/A = Not applicable; NK = Natural killer cells; NMR = Nuclear magnetic resonance; n.s. = not significant; OCT = Optical coherence tomography; TEWL = Transepidermal water loss

### Extended Data Table 2. Antibodies used in flow cytometry based analyses

| Panel | Fluorochrome | Cell surface marker | Clone | Company | Dilution |
|---|---|---|---|---|---|
| Panel 1 | FITC | CD11c | HL3 | BD Pharmingen, #557400 | 1:100 |
| | PE | NK1.1 | PK136 | BD Pharmingen, #553165 | 1:200 |
| | PE | NKp46 | 29A1.4 | eBioscience, #12-3351-82 | 1:200 |
| | PE-CF594 | CD3e | 145-2C11 | BD Horizon, #562332 | 1:100 |
| | PerCP Cy5.5 | Ly6C | HK1.4 | eBioscience, #45-5932-82 | 1:400 |
| | PECy7 | CD19 | 1D3 | BD Pharmingen, #552854 | 1:1000 |
| | APC | CD5 | 53-7.3 | BD Pharmingen, #550035 | 1:2000 |
| | Alexa Fluor 700 | CD45 | 30-F11 | BioLegend, #103128 | 1:1000 |
| | APC-A750 | B220 | RA3-6B2 | Life Technologies, #RM2627 | 1:100 |
| | PacBlue | CD11b | M1/70.15 | Life Technologies, #RM2828 | 1:800 |
| | PO | Gr1 | RB6-8C5 | Life Technologies, #RM3030 | 1:1000 |
| Panel 2 | PE-CF594 | Ly6C | AL-21 | BD Horizon, #562728 | 1:200 |
| | PerCP Cy5.5 | CD4 | RM4-5 | TONBO Biosciences, #65-0042-U025 | 1:1000 |
| | PECy7 | CD62L | MEL-14 | eBioscience, #25-0621-82 | 1:2000 |
| | APC | CD25 | PC61 | BD Pharmingen, #557192 | 1:100 |
| | Alexa Fluor 700 | CD45 | 30-F11 | BioLegend, #103128 | 1:1000 |
| | APC-A750 | CD8a | 5H10 | Life Technologies, #MCD0827 | 1:400 |
| | eF450 | CD5 | 53-7.3 | eBioscience, #48-0051-82 | 1:1000 |
| | bv570 | CD44 | IM7 | BioLegend, #103037 | 1:100 |

APC = allophycocyanin; Cy7 = cyanine-7; FITC = fluorescein-5-isothiocyanate; PE = phycoerythrin; PerCP = peridin chlorophyll; PO = pacific orange

### Extended Data Table 3. Molecular assays to study putative drivers of aging

| Aging hallmark | Subcategory | Target | Method |
|---|---|---|---|
| **Altered intercellular communication** | **Lipid hormone** | Cox1 | WB |
| | **Inflammation** | *Ccl2* | qPCR |
| | | *Ifng* | qPCR |
| | | *Il1b* | qPCR |
| | | *Il4* | qPCR |
| | | *Il6* | qPCR |
| | | *Il10* | qPCR |
| | | *Il13* | qPCR |
| | | *Tnf* | qPCR |
| **Cellular senescence** | **Senescence markers** | *Cdkn2a/p16Ink4a* | qPCR |
| | | *Cdkn2a/p19Arf* | qPCR |
| | | *Cdkn1a/p21* | qPCR |
| | **Tumor suppressor** | *Trp53* | qPCR |
| **Deregulated nutrient sensing** | **IGF1-signaling** | Igf1 | WB |
| | **mTOR-signaling** | mTOR | WB |
| | | p-4Ebp1 (T37/46)/4Ebp1 | WB |
| | | total 4Ebp1 | WB |
| | | p-Rps6 (S240/244)/Rps6 | WB |
| | | total Rps6 | WB |
| | | p-Akt (S473)/Akt | WB |
| | | total Akt | WB |
| **Genomic instability** | **DNA damage** | 8-oxo-guanosine | ELISA |
| | | p-H2ax (S139)/H2ax | WB |
| | | total H2ax | WB |
| | | Tp53bp1 | WB |
| | **Transposons** | *LINE* | qPCR |
| | | *L1 5'UTR* | qPCR |
| | | *L1 3'UTR* | qPCR |
| | | *MusD* | qPCR |
| | | *B1* | qPCR |
| | | *B2* | qPCR |
| **Loss of proteostasis** | **Autophagy** | Atg3 | WB |
| | | Atg5 | WB |
| | | Lc3a/b II/I | WB |
| | | total Lc3a/b | WB |
| | **Chaperones** | Hsp60 | WB |
| | | Hsp70 | WB |
| | | Hsp90 | WB |
| | **Proteasome activity** | 20S activity | activity assay |
| | **Ubiquitin** | Mono-ubiquitin | WB |
| | | Poly-ubiquitin | WB |
| **Mitochondrial dysfunction** | **Lipid peroxidation** | TBA reactive species | chemical reaction |
| | **Mitochondrial integrity** | Citrate synthase | WB |
| | | Cox IV | WB |
| | | Sod2 | WB |
| | **Oxidative stress** | ROS production | chemical reaction |
| | | Nitrotyrosine | WB |
| **Reduced cell proliferation** | **Cell cycle regulators** | *Ccna1* | qPCR |
| | | *Ccna2* | qPCR |
| | | *Ccnb1* | qPCR |
| | | *Ccnb2* | qPCR |
| | | *Ccnb3* | qPCR |
| | | *Ccnc* | qPCR |
| | | *Ccnd1* | qPCR |
| | | *Ccnd2* | qPCR |
| | | *Ccnd3* | qPCR |
| | | *Ccne1* | qPCR |
| | | *Ccne2* | qPCR |
| | **Cell proliferation marker** | *Mki67* | qPCR |

Akt = Protein kinase B; Cox1 = Cyclooxygenase 1; Cox IV = Cytochrom c oxidase IV; ELISA = Enzyme-linked immunosorbent assay; Hsp = Heat shock protein; Igf1 = Insulin-like growth factor 1; Lc3 = Microtubule associated protein 1A/1B light chain 3; mTOR = Mechanistic target of rapamycin; qPCR = Quantitative polymerase chain reaction; ROS = Reactive oxygen species; Rps6 = Ribosomal protein S6; Sod2 = Superoxide dismutase 2; TBA = Thiobarbituric acid; Tp53bp1 = Tumor suppressor p53-binding protein 1; WB = Western blot

**Extended Data Table 4. Primer sequences used for real-time quantitative PCR analyses**

| Aging hallmark | Gene/transposon | Primer forward | Primer reverse |
|---|---|---|---|
| **Altered intercellular communication** | *Ccl2* | AAGAGATCAGGGAGTTTGCT | CTGCCTCCATCAACCACTTT |
| | *Ifng* | CTTTGGACCCTCTGACTTGAG | TCAATGACTGTGCCGTGG |
| | *Ilb1* | GAAGAAGAGCCCATCCTCTG | TCATCTCGGAGCCTGTAGTG |
| | *Il4* | GCATTTTGAACGAGGTCACAG | TGGAAGCCCTACAGACGAG |
| | *Iil6* | AGTCCGGAGAGGAGACTTCA | ATTTCCACGATTTCCCAGAG |
| | *Il10* | AGCCGGGAAGACAATAACTG | GGAGTCGGTTAGCAGTATGTTG |
| | *Il13* | ACCAAAATCGAAGTAGCCCAC | GCAAAGTCTGATGTGAGAAAGG |
| | *Tnf* | CTTCTGTCTACTGAACTTCGGG | CAGGCTTGTCACTCGAATTTTG |
| **Cellular proliferation** | *Ccna1* | GGGTGTTGACTGAAAATGAGC | CACGTTTGGCTGGTTCATTG |
| | *Ccna2* | GTCCTTGCTTTTGACTTGGC | ACGGGTCAGCATCTATCAAAC |
| | *Ccnb1* | CTGACCCAAACCTCTGTAGTG | CCTGTATTAGCCAGTCAATGAGG |
| | *Ccnb2* | CCTCAGAACACCAAAGTACCAG | CCTTCATGGAGACATCCTCAG |
| | *Ccnb3* | TCCAGTGCTATCATGCCAAG | CTGTCACTGTCATCCTGTATGG |
| | *Ccnc* | GCATTTGTATCAGGGCAAGC | GAAACTTTAGGTCCTTTTGGCG |
| | *Ccnd1* | GCCCTCCGTATCTTACTTCAAG | GCGGTCCAGGTAGTTCATG |
| | *Ccnd2* | GTGTTCCTATTTCAAGTGCGTG | AGCCAAGAAACGGTCCAG |
| | *Ccnd3* | GCGTGCAAAAGGAGATCAAG | GATCCAGGTAGTTCATAGCCAG |
| | *Ccne1* | GCGAGGATGAGAGCAGTTC | AAGTCCTGTGCCAAGTAGAAC |
| | *Ccne2* | GACGTTCATCCAGATAGCTCAG | TCCCATTCCAAACCTGAAGC |
| | *Mki67* | TGCCCGACCCTACAAAATG | GAGCCTGTATCACTCATCTGC |
| **Cellular senescence** | *Cdkn2a/p16Ink4a* | CCCAACGCCCCGAACT | GCAGAAGAGCTGCTACGTGAA |
| | *Cdkn2a/p19Arf* | CTCTGGCTTTCGTGAACATG | TCGAATCTGCACCGTAGTTG |
| | *Cdkn1a/p21* | CAGATCCACAGCGATATCCAG | AGAGACAACGGCACACTTTG |
| | *Trp53* | ATGTTCCGGGAGCTGAATG | CCCCACTTTCTTGACCATTG |
| **Genomic instability** | *LINE* | TGAGTGGAACACAACTTCTGC | CAGGCAAGCTCTCTTCTTGC |
| | *L1 5'UTR* | CTGCCTTGCAAGAAGAGAGC | AGTGCTGCGTTCTGATGATG |
| | *L1 3'UTR* | CCAGCAAACACAGAAGTGGATGCTCA | TTTGCAAGTCCAATGGGCCTCTCT |
| | *MusD* | ATAGAGGCCGCTTCTTTGC | TGAGACTCCACCAAATGTCC |
| | *B1* | CATGGTGGCGCACGCCTTTAATCC | CCAGGCTGGCCTCGAACTCAGAAA |
| | *B2* | GGGCTGGAGAGATGGCTCAGTGGT | GCCACCATGTGGTTGCTGGGAATTG |
| | *Actb* | CCCTGAAGTACCCCATTGAAC | CCATGTCGTCCCAGTTGGTAA |

### Extended Data Table 5. Antibodies used in the context of Western Blot based analyses

| Aging hallmark | Target | Antibody used | Dilution |
|---|---|---|---|
| **Altered intercellular communication** | Cox1 | Cell Signaling Technologies, #4841 | 1:2000 |
| **Deregulated nutrient sensing** | Igf1 | Abcam, ab9572 | 1:1000 |
| | mTOR | Cell Signaling Technologies, #2983 | 1:2000 |
| | p-4Ebp1 (T37/46) | Cell Signaling Technologies, #2855 | 1:2000 |
| | total 4Ebp1 | Cell Signaling Technologies, #9644 | 1:30000 |
| | p-Rps6 (S240/244) | Cell Signaling Technologies, #2215 | 1:2000 |
| | total Rps6 | Cell Signaling Technologies, #2217 | 1:10000 |
| | p-Akt (S473) | Cell Signaling Technologies, #9271 | 1:2000 |
| | total Akt | Cell Signaling Technologies, #9272 | 1:5000 |
| **Genomic instability** | p-H2ax (S139) | Cell Signaling Technologies, #2577 | 1:2000 |
| | H2ax | Cell Signaling Technologies, #2595 | 1:2000 |
| | Tp53bp1 | Abnova, PAB12506 | 1:2000 |
| **Loss of proteostasis** | Atg3 | Cell Signaling Technologies, #3415 | 1:2000 |
| | Atg5 | Cell Signaling Technologies, #12994 | 1:2000 |
| | Lc3a/b | Cell Signaling Technologies, #12741 | 1:3000 |
| | Hsp60 | Cell Signaling Technologies, #4870 | 1:10000 |
| | Hsp70 | Cell Signaling Technologies, #4872 | 1:10000 |
| | Hsp90 | Cell Signaling Technologies, #4874 | 1:10000 |
| | Mono-/Poly-ubiquitin | Thermo Fisher Scientific, PA5-76144 | 1:2000 |
| **Mitochondrial dysfunction** | Citrate synthase | Cell Signaling Technologies, #14309 | 1:2000 |
| | Cox IV | Cell Signaling Technologies, #4850 | 1:2000 |
| | Sod2 | Cell Signaling Technologies, #13194 | 1:2000 |
| | Nitrotyrosine | Enzo Life Science, BML-SA297 | 1:2000 |
| | Actin | MP Biomedicals, SKU 0869100 | 1:30000 |

Akt = Protein kinase B; Cox1 = Cyclooxygenase 1; Cox IV = Cytochrom c oxidase IV; Hsp = Heat shock protein; Igf-1 = Insulin-like growth factor 1; Lc3 = Microtubule associated protein 1A/1B light chain 3; mTOR = Mechanistic target of rapamycin; Rps6 = Ribosomal protein S6; Sod2 = Superoxide dismutase 2; Tp53bp1 = Tumor suppressor p53-binding protein 1

### Extended Data Table 6. Age-sensitive phenotypes countered or accentuated in *Ghrhr*[lit/lit] mice

The columns “Age”, “Genotype” and “Interaction” summarize results of two-way ANOVAs/aligned rank transform (summarizing main effects of age, genotype and genotype x age interactions, respectively). Findings summarized in the “Intervention effect” column are based on statistical comparison of Cohen’s d intervention effect sizes in young vs. old mice. Comparisons, for which a calculation of Cohen’s d was not possible, are classified as “not applicable” (N/A).

| Domain | Test/method | Phenotype | Category | Age | Genotype | Interaction | Intervention effect |
|---|---|---|---|---|---|---|---|
| Anatomy and physiology | Bone densitometry | Bone mineral density | Promoting aging | ↓ | ↓ | yes | in young > old |
| Anatomy and physiology | Organ weight | Heart weight | Opposing aging | ↑ | ↓ | no | equal in young and old |
| Anatomy and physiology | Organ weight | Kidney weight | Opposing aging | ↑ | ↓ | no | equal in young and old |
| Anatomy and physiology | Organ weight | Liver weight | Opposing aging | ↑ | ↓ | no | in young < old |
| Cardiovascular health | Echocardiography | Corrected mass of the left ventricle | Opposing aging | ↑ | ↓ | yes | equal in young and old |
| Cardiovascular health | Echocardiography | Left ventricular end-diastolic internal diameter | Opposing aging | ↑ | ↓ | yes | equal in young and old |
| Cardiovascular health | Echocardiography | Left ventricular end-systolic internal diameter | Opposing aging | ↑ | ↓ | yes | in young < old |
| Cardiovascular health | Echocardiography | Stroke volume | Opposing aging | ↑ | ↓ | no | equal in young and old |
| Cardiovascular health | Electrocardiography | Duration of the PR interval | Opposing aging | ↑ | ↓ | no | equal in young and old |
| Clinical chemistry | Clinical chemistry | Alkaline phosphatase | Opposing aging | ↑ | ↓ | yes | in young < old |
| Clinical chemistry | Clinical chemistry | Alpha amylase | Opposing aging | ↑ | ↓ | no | equal in young and old |
| Clinical chemistry | Clinical chemistry | Phosphate | Opposing aging | ↑ | ↓ | no | equal in young and old |
| Clinical chemistry | Clinical chemistry | Total protein | Opposing aging | ↑ | ↓ | no | in young > old |
| Clinical chemistry | Clinical chemistry | Unsaturated iron binding capacity | Opposing aging | ↑ | ↓ | no | equal in young and old |
| Hematology | Hematology | Hematocrit | Opposing aging | ↓ | ↑ | yes | in young < old |
| Hematology | Hematology | Hemoglobin concentration | Opposing aging | ↓ | ↑ | yes | in young < old |
| Hematology | Hematology | Mean corpuscular volume | Promoting aging | ↑ | ↑ | no | equal in young and old |
| Hematology | Hematology | Mean platelet volume | Opposing aging | ↑ | ↓ | no | equal in young and old |
| Hematology | Hematology | Platelet distribution width | Opposing aging | ↑ | ↓ | no | equal in young and old |
| Hematology | Hematology | Platelet count | Opposing aging | ↑ | ↓ | no | in young > old |
| Immune system | ELISA | Interleukin-6 | Opposing aging | ↑ | ↓ | no | equal in young and old |
| Metabolism | BCA | Body mass NMR | Opposing aging | ↑ | ↓ | no | in young > old |
| Metabolism | Infrared thermovision | Average body surface temperature | Promoting aging | ↓ | ↓ | no | equal in young and old |
| Metabolism | Indirect calorimetry | Average heat production | Opposing aging | ↑ | ↓ | no | equal in young and old |
| Metabolism | Indirect calorimetry | Average respiratory exchange rate | Promoting aging | ↓ | ↓ | no | equal in young and old |
| Metabolism | Indirect calorimetry | Average oxygen consumption | Opposing aging | ↑ | ↓ | no | equal in young and old |
| Metabolism | Indirect calorimetry | Maximal heat production | Opposing aging | ↑ | ↓ | no | equal in young and old |
| Metabolism | Indirect calorimetry | Maximal oxygen consumption | Opposing aging | ↑ | ↓ | no | equal in young and old |
| Metabolism | Indirect calorimetry | Maximal respiratory exchange rate | Promoting aging | ↓ | ↓ | no | equal in young and old |
| Metabolism | Indirect calorimetry | Minimal heat production | Opposing aging | ↑ | ↓ | no | in young > old |
| Metabolism | Indirect calorimetry | Minimal oxygen consumption | Opposing aging | ↑ | ↓ | no | in young > old |
| Metabolism | Indirect calorimetry | Minimal respiratory exchange rate | Promoting aging | ↓ | ↓ | no | equal in young and old |

| Domain | Test/method | Phenotype | Category | Age | Genotype | Interaction | Intervention effect |
|---|---|---|---|---|---|---|---|
| Metabolism | Indirect calorimetry | Total number of fine movements | Opposing aging | ↑ | ↓ | yes | equal in young and old |
| Metabolism | Indirect calorimetry | Total distance traveled | Opposing aging | ↑ | ↓ | no | equal in young and old |
| Metabolism | Indirect calorimetry | Total number of rearings | Opposing aging | ↑ | ↓ | no | equal in young and old |
| Metabolism | IpGTT | Area under the curve | Opposing aging | ↓ | ↑ | no | equal in young and old |
| Neuropsychiatric functions | ABR | Click ABR | N/A | ↑ | N/A | no | N/A |
| Neuropsychiatric functions | ABR | Threshold at 6kHz | N/A | ↑ | N/A | no | N/A |
| Neuropsychiatric functions | ABR | Threshold at 12kHz | Promoting aging | ↑ | ↑ | no | equal in young and old |
| Neuropsychiatric functions | ABR | Threshold at 18kHz | N/A | ↑ | N/A | yes | N/A |
| Neuropsychiatric functions | ABR | Threshold at 24kHz | N/A | ↑ | N/A | yes | N/A |
| Neuropsychiatric functions | ABR | Threshold at 30kHz | N/A | ↑ | N/A | yes | N/A |
| Neuropsychiatric functions | Grip strength | 2-paws grip strength | Promoting aging | ↓ | ↓ | no | in young > old |
| Neuropsychiatric functions | Open field | Distance traveled during the first 5 minutes | Promoting aging | ↓ | ↓ | no | equal in young and old |
| Neuropsychiatric functions | Open field | Total distance traveled | Promoting aging | ↓ | ↓ | no | equal in young and old |
| Neuropsychiatric functions | Open field | Number of rearings during the first 5 minutes | Promoting aging | ↓ | ↓ | no | equal in young and old |
| Neuropsychiatric functions | Open field | Total number of rearings | Promoting aging | ↓ | ↓ | no | equal in young and old |
| Neuropsychiatric functions | Open field | Average speed | Promoting aging | ↓ | ↓ | no | equal in young and old |
| Neuropsychiatric functions | Rotarod | Mean latency to fall | Promoting aging | ↓ | ↓ | no | equal in young and old |
| Neuropsychiatric functions | SHIRPA | Gait | Opposing aging | ↑ | ↓ | yes | equal in young and old |
| Neuropsychiatric functions | SHIRPA | Grade_of_alopecia | Opposing aging | ↑ | ↓ | yes | equal in young and old |
| Neuropsychiatric functions | SHIRPA | Limb grasp | Opposing aging | ↑ | ↓ | yes | equal in young and old |
| Neuropsychiatric functions | SHIRPA | Locomotor activity | Promoting aging | ↓ | ↓ | no | equal in young and old |
| Neuropsychiatric functions | SHIRPA | Startle response | Promoting aging | ↓ | ↓ | no | equal in young and old |
| Sensory systems | Hot plate | Time to first response | Opposing aging | ↑ | ↓ | no | equal in young and old |
| Sensory systems | Hot plate | Time to second response | Opposing aging | ↑ | ↓ | no | equal in young and old |
| Sensory systems | LIB | Anterior chamber depth | Opposing aging | ↑ | ↓ | yes | equal in young and old |
| Sensory systems | LIB | Axial length | Opposing aging | ↑ | ↓ | no | equal in young and old |
| Sensory systems | LIB | Eye body length | Opposing aging | ↑ | ↓ | no | equal in young and old |
| Sensory systems | LIB | Lens thickness | Promoting aging | ↑ | ↑ | yes | in young < old |
| Sensory systems | Virtual drum | Spatial frequency threshold | Promoting aging | ↓ | ↓ | no | equal in young and old |

ABR = Auditory brain stem response; BCA = Body composition analysis; ELISA = Enzyme-linked immunosorbent assay; IpGTT = Intraperitoneal glucose tolerance test; LIB = Laser interference biometry; N/A = Not applicable; NMR = Nuclear magnetic resonance

**Extended Data Table 7. Age and genotype effect in *GHRHR*-related (endo)phenotypic measures in humans**
Boldface indicates significance. *) Age was mean-centered before inclusion in the regression models.

| Determinant | Change in outcome (SD) [estimate (95% CI)] | | | | | |
|---|---|---|---|---|---|---|
| | *Platelet* | *p-value* | *Cholesterol level* | *p-value* | *LDL level* | *p-value* |
| *GHRHR eQTL* | ***-0.067 (-0.123, -0.011)*** | ***0.019*** | ***-0.059 (-0.115, -0.003)*** | ***0.039*** | ***-0.076 (-0.133, -0.019)*** | ***0.009*** |
| *Age** | ***-0.007 (-0.013, -0.001)*** | ***0.026*** | ***0.019 (0.013, 0.025)*** | ***$1.5*10^{-09}$*** | ***0.016 (0.010, 0.022)*** | ***$5.7*10^{-07}$*** |
| *GHRHR eQTL x age* | 0.000 (-0.004, 0.004) | 0.891 | -0.003 (-0.007, 0.001) | 0.143 | -0.003 (-0.008, 0.001) | 0.096 |

CI = confidence interval; eQTL = expression quantitative trait loci; *GHRHR* = growth hormone releasing hormone receptor; LDL = low-density lipoproteins; SD = standard deviation

**Extended Data Table 8. Characteristics of the human study population**

| | Overall (n= 3034) |
|---|---|
| Age [year], mean ± SD (range) | 56.2 ± 14.3 (30 - 95) |
| Sex, n (%) | |
| Women | 1700 (56) |
| Men | 1334 (44) |
| *MTOR* eQTL genotype, n (%) | |
| GG | 1506 (49.6) |
| CG | 1203 (39.7) |
| CC | 281 (9.3) |
| *GHRHR* eQTL genotype, n (%) | |
| GG | 254 (8.4) |
| AG | 1339 (44.1) |
| AA | 1388 (45.7) |

eQTL = expression quantitative trait locus; *GHRHR* = growth hormone releasing hormone receptor; *MTOR* = mammalian target of rapamycin; SD = standard deviation.
*MTOR* eQTL genotype: 44 missing; *GHRHR* eQTL genotype: 53 missing

**Extended Data Table 9. Age-sensitive phenotypes countered or accentuated in *mTOR*$^{KI/KI}$ mice**

The columns "Age", "Genotype" and "Interaction" summarize results of two-way ANOVAs/aligned rank transform (summarizing main effects of age, genotype and genotype x age interactions, respectively). Findings summarized in the "Intervention effect" column are based on statistical comparison of Cohen's d intervention effect sizes in young vs. old mice. Comparisons, for which a calculation of Cohen's d was not possible, are classified as "not applicable" (N/A).

| Domain | Test/method | Phenotype | Category | Age | Genotype | Interaction | Intervention effect |
|---|---|---|---|---|---|---|---|
| Anatomy and physiology | Bone densitometry | Bone mineral density | Promoting aging | ↓ | ↓ | yes | in young > old |
| Anatomy and physiology | Organ weight | Heart weight | Opposing aging | ↑ | ↓ | no | equal in young and old |
| Anatomy and physiology | Organ weight | Kidney weight | Opposing aging | ↑ | ↓ | no | equal in young and old |
| Anatomy and physiology | Organ weight | Liver weight | Opposing aging | ↑ | ↓ | no | equal in young and old |
| Anatomy and physiology | Organ weight | Lung weight | Opposing aging | ↑ | ↓ | no | in young > old |
| Anatomy and physiology | Organ weight | Muscle weight | Promoting aging | ↓ | ↓ | no | equal in young and old |
| Anatomy and physiology | Organ weight | Pancreas weight | Opposing aging | ↑ | ↓ | no | equal in young and old |
| Anatomy and physiology | Organ weight | Testis weight | Promoting aging | ↓ | ↓ | no | in young > old |
| Cardiovascular health | Echocardiography | Corrected mass of the left ventricle | Opposing aging | ↑ | ↓ | yes | equal in young and old |
| Cardiovascular health | Echocardiography | Ejection fraction | Opposing aging | ↓ | ↑ | no | equal in young and old |
| Cardiovascular health | Echocardiography | Left ventricular end-diastolic internal diameter | Opposing aging | ↑ | ↓ | yes | equal in young and old |
| Cardiovascular health | Echocardiography | Left ventricular end-systolic internal diameter | Opposing aging | ↑ | ↓ | no | equal in young and old |
| Cardiovascular health | Echocardiography | Stroke volume | Opposing aging | ↑ | ↓ | yes | equal in young and old |
| Cardiovascular health | Electrocardiography | Duration of the RR interval | Opposing aging | ↑ | ↓ | no | equal in young and old |
| Cardiovascular health | Electrocardiography | Heart rate | Opposing aging | ↓ | ↑ | no | equal in young and old |
| Clinical chemistry | Clinical chemistry | Alpha amylase | Promoting aging | ↑ | ↑ | yes | in young < old |
| Clinical chemistry | Clinical chemistry | Ca | Opposing aging | ↑ | ↓ | no | equal in young and old |
| Clinical chemistry | Clinical chemistry | Glycerol after fasting | Promoting aging | ↑ | ↑ | no | equal in young and old |
| Clinical chemistry | Clinical chemistry | K | Opposing aging | ↑ | ↓ | no | equal in young and old |
| Clinical chemistry | Clinical chemistry | Total protein | Opposing aging | ↑ | ↓ | no | equal in young and old |
| Clinical chemistry | Clinical chemistry | Triglycerides | Opposing aging | ↓ | ↑ | yes | in young < old |
| Clinical chemistry | Clinical chemistry | Unsaturated iron binding capacity | Opposing aging | ↑ | ↓ | no | equal in young and old |
| Hematology | Hematology | Hematocrit | Opposing aging | ↓ | ↑ | yes | in young < old |
| Hematology | Hematology | Hemoglobin concentration | Opposing aging | ↓ | ↑ | yes | in young < old |
| Hematology | Hematology | Mean corpuscular hemoglobin content | Promoting aging | ↓ | ↓ | no | equal in young and old |
| Hematology | Hematology | Mean corpuscular volume | Promoting aging | ↓ | ↓ | no | equal in young and old |
| Hematology | Hematology | Mean platelet volume | Opposing aging | ↑ | ↓ | no | equal in young and old |
| Hematology | Hematology | Number of eosinophils | Promoting aging | ↑ | ↑ | no | equal in young and old |
| Hematology | Hematology | Number of neutrophils | Opposing aging | ↑ | ↓ | no | equal in young and old |
| Immune system | ELISA | IgE | Opposing aging | ↑ | ↓ | no | equal in young and old |
| Immune system | Flow cytometry | CD19$^+$B220$^+$/CD11b$^+$ % of B cells | Opposing aging | ↑ | ↓ | yes | equal in young and old |
| Immune system | Flow cytometry | CD3$^+$CD5$^+$ T cells % of all leukocytes | Opposing aging | ↓ | ↑ | no | equal in young and old |

| Domain | Test/method | Phenotype | Category | Age | Genotype | Interaction | Intervention effect |
|---|---|---|---|---|---|---|---|
| Immune system | Flow cytometry | $CD5^{+}/CD4^{+}CD44^{++}$ % of $CD4^{+}$ T cells | Opposing aging | ↑ | ↓ | yes | equal in young and old |
| Immune system | Flow cytometry | $CD5^{+}/CD4^{+}CD44^{+}CD62L^{-}Ly6C^{-}$ % of $CD4^{+}$ T cells | Opposing aging | ↑ | ↓ | yes | in young < old |
| Immune system | Flow cytometry | $CD5^{+}/CD4^{+}CD44^{+}CD62L^{-}Ly6C^{+}$ % of $CD4^{+}$ T cells | Opposing aging | ↓ | ↑ | yes | in young < old |
| Immune system | Flow cytometry | $CD5^{+}/CD4^{+}CD62L^{+}$ % of $CD4^{+}$ T cells | Opposing aging | ↓ | ↑ | yes | in young < old |
| Immune system | Flow cytometry | $CD5^{+}/CD8^{+}CD44^{+}$ % of $CD8^{+}$ T cells | Opposing aging | ↑ | ↓ | no | equal in young and old |
| Immune system | Flow cytometry | $CD5^{+}/CD8^{+}CD44^{++}CD62L^{+}Ly6C^{-}$ % of $CD8^{+}$ T cells | Opposing aging | ↑ | ↓ | yes | in young < old |
| Immune system | Flow cytometry | $CD5^{+}/CD8^{+}CD62L^{+}$ % of $CD8^{+}$ T cells | Opposing aging | ↓ | ↑ | no | equal in young and old |
| Immune system | Flow cytometry | $CD5^{+}/CD8^{+}Ly6C^{+}$ % of $CD8^{+}$ T cells | Promoting aging | ↑ | ↑ | yes | in young < old |
| Immune system | Flow cytometry | $CD11b^{+}/CD11c^{+}Ly6C^{+}$ % of monocytes | Opposing aging | ↑ | ↓ | no | equal in young and old |
| Immune system | Flow cytometry | $CD11b^{+}/CD11c^{+}Ly6C^{-}$ % of monocytes | Opposing aging | ↑ | ↓ | yes | equal in young and old |
| Immune system | Flow cytometry | $CD11b^{+}$ monocytes % of all leukocytes | Opposing aging | ↑ | ↓ | yes | equal in young and old |
| Immune system | Flow cytometry | $NK1.1^{+}/NKp46^{+}$ NK cells % of all leukocytes | Promoting aging | ↓ | ↓ | no | equal in young and old |
| Immune system | Flow cytometry | $NK1.1^{+}/NKp46^{+}/CD11c^{+}$ % of NK cells | Opposing aging | ↑ | ↓ | no | equal in young and old |
| Metabolism | BCA | Body mass NMR | Opposing aging | ↑ | ↓ | yes | equal in young and old |
| Metabolism | BCA | Fat mass NMR | Opposing aging | ↑ | ↓ | yes | equal in young and old |
| Metabolism | BCA | Free fluid NMR | Opposing aging | ↑ | ↓ | yes | equal in young and old |
| Metabolism | BCA | Lean mass NMR | Opposing aging | ↑ | ↓ | no | equal in young and old |
| Metabolism | Indirect calorimetry | Average heat production | Opposing aging | ↑ | ↓ | no | equal in young and old |
| Metabolism | Indirect calorimetry | Average oxygen consumption | Opposing aging | ↑ | ↓ | no | equal in young and old |
| Metabolism | Indirect calorimetry | Delta respiratory exchange rate | Opposing aging | ↓ | ↑ | no | equal in young and old |
| Metabolism | Indirect calorimetry | Food intake | Opposing aging | ↓ | ↑ | no | in young < old |
| Metabolism | Indirect calorimetry | Maximal heat production | Opposing aging | ↑ | ↓ | no | equal in young and old |
| Metabolism | Indirect calorimetry | Maximal oxygen consumption | Opposing aging | ↑ | ↓ | yes | equal in young and old |
| Metabolism | Indirect calorimetry | Minimal heat production | Opposing aging | ↑ | ↓ | no | equal in young and old |
| Metabolism | Indirect calorimetry | Minimal oxygen consumption | Opposing aging | ↑ | ↓ | no | equal in young and old |
| Metabolism | IpGTT | Area under the curve | Promoting aging | ↑ | ↑ | no | equal in young and old |
| Metabolism | IpGTT | Body weight loss after fasting | Opposing aging | ↑ | ↓ | no | equal in young and old |
| Neuropsychiatric functions | ABR | Click ABR | Opposing aging | ↑ | ↓ | no | in young < old |
| Neuropsychiatric functions | ABR | Threshold at 6kHz | Opposing aging | ↑ | ↓ | yes | in young < old |
| Neuropsychiatric functions | ABR | Threshold at 12kHz | Opposing aging | ↑ | ↓ | yes | in young < old |
| Neuropsychiatric functions | ABR | Threshold at 24kHz | N/A | ↑ | N/A | yes | N/A |
| Neuropsychiatric functions | ABR | Threshold at 30kHz | N/A | ↑ | N/A | yes | N/A |
| Neuropsychiatric functions | ASPPI | Pre-pulse inhibition at 73 dB | Opposing aging | ↓ | ↑ | yes | equal in young and old |
| Neuropsychiatric functions | ASPPI | Startle response at 110 dB | Promoting aging | ↓ | ↓ | yes | in young < old |
| Neuropsychiatric functions | Grip strength | 2-paws grip strength | Opposing aging | ↑ | ↓ | no | equal in young and old |
| Neuropsychiatric functions | Open field | Number of rearings during the first 5 minutes | Promoting aging | ↓ | ↓ | no | equal in young and old |
| Neuropsychiatric functions | Open field | Total number of rearings | Promoting aging | ↓ | ↓ | no | equal in young and old |

| Domain | Test/method | Phenotype | Category | Age | Genotype | Interaction | Intervention effect |
|---|---|---|---|---|---|---|---|
| Neuropsychiatric functions | Rotarod | Mean latency to fall | Opposing aging | ↓ | ↑ | no | equal in young and old |
| Neuropsychiatric functions | SHIRPA | Gait | Opposing aging | ↑ | ↓ | yes | equal in young and old |
| Neuropsychiatric functions | SHIRPA | Startle response | Opposing aging | ↓ | ↑ | no | equal in young and old |
| Neuropsychiatric functions | SHIRPA | Tremor | N/A | ↑ | ↑ | yes | N/A |
| Neuropsychiatric functions | SHIRPA | Vocalisation | N/A | ↓ | N/A | yes | N/A |
| Sensory systems | OCT | Average retina thickness | Promoting aging | ↓ | ↓ | no | equal in young and old |

ABR = Auditory brain stem response; ASPPI = Acoustic startle and pre-pulse inhibition; BCA = Body composition analysis; CD = Cluster of differentiation; ELISA = Enzyme-linked immunosorbent assay; IpGTT = Intraperitoneal glucose tolerance test; N/A = not applicable; NK = Natural killer cells; NMR = Nuclear magnetic resonance; OCT = Optical coherence tomography

**Extended Data Table 10. Age and genotype effect in *MTOR*-related (endo)phenotypic measures in humans**
Boldface indicates significance. ***)** Age was mean-centered before inclusion in the regression models.

| Determinant | Change in outcome (SD) [estimate (95% CI)] | | | | | | | | | |
|---|---|---|---|---|---|---|---|---|---|---|
| | *Body fat* | *p-value* | *% Body fat* | *p-value* | *Body weight* | *p-value* | *Creatine level* | *p-value* | *MET hours* | *p-value* |
| *MTOR eQTL* | ***0.066*** | ***0.018*** | ***0.048*** | ***0.042*** | ***0.055*** | ***0.024*** | ***0.079*** | ***0.013*** | ***0.059*** | ***0.049*** |
| | ***(0.011, 0.12)*** | | ***(0.002, 0.095)*** | | ***(0.007, 0.103)*** | | ***(0.016, 0.141)*** | | ***(0, 0.118)*** | |
| *Age** | ***0.014*** | ***$2.2*10^{-15}$*** | ***0.022*** | ***$< 2.0*10^{-16}$*** | ***-0.003*** | | ***0.011*** | ***$7.3*10^{-08}$*** | ***-0.013*** | ***$1.6*10^{-12}$*** |
| | ***(0.01, 0.017)*** | | ***(0.02, 0.025)*** | | ***(-0.006, 0)*** | ***0.033*** | ***(0.007, 0.015)*** | | ***(-0.016, -0.009)*** | |
| *MTOR eQTL x* | ***-0.005*** | ***0.021*** | ***-0.004*** | ***0.031*** | -0.002 | 0.149 | 0.000 | 0.960 | 0.801 | 0.169 |
| *age* | ***(-0.008, -0.001)*** | | ***(-0.007, 0)*** | | (-0.006, 0.001) | | (-0.004, 0.005) | | (-1.400, 3.010) | |

CI = confidence interval; eQTL = expression quantitative trait loci; MET = metabolic equivalent of task; mTOR = mammalian target of rapamycin; SD = standard deviation

**Extended Data Table 11. Age-sensitive phenotypes countered or accentuated in mice subjected to intermittent fasting**

The columns "Age", "Diet" and "Interaction" summarize results of two-way ANOVAs/aligned rank transform (summarizing main effects of age, genotype and diet x age interactions, respectively). Findings summarized in the "Intervention effect" column are based on statistical comparison of Cohen's d intervention effect sizes in young vs. old mice. Comparisons, for which a calculation of Cohen's d was not possible, are classified as "not applicable" (N/A).

| Domain | Test/method | Phenotype | Category | Age | Diet | Interaction | Intervention effect |
|---|---|---|---|---|---|---|---|
| Anatomy and physiology | Body weight | Body weight | Opposing aging | ↑ | ↓ | no | equal in young and old |
| Anatomy and physiology | Bone densitometry | Cortical thickness | Promoting aging | ↓ | ↓ | no | equal in young and old |
| Anatomy and physiology | Bone densitometry | Marrow area | Opposing aging | ↓ | ↑ | yes | in young < old |
| Anatomy and physiology | Organ weight | Liver weight | Opposing aging | ↑ | ↓ | no | equal in young and old |
| Anatomy and physiology | Organ weight | Spleen weight | Opposing aging | ↑ | ↓ | no | equal in young and old |
| Cardiovascular health | Echocardiography | Corrected mass of the left ventricle | Opposing aging | ↑ | ↓ | no | equal in young and old |
| Cardiovascular health | Echocardiography | Diastolic septal wall thickness | Opposing aging | ↑ | ↓ | no | equal in young and old |
| Cardiovascular health | Echocardiography | Left ventricular end-diastolic internal diameter | Opposing aging | ↑ | ↓ | no | equal in young and old |
| Cardiovascular health | Electrocardiography | Corrected duration of the QT interval | Promoting aging | ↑ | ↑ | yes | in young < old |
| Cardiovascular health | Electrocardiography | Duration of the QRS interval | Promoting aging | ↑ | ↑ | no | equal in young and old |
| Cardiovascular health | Electrocardiography | Duration of the QT interval | Promoting aging | ↑ | ↑ | yes | in young < old |
| Cardiovascular health | Electrocardiography | Duration of the RR interval | Promoting aging | ↑ | ↑ | yes | in young < old |
| Cardiovascular health | Electrocardiography | Duration of the ST interval | Opposing aging | ↓ | ↑ | yes | equal in young and old |
| Cardiovascular health | Electrocardiography | Heart rate | Promoting aging | ↓ | ↓ | yes | in young < old |
| Clinical chemistry | Clinical chemistry | Insulin | Opposing aging | ↑ | ↓ | yes | in young < old |
| Clinical chemistry | Clinical chemistry | Lactate dehydrogenase | Opposing aging | ↑ | ↓ | no | equal in young and old |
| Clinical chemistry | Clinical chemistry | Unsaturated iron binding capacity | Opposing aging | ↑ | ↓ | no | in young > old |
| Clinical chemistry | Clinical chemistry | Urea | Opposing aging | ↑ | ↓ | yes | in young < old |
| Hematology | Hematology | Hematocrit | Opposing aging | ↓ | ↑ | no | equal in young and old |
| Hematology | Hematology | Hemoglobin concentration | Opposing aging | ↓ | ↑ | no | equal in young and old |
| Hematology | Hematology | Platelet distribution width | Opposing aging | ↑ | ↓ | no | equal in young and old |
| Hematology | Hematology | Platelet large cell ratio | Opposing aging | ↑ | ↓ | no | equal in young and old |
| Hematology | Hematology | Mean platelet volume | Opposing aging | ↑ | ↓ | no | equal in young and old |
| Hematology | Hematology | Red blood cell distribution width | Opposing aging | ↑ | ↓ | no | equal in young and old |
| Immune system | Flow cytometry | CD3$^{+}$CD5$^{+}$ T cells % of all leukocytes | Opposing aging | ↓ | ↑ | no | equal in young and old |
| Immune system | Flow cytometry | CD11b$^{+}$/CD11c$^{+}$Ly6C$^{+}$ % of monocytes | Opposing aging | ↑ | ↓ | yes | in young < old |
| Immune system | Flow cytometry | CD11b$^{+}$/CD11c$^{+}$Ly6C$^{-}$ % of monocytes | Opposing aging | ↑ | ↓ | yes | in young < old |
| Immune system | Flow cytometry | NK1.1$^{+}$/NKp46$^{+}$/CD11b$^{+}$ % of NK cells | Promoting aging | ↓ | ↓ | no | equal in young and old |
| Immune system | Flow cytometry | NK1.1$^{+}$/NKp46$^{+}$/CD11c$^{+}$ % of NK cells | Opposing aging | ↑ | ↓ | yes | equal in young and old |
| Immune system | Flow cytometry | CD3$^{+}$CD5$^{+}$NK1.1$^{+}$/NKp46$^{+}$ NKT cells % of all leukocytes | Opposing aging | ↓ | ↑ | yes | in young < old |
| Metabolism | BCA | Body mass NMR | Opposing aging | ↑ | ↓ | yes | equal in young and old |
| Metabolism | BCA | Fat mass NMR | Opposing aging | ↑ | ↓ | yes | in young < old |

| Domain | Test/method | Phenotype | Category | Age | Diet | Interaction | Intervention effect |
|---|---|---|---|---|---|---|---|
| Metabolism | BCA | Lean mass NMR | Opposing aging | ↑ | ↓ | yes | equal in young and old |
| Metabolism | Indirect calorimetry | Average heat production | Opposing aging | ↑ | ↓ | no | equal in young and old |
| Metabolism | Indirect calorimetry | Average oxygen consumption | Opposing aging | ↑ | ↓ | no | equal in young and old |
| Metabolism | Indirect calorimetry | Average respiratory exchange rate | Opposing aging | ↓ | ↑ | yes | in young < old |
| Metabolism | Indirect calorimetry | Delta respiratory exchange rate | Opposing aging | ↓ | ↑ | no | in young < old |
| Metabolism | Indirect calorimetry | Maximal heat production | Opposing aging | ↑ | ↓ | no | equal in young and old |
| Metabolism | Indirect calorimetry | Maximal oxygen consumption | Opposing aging | ↑ | ↓ | no | equal in young and old |
| Metabolism | Indirect calorimetry | Maximal respiratory exchange rate | Opposing aging | ↓ | ↑ | yes | in young < old |
| Metabolism | Indirect calorimetry | Minimal respiratory exchange rate | Promoting aging | ↓ | ↓ | no | equal in young and old |
| Metabolism | Indirect calorimetry | Water consumption | Opposing aging | ↓ | ↑ | no | equal in young and old |
| Neuropsychiatric functions | ASPPI | Startle response at 110 dB | Promoting aging | ↓ | ↓ | no | in young < old |
| Neuropsychiatric functions | Open field | Average speed | Opposing aging | ↓ | ↑ | no | equal in young and old |
| Neuropsychiatric functions | Open field | Time spent in the center during the first 5 minutes | Opposing aging | ↓ | ↑ | no | equal in young and old |
| Neuropsychiatric functions | Open field | Total distance traveled | Opposing aging | ↓ | ↑ | no | equal in young and old |
| Neuropsychiatric functions | Open field | Total number of rearings | Opposing aging | ↓ | ↑ | no | equal in young and old |
| Neuropsychiatric functions | Open field | Total time spent in the center | Promoting aging | ↑ | ↑ | no | equal in young and old |
| Neuropsychiatric functions | SHIRPA | Locomotor activity | Opposing aging | ↓ | ↑ | no | equal in young and old |
| Neuropsychiatric functions | SHIRPA | Transfer arousal | Opposing aging | ↓ | ↑ | yes | in young > old |
| Neuropsychiatric functions | SHIRPA | Urination | Opposing aging | ↑ | ↓ | no | equal in young and old |
| Neuropsychiatric functions | SHIRPA | Vocalization | N/A | ↓ | N/A | yes | N/A |
| Sensory systems | Hot plate | Time to second response | Promoting aging | ↑ | ↑ | no | equal in young and old |
| Sensory systems | Virtual drum | Spatial frequency threshold | Opposing aging | ↓ | ↑ | yes | equal in young and old |

ASPPI = Acoustic startle and pre-pulse inhibition; BCA = Body composition analysis; CD = Cluster of differentiation; N/A = not applicable; NK = Natural killer cells; NMR = Nuclear magnetic resonance
